# Genome-wide association study of gastrointestinal disorders reinforces the link between the digestive tract and the nervous system

**DOI:** 10.1101/811737

**Authors:** Yeda Wu, Graham K. Murray, Enda M. Byrne, Julia Sidorenko, Peter M. Visscher, Naomi R. Wray

## Abstract

Genetic factors are recognized to contribute to common gastrointestinal (GI) diseases such as gastro-oesophageal reflux disease (GORD), peptic ulcer disease (PUD), irritable bowel syndrome (IBS) and inflammatory bowel disease (IBD). We conducted genome-wide association analyses based on 456,414 individuals and identified 27 independent and significant loci for GORD, PUD and IBS, including SNPs associated with PUD at or near genes *MUC1, FUT2, PSCA* and *CCKBR*, for which there are previously established roles in *Helicobacter pylori* infection, response to counteract infection-related damage, gastric acid secretion and gastrointestinal motility. Post-GWAS analyses implicate putative functional links between the nervous system and gastrointestinal tract for GORD, PUD and IBS, including the central nervous system, the enteric nervous system and their connection. Mendelian Randomisation analyses imply potentially bi-directional causality (the risk of GORD in liability to major depression and the risk of major depression in liability to GORD) or pleiotropic effect between them. A stronger genetic similarity among GORD, PUD and IBS than between these disorders and IBD is reported. These findings advance understanding the role of genetic variants in the etiology of GORD, PUD and IBS and add biological insights into the link between the nervous system and the gastrointestinal tract.

## Introduction

Gastrointestinal (GI) diseases are highly prevalent in western countries. They use substantial health care resources, are a heavy societal economic burden^1,2^, and impact the quality of life of those affected. Common GI disorders include gastro-oesophageal reflux disease (GORD), peptic ulcer disease (PUD), irritable bowel syndrome (IBS) and inflammatory bowel disease (IBD). In GORD, the stomach contents leak back from the stomach into the esophagus^3^. PUD involves breaks (ulcers) in the inner lining of the digestive tract, usually located in the stomach or proximal duodenum. IBS is a chronic functional disorder of the GI system. Patients with IBS often manifest abdominal pain and altered bowel habit, with either predominantly diarrhea, constipation or both. IBD includes Crohn’s disease (CD) and ulcerative colitis (UC), which are chronic idiopathic disorders causing inflammation of the GI tract.

GORD is a multifactorial disorder and is more common in individuals with obesity and hiatal hernia^4^. Lifetime risk estimates of GORD have a wide range (9%-26%), with a sample size-weighted mean of 15%^5^. An increase in the prevalence of GORD since 1995 has been reported^5^. PUD is a complex disorder, for which *Helicobacter pylori* infection and the use of non-steroidal anti-inflammatory drugs (NSAIDs) are the main risk factors^6^. The development of infection-relevant PUD is recognised to be a multistep process, with contributions from both *Helicobacter pylori* infection and subsequent inflammation and damage of mucosa^6^. Eradicating *Helicobacter pylori* is effective for infection-relevant PUD treatment^6^. However, understanding the host factors influencing *Helicobacter pylori* infection and subsequent response could contribute to earlier risk identification and/or prevention, especially given the increasing antimicrobial resistance worldwide^6^. Moreover, clinical presentation of PUD that is not associated with *Helicobacter pylori* infection, nor with the use of NSAIDs, are now also imposing substantial diagnostic and therapeutic challenges^6^. Lifetime prevalence of PUD in the general population has been estimated to be about 5-10%^6^. IBS, a common disorder with a population lifetime risk of 11% globally^7^, is also likely a multifactorial disease, where hypervigilance of the central nervous system, immune activation of the intestinal mucosa, microbiome, prior infections and diet are all suspected to play a role^8^. Similarly, IBD is associated with many dietary and lifestyle risk factors^9^ and lifetime risk for IBD is around 0.3% in most countries of Europe^10^. The genetic contribution to IBD has been well-recognised^11-14^, and well-powered genome-wide association studies (GWASs) have identified >200 approximately independent susceptibility loci associated with IBD^15^. These loci implicate pathways such as autophagy and the IL-17/IL-23 axis and provide insights into IBD pathogenesis^15^. While IBD has been extensively studied through the GWAS paradigm, only a few GWASs for GORD^16^, PUD^17^ and IBS^18-20^ have been conducted to date, most of which were under-powered.

Here, we aim to identify genetic susceptibility factors for GORD, PUD, IBS using the genome-wide association study (GWAS) paradigm. We investigate the shared genetic architecture among these three disorders, and contrast with published GWAS results from IBD. Although a recent study shows that the gut microbiota composition distinguishes IBD from IBS^21^ and although a difference between IBD and IBS from a genetic perspective is expected, it has not yet been quantified. In addition, there is increasing evidence for the importance of bidirectional signalling between the brain and the gut^22-24^, possibly contributing to observational associations between depression and GORD^25^, PUD^26^, IBS^27^ and IBD^28^; however, the potential causal role of depression in each of the four disorders has not yet been established. Here, we also explore the potential causal relationships between major depression (MD) and the four disorders using Mendelian Randomisation (MR), which may help clarify the role of MD in the aetiology of these digestion disorders.

## Results

The workflow for the study is given in **Figure S1**.

### Prevalence and comorbidity for digestion disorders

Based on disease-diagnosis (self-reported or primary/secondary diagnosis in hospital admission records) in the UK Biobank (UKB), four case-control digestion disorder datasets were identified (**Table 1**). GORD is the most common of the GI disorders (8.7%), while prevalence for PUD, IBS and IBD are 2.7%, 3.3% and 1.3%, respectively (**Table 1**). The male/female odds ratio for being PUD case is 1.62 while for IBS it is 0.40 (**Table 1**), consistent with PUD being more common in men and IBS more common in women. The odds of co-occurring diagnosis of a second disorder given diagnosis of a first disorder (**Figure 1A, 1B**) may reflect the natural course of the symptom presentation and/or misdiagnosis. For example, while rates of PUD, IBS or IBD in those diagnosed with GORD were significantly lower than the rate of GORD cases in the UKB as a whole, those with a PUD, IBS or IBD diagnosis were significantly more likely to also have a GORD diagnosis. For each of the GORD, PUD, IBS and IBD (defined as the index disease), competitive comorbidity analyses tested among the other three diseases which disease is more prone to be comorbid with the index disease. We found that PUD is more prone to be comorbid with GORD while IBS is more likely to be comorbid with IBD (**Figure 1C**). In the UKB data there is no information about date of diagnosis for self-reported diseases. Thus, it is not possible to infer a time course in baseline data, and these cases are considered as prevalent cases. There is some information on incident cases that are present in medical records accessed after the baseline visit; however, the sample size is too small to conduct analyses.

**Table 1.** Full-sibling relative risk and heritability estimation for GORD, PUD, IBS and IBD

| Digestion phenotypes | GORD | PUD | IBS | IBD |
| --- | --- | --- | --- | --- |
| N case:N control | 39851:416563 | 12226:444188 | 14994:441420 | 6115:450299 |
| N case/(N case + N control) | 0.087 | 0.027 | 0.033 | 0.013 |
| No. of male case:No. of female case | 18332:21519 | 7025:5201 | 3826:11168 | 2961:3154 |
| No. of male control:No. of female control | 190500:226063 | 201807:242381 | 205006:236414 | 205871:244428 |
| Odds of being male case/Odds of being female case | 0.096/0.095 | 0.035/0.021 | 0.019/0.047 | 0.014/0.013 |
| Male:female odds ratio for being case | 1.01 | 1.62 | 0.40 | 1.11 |
| No. of full-sibling pairs where both proband and full-sibling are cases | 490 | 54 | 72 | 36 |
| No. of full-sibling pairs where only the proband is a case | 3484 | 1086 | 1454 | 526 |
| Full-sibling relative risk (95% confidence interval) | 1.41 (1.30-1.54) | 1.77 (1.36-2.30) | 1.44 (1.15-1.80) | 4.78 (3.48-6.56) |
| Heritability (95% confidence interval) <sup>1</sup> | 0.23 (0.17-0.29) | 0.23 (0.12-0.35) | 0.15 (0.06-0.26) | 0.59 (0.45-0.74) |

<sup>1</sup> The corresponding lower and upper values of 95% confidence interval (CI) for risk in full-sibling were used to calculate the 95% CI for heritability estimation.

**Figure 1.**
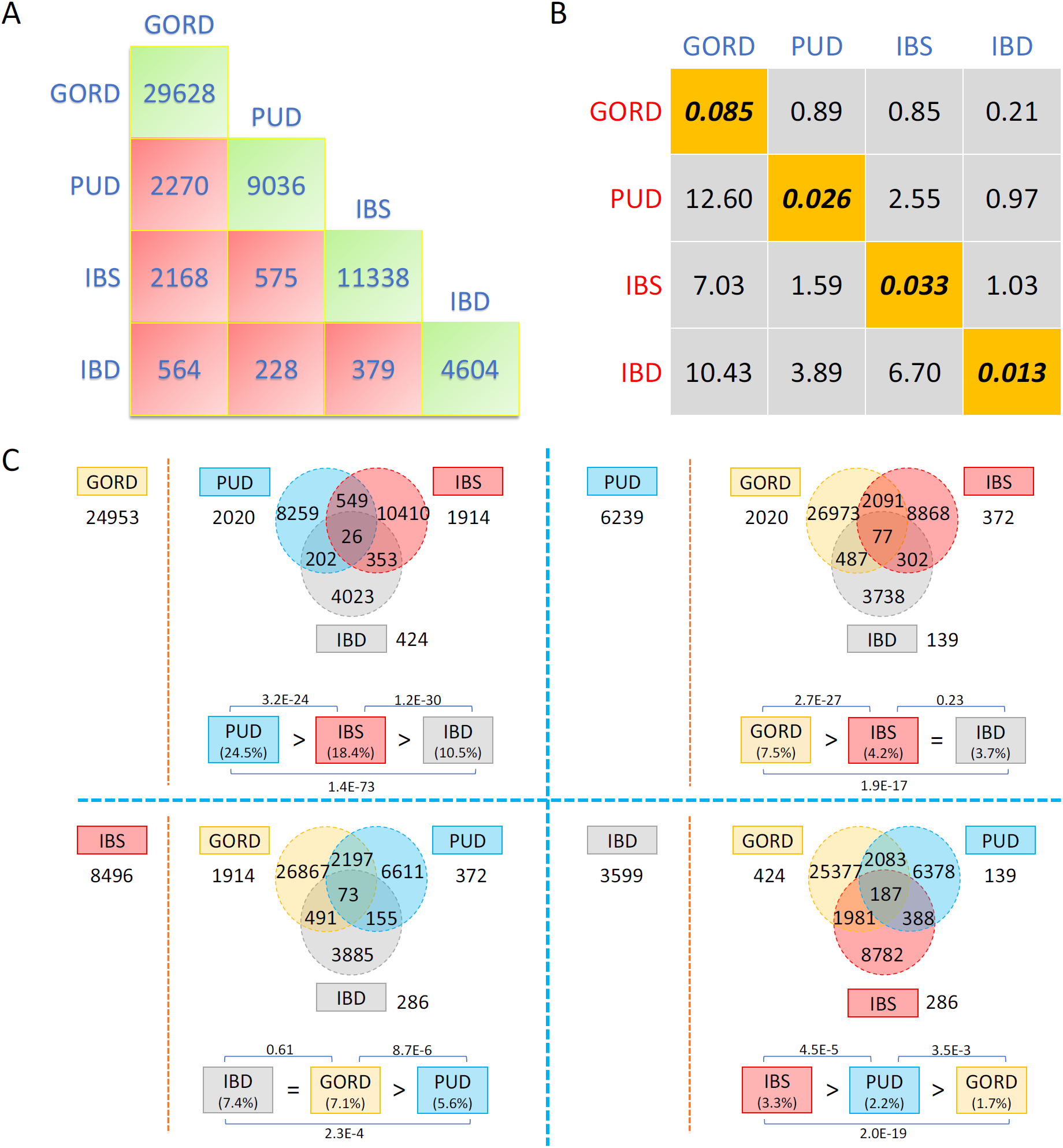
Comorbidity analyses for GORD, PUD, IBS and IBD in unrelated European individuals. **Panel A.** The number of unrelated individuals with each diagnosis (green boxes) and the number of overlapped individuals between each pair of GORD, PUD, IBS and IBD cases (red boxes). **Panel B.** Cells represent ratio of the odds of disease cases from each column in those with disease from each row and the odds of each row disease cases in unrelated European-ancestry individuals. The diagonal elements are the sample risk rates in unrelated European individuals. **Panel C.** For each of GORD, PUD, IBS and IBD disease (defined as index disease), we conduct competitive comorbidity analyses to test among the other three diseases which disease is more prone to be comorbid with the index disease. The disease on the left of red dashed line is the index disease and the number shows the number of cases without any comorbidity. The corresponding Venn diagram shows the number of individuals diagnosed with at least one of the other three diseases. After removing the overlapped individuals for these three diseases, we calculated the number of individuals diagnosed both with the index disease and each of the other three diseases (number outside of the Venn diagram). We then calculated the proportion of the index disease cases in the other three diseases respectively and compared them in pairs by using two-proportion *Z* test to conclude which disease is more prone to be comorbid with the index disease, as shown in the bottom of each Venn diagram (order represents from more prone to less prone).

### Full-sibling risk and heritability estimation

Using inferred coefficients of genetic relationship between individuals in the UKB, we estimated the full-sibling relative risk for each of the GORD, PUD, IBS and IBD and the heritability of liability, with the assumption that the increase risk in relatives only reflect shared genetic factors (**Table 1**). The estimated heritability for GORD, PUD, IBS and IBD were 0.23 (95% CI: 0.17-0.29), 0.23 (95% CI: 0.12-0.35), 0.15 (95% CI: 0.06-0.26), and 0.59 (95% CI: 0.45-0.74), respectively, all significantly different from zero. Sensitivity analyses retained individuals with only one recorded GI disorder and the heritability estimates are similar as above (**Table S1**) and all significantly different from zero.

### GWAS of six digestion phenotypes

Genome-wide association analyses were conducted for six digestion phenotypes, the four disease-diagnosis traits (GORD, PUD, IBS, IBD) and two traits that combined the disease-diagnosis and taking of corresponding medications (i.e. _+_M, **Table S2**). In clinical practice, medications for PUD also have a therapeutic effect on GORD, hence we generated GP_+_M phenotype - a combination of disease-diagnosis of GORD and PUD and corresponding medication-use. We tested for association between 8,546,066 DNA variants and each of the six digestion phenotypes (GORD, PUD, GP_+_M, IBS, IBS_+_M and IBD) in 456,414 UKB participants. A total of 50 independent variants were genome-wide significant (P < 5.0E-8) across the six digestion phenotypes analysed, of which 5 were associated with GORD, 4 with PUD, 0 with IBS, 15 with GP_+_M, 3 with IBS_+_M and 23 with IBD. Given the focus of our study, **Table 2** lists the 27 genome-wide significant SNPs for GORD, PUD, GP_+_M and IBS_+_M and SNPs associated with IBD are in **Table S3**. The GERA cohort data were available as a replication sample for PUD (1,004 cases, 60,843 controls). Of the four genome-wide significant SNPs for PUD in UKB, all have very similar effect size estimates in GERA (**Table S4**) but only rs681343 is formally significant (P = 5.0E-4). Many of the SNPs reported in **Table 2** are novel. Genes around these SNPs have biological support for their mechanistic involvement in corresponding diseases, even known therapeutic-effect mediating target genes for the treatment of corresponding diseases. Notable results are presented in the discussion section. Among 23 IBD-associated SNPs in UKB, 21 have been previously linked with inflammatory bowel diseases (**Table S3**). **Figure 2** shows Manhattan plots for GORD, PUD, GP_+_M and IBS_+_M and **Figure S2** shows Manhattan plots for the other two phenotypes. Quantile-Quantile (Q-Q) plots of all the variants analysed in UKB are provided in **Figure S3** for the six phenotypes. Regional visualisation plots of the 50 independent variants are in **Supplementary Data 1**. Detailed pleiotropy results derived from GWAS Catalog^29^ are provided in **Supplementary Data 2**.

**Table 2.**
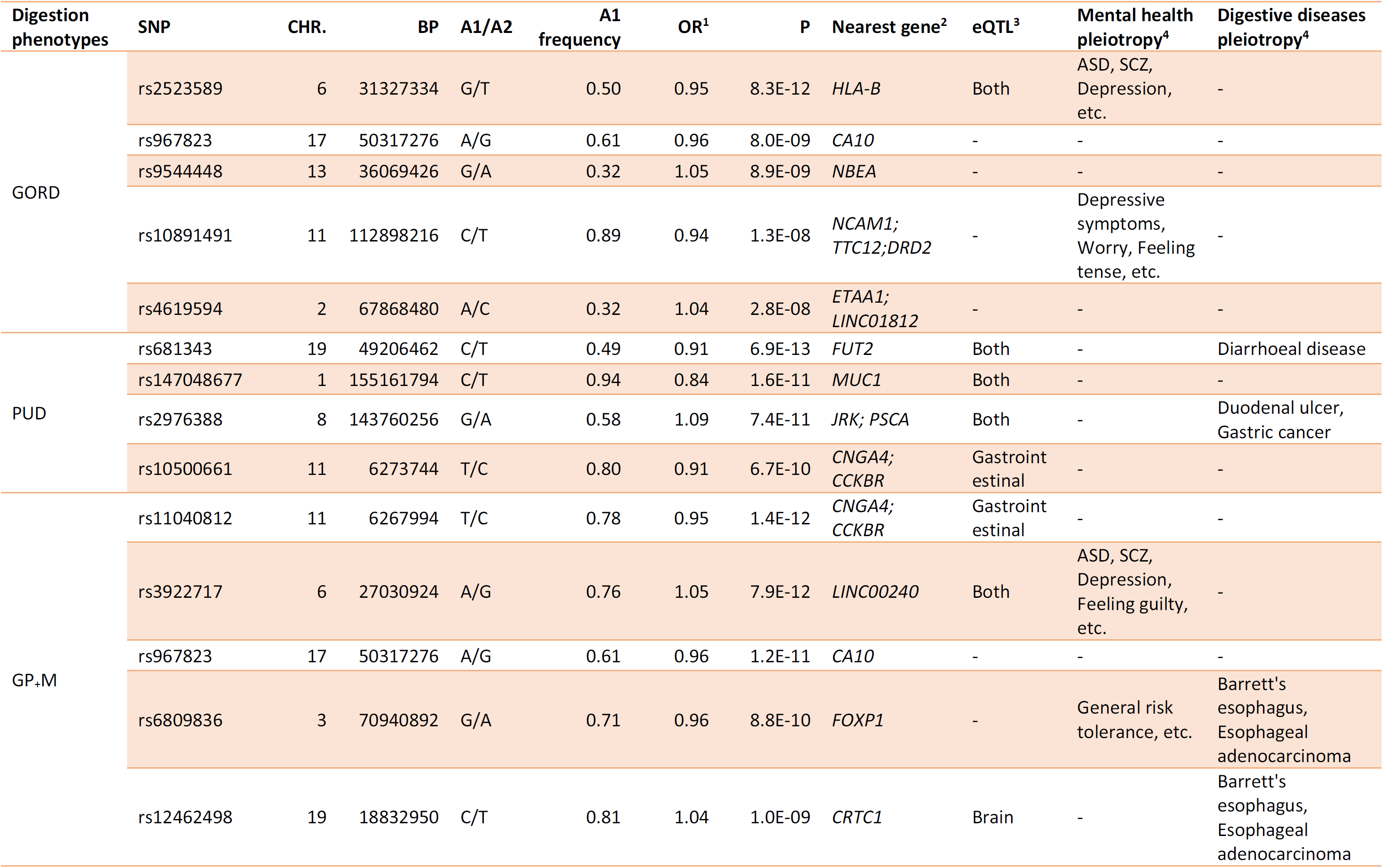
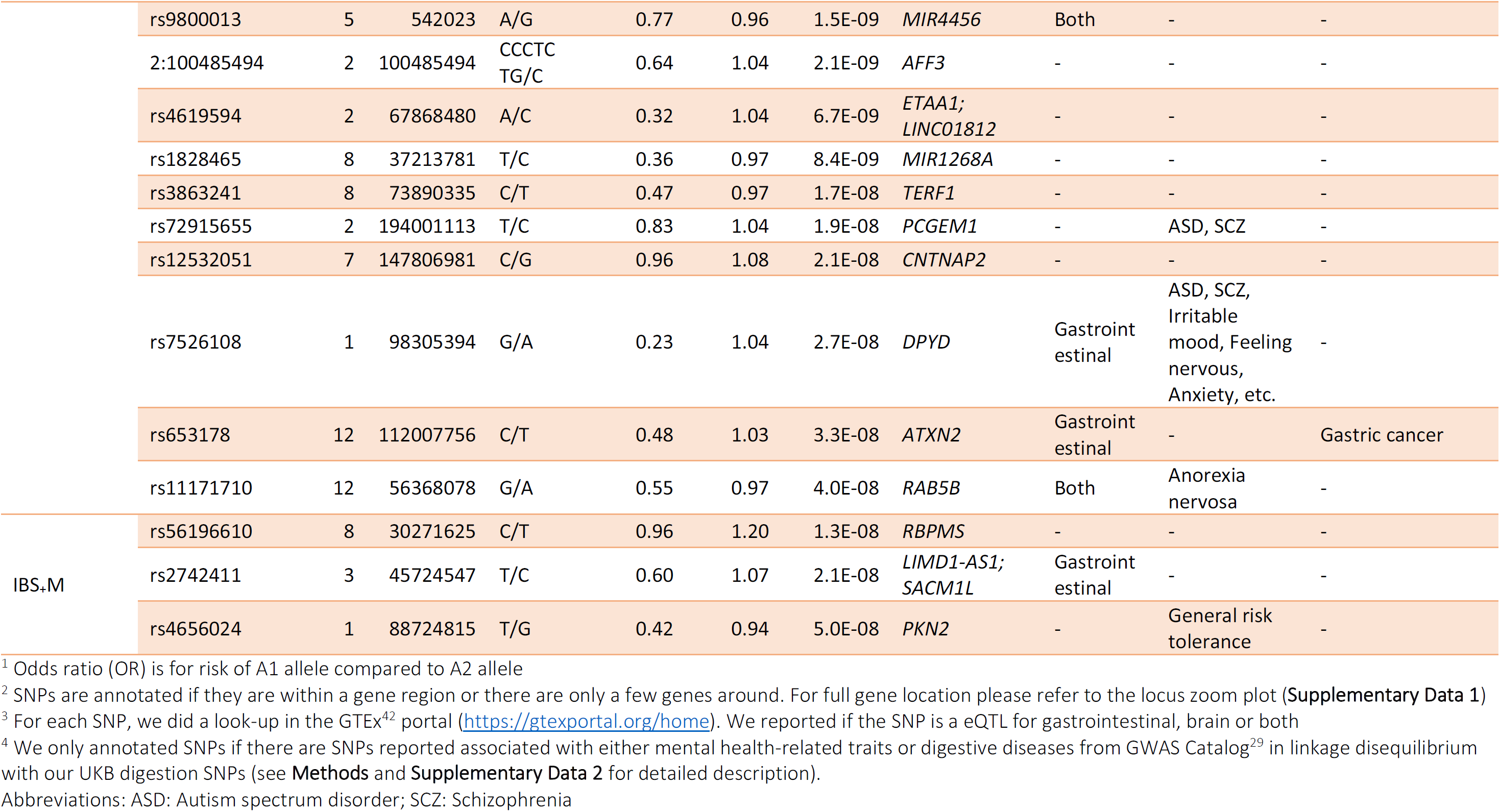
Genome-wide significant SNPs associated with GORD, PUD, GP_+_M and IBS_+_M in the UK Biobank

**Figure 2.**
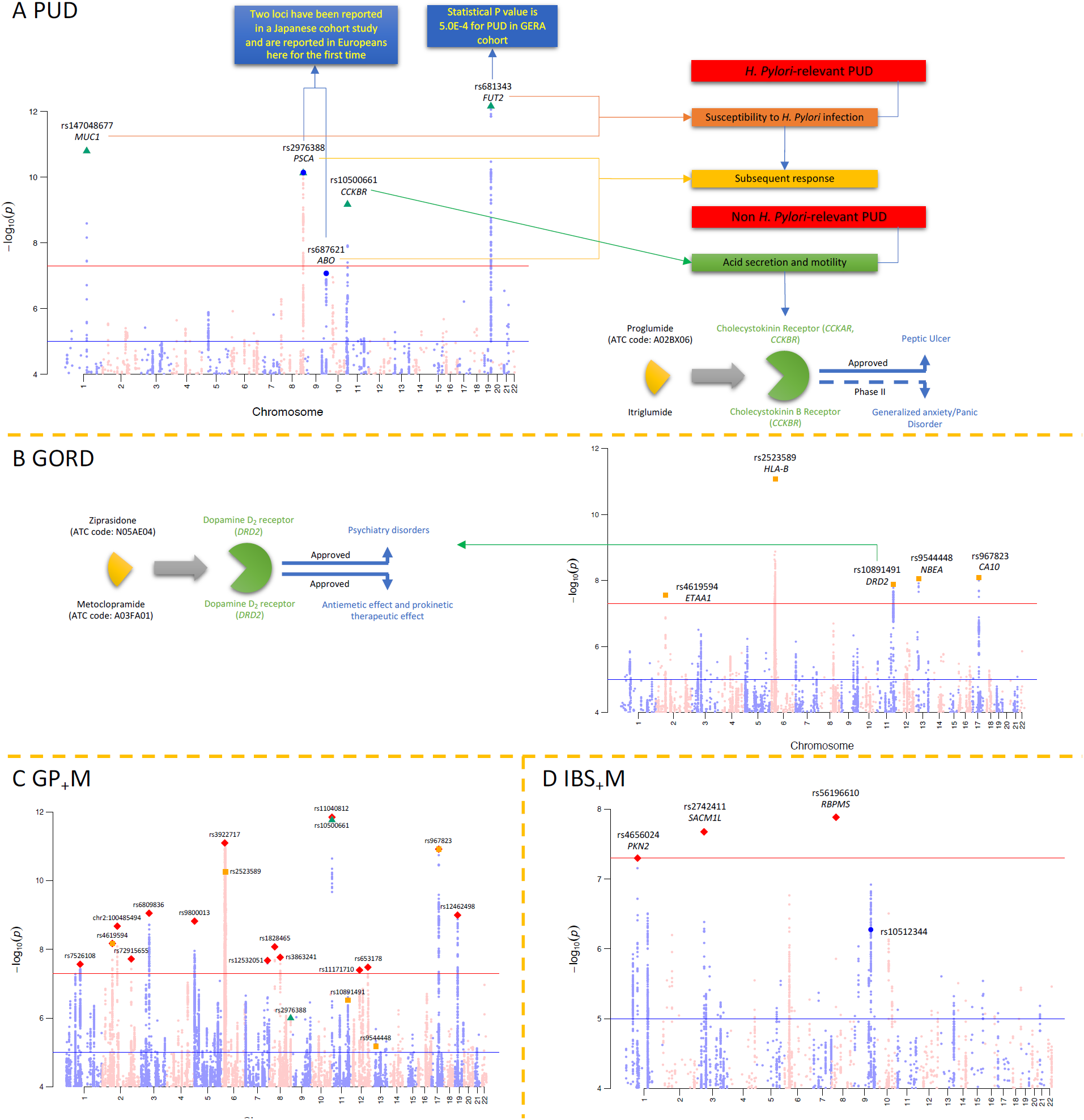
Manhattan plots for GORD, PUD, GP_+_M and IBS_+_M for SNPs associated P < 1.0E-5. **Panel A**. Association with PUD. SNPs highlighted with green triangles are independent loci with P < 5.0E-8, corresponding to the green triangles in Panel C. The blue dots are for SNPs rs2976388 and rs687621, the only two loci associated with duodenal ulcer in a Japanese cohort^17^. rs681343 showed statistically significant in GERA PUD GWAS summary statistics, as annotated in the blue box. Schematic diagram on the right side represents the reported biological evidence supporting genes around the PUD associated loci in peptic ulcer involvement. *MUC1* and *FUT2* have been linked to susceptibility to *Helicobacter Pylori* infection and *PSCA* and *ABO* have been proposed to be associated with subsequent response after infection. *CCKBR* encodes cholecystokinin receptor mediating therapeutic effect for peptic ulcer treatment by reducing acid secretion and inhibiting gastrointestinal motility. The cholecystokinin receptor is also an effect-mediating target of itriglumide on phase II clinical trial for anxiety and panic disorder. **Panel B**. Association with GORD. SNPs highlighted with orange squares are genome-wide statistically significant (P < 5.0E-8) independent loci, which correspond to the orange squares in Panel C. *DRD2* is near GORD-associated SNP rs10891491and encodes dopamine D_2_ receptor, which is a target for psychiatry disorders and blocking this receptors in the chemoreceptor trigger zone relieves nausea and vomiting feeling and in the gastrointestinal tract increases motility^61^, as shown in the left side of Manhattan plot for GORD. **Panel C**. Association with GP_+_M. SNPs highlighted with red diamond are independent loci with P < 5.0E-8. All the SNPs associated with GORD (highlighted with orange squares) have P < 1.0E-5 in GP_+_M and only 2 of 4 SNPs associated with PUD (highlighted with green triangles) are with P < 1.0E-5 in GP_+_M. rs2976388 is also highlighted with blue dots given the previous reported association with duodenal ulcer in a Japanese cohort. **Panel D**. Association with IBS_+_M. SNPs highlighted with red diamond are independent loci with P < 5.0E-8. The blue dot is for rs10512344 that has been reported associated with female IBS in the UKB data^20^.

### SNP-based heritability and genetic correlation of the six digestion phenotypes

LDSC^30^ SNP-based heritability 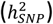 estimates on the liability scale were all significantly different from zero: GORD 0.08 (SE = 0.005), PUD 0.05 (SE = 0.008), GP_+_M 0.10 (SE = 0.004), IBS 0.06 (SE = 0.008), IBS_+_M 0.07 (SE = 0.008) and IBD 0.12 (SE = 0.017) (**Figure 3A, Table S5**). The SNP-based genetic correlations (*r*_*g*_) between GORD and PUD is 0.65 (SE = 0.06, P_H0:rg=0_ = 6.5E-28, phenotypic correlation (*r*_*p*_) = 0.10), similar to the *r*_*g*_ estimate for GORD and IBS (0.61, SE = 0.06, P_H0:rg=0_ = 1.5E-26, *r*_*p*_ = 0.08). The *r*_*g*_ between PUD and IBS is 0.48 (SE = 0.10, P_H0:rg=0_ = 8.0E-7, *r*_*p*_ = 0.03) (**Table S6**), while the *r*_*g*_ between IBD and each of GORD, PUD, GP_+_M, IBS and IBD are not statistically significantly different from zero after Bonferroni correction (**Figure 3B**). In sensitivity analyses, all individuals with more than one GI diagnosis were excluded (**Figure S4A**). As expected, the 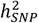 estimates were lower but still significantly different from zero (**Figure S4B**). GORD, PUD and IBS are significantly genetically correlated while none of them showed statistically significant *r*_*g*_ with IBD (**Figure S4C**). Detailed results of sensitivity analyses are discussed in **Supplementary Note 1** with the corresponding data presented in **Table S7** and **Figure S4**.

**Figure 3.**
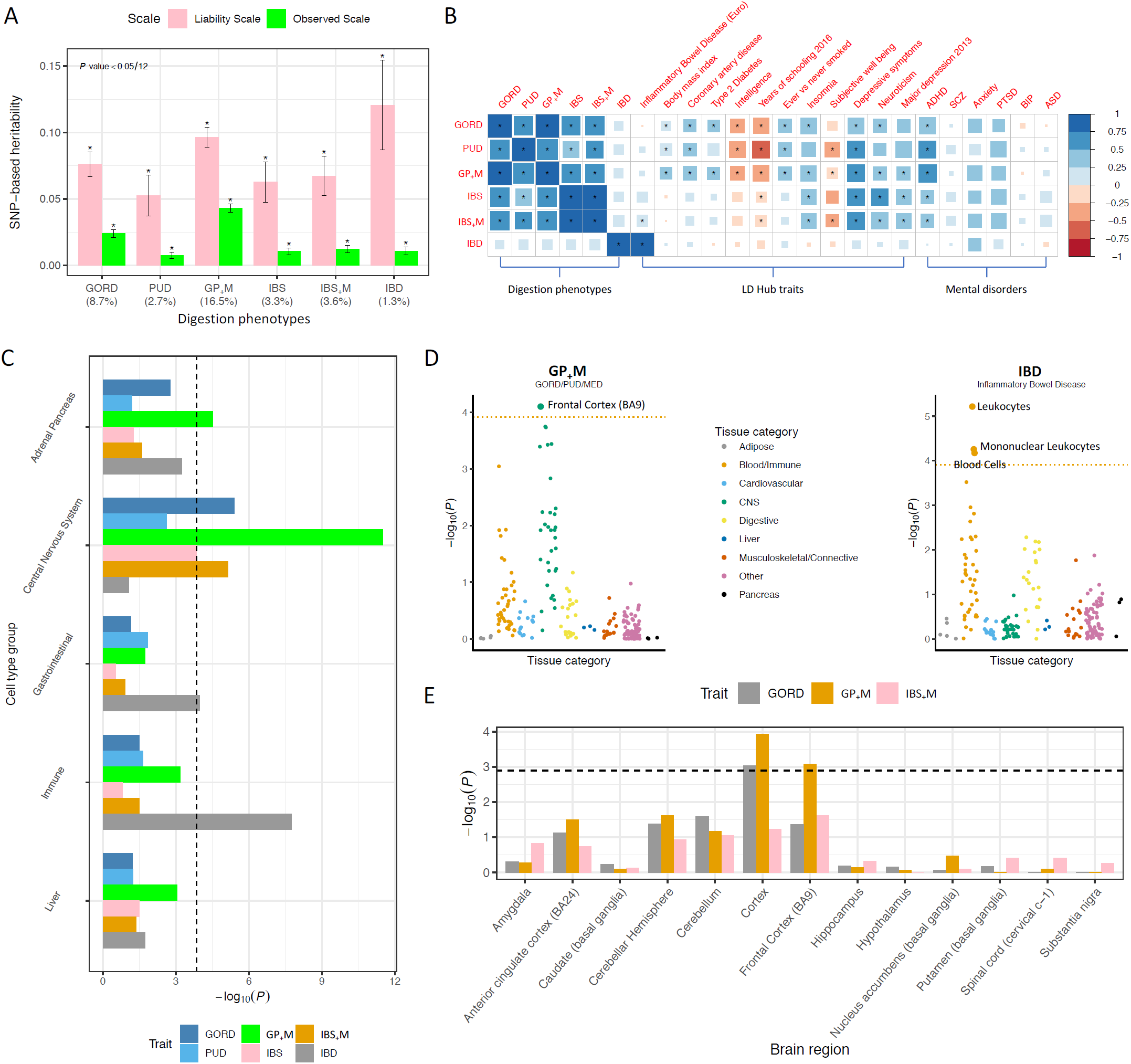
LD score regression SNP-based heritability and genetic correlation analyses for the six digestion phenotypes. **Panel A**. SNP-based heritability of the six digestion phenotypes both on the observed and liability scales. We took sample risk, i.e. the proportion cases in the UKB cohort, as the population lifetime risk to calculate the SNP-based heritability on the liability scale for each digestion phenotype; the sample risk percentage is shown below x axis in parentheses. **Panel B**. Genetic correlation within digestion phenotypes, between each of the digestion phenotypes with traits from LD Hub and six published mental disorder studies. “*” represent that genetic correlation estimates are still significant after Bonferroni correction (P < 0.05/(6*6+252*6+6*6)). **Panel C**. SNP-based heritability enrichment analysis of each digestion phenotypes partitioned by cell type groups annotated by histone marks. The dashed line represents the Bonferroni correction threshold (0.05/(53*6+5*6)). **Panel D**. SNP-based heritability enrichment analysis for GP_+_M and IBD partitioned by cell types annotated using cell-type specific gene expression data. The dotted lines represent the Bonferroni correction threshold (P < 0.05/(205*2)). **Panel E**. SNP-based heritability enrichment analysis for GORD, GP_+_M and IBS_+_M partitioned by cell types annotated using GTEx brain gene expression data from 13 brain regions. The dashed line represents the Bonferroni correction threshold (P < 0.05/(13*3)).

The *r*_*g*_ between each of the six phenotypes and six published psychiatric traits^31-36^ and 252 other traits from LD Hub^37^ (**Table S8** and **Supplementary Data 3**, respectively) included 27, 20, 45, 11, 14 and 3 significant correlations for GORD, PUD, GP_+_M, IBS, IBS_+_M and IBD, respectively, after Bonferroni correction (P < 3.2E-5). **Figure 3B** shows the *r*_*g*_ between each of the six phenotypes and selected traits from the 258 traits. Interestingly, we observed significant *r*_*g*_ between three digestion phenotypes and depressive symptoms^38^ (GORD (0.46, SE = 0.05, P_H0:rg=0_ = 1.7E-21), PUD (0.52, SE = 0.09, P_H0:rg=0_ = 1.2E-9), IBS (0.52, SE = 0.07, P_H0:rg=0_ = 1.5E-12)). IBS was significantly genetically correlated with major depression (MD)^39^ (0.43, SE = 0.10, P_H0:rg=0_ = 2.1E-5)). We also observed significant *r*_*g*_ between IBS and neuroticism^40^ (0.50, SE = 0.06, P_H0:rg=0_ = 1.1E-18), and between GORD and neuroticism^40^ (0.35, SE = 0.04, P_H0:rg=0_ = 3.6E-15). Attention deficit hyperactivity disorder (ADHD)^31^ showed significant *r*_*g*_ with GORD (0.49, SE = 0.04, P_H0:rg=0_ = 6.7E-32), PUD (0.54, SE = 0.07, P_H0:rg=0_ = 9.0E-16) and IBS (0.32, SE = 0.06, P_H0:rg=0_ = 5.3E-7). However, there was no statistically significant *r*_*g*_ between IBD and depressive symptoms or MD. In sensitivity analyses, all individuals with more than one GI diagnosis were excluded and the results are similar as above. Detailed results of sensitivity analysis are discussed in **Supplementary Note 1** with the corresponding data presented in **Table S9, Figure S4** and **Supplementary Data 4**.

### Linking GWAS findings to gene expression

Functional annotation SNP-based heritability analyses estimate enrichment of 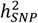 based on SNP annotations compared to the expectation assuming equal partitioning of 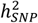 across the genome. After Bonferroni correction, GORD, GP_+_M and IBS_+_M showed significant enrichment of 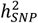 in conserved regions while 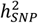 enrichment for IBD was in the super enhancer category (**Figure S5** and **Table S10**). In analyses based on SNP annotations derived from cell-type histone mark data (**Figure 3C** and **Table S11**), IBD showed significant 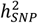 enrichment in immune and gastrointestinal cell-type groups, while GORD, GP_+_M and IBS_+_M showed enrichment in the CNS cell-type. Based on cell-type specific SNP annotations^41^ derived from gene expression data of 205 different tissues (53 from GTEx^42^ and 152 from Franke lab^43,44^), GP_+_M showed significantly enriched association with genes expressed in the frontal cortex of the brain (Brodmann Area, BA9) and IBD showed enriched associations in leukocytes (**Figure 3D** and **Table S12**). In addition, for GORD, GP_+_M and IBS_+_M we conducted the same analysis using the GTEx brain gene expression data which includes data from 13 brain regions^42^. GWAS associations for GP_+_M were consistently enriched in the frontal cortex (BA9) (**Figure 3E** and **Table S13**). We also investigated whether associations between SNPs and the six digestion phenotypes were consistent with mediation through gene expression using the Mendelian randomisation (MR) method, SMR^45^. A total of 9 unique genes for which expression is significantly associated with 3 digestion phenotypes, including 3 genes for PUD, 1 gene for GP_+_M, and 5 genes for IBD, were identified (**Table S14**). Comment on notable results is given in the discussion section.

### Gene-based and gene-set enrichment analyses

We used MAGMA^46^ software to identify genes significantly (P < 2.7E-6) associated with each of the six digestion phenotypes. We identified genes significantly associated with the six digestion phenotypes: 54 for GORD, 18 for PUD, 138 for GP_+_M, 24 for IBS, 25 for IBS_+_M and 96 for IBD (**Supplementary Data 5**). For gene-set enrichment analysis, gene-based summary statistics of GP_+_M showed enrichment in “GO (gene ontology):NEURON PROJECTION MORPHOGENESIS”, “GO:NEUROGENESIS”, “GO:NEURON PROJECTION GUIDANCE” and “GO:ANTIGEN PROCESSING AND PRESENTATION OF ENDOGENOUS ANTIGEN” gene sets. The top enriched gene sets for IBD is “GO RESPONSE TO INTERFERON GAMMA” (**Supplementary Data 6**).

### Comorbidity with depression and Mendelian Randomisation (MR)

Following Cai *et al.*^47^, we derived eight depression phenotypes from UKB (**Methods**) and tested whether the number of individuals who are cases for both a digestion phenotype and depression phenotype is statistically significantly different from the expected number (**Figure S6)**. All eight depression phenotypes showed statistically significant comorbidity relationship with each of GORD, PUD, and IBS. For IBD, only ICD10 defined depression (ICD10Dep), DSM-V clinical guideline defined major depression (LifetimeMDD) and major depression recurrence (MDDRecur) showed statistical significance (**Figure S6)**. We then used Generalized Summary-data-based MR (GSMR)^48^ to test for putative causal association between MD and each of the six digestion phenotypes in UKB, and we also examined reverse causality (**Table S15, Figure S7, Figure S8**). The genetic instruments for MD were from cohorts reported in Wray *et al.*^49^ but from a meta-analyses that excluded the UKB. We observed bidirectional statistically significant results between MD and GP_+_M, i.e. 1.26-fold increased risk for GP_+_M per standard deviation (SD) in liability to MD (P = 1.1E-12), and 1.16-fold increased risk for MD per SD in liability to GP_+_M (P = 8.0E-05). No SNPs were identified as outliers by the HEIDI test. The pattern of results was the same when other MR methods were applied, which as expected showed less significant results (see **Supplementary Note 2, Table S16** and **Figure S9**). For the relationship between MD and IBD, GSMR estimates were not statistically significant either in forward direction (the effect of MD on IBD) or the reverse direction (the effect of IBD on MD). Last, the effect of MD on GORD, PUD, IBS and IBS_+_M showed statistically significant estimates of 1.25-fold, 1.20-fold, 1.36-fold, and 1.36-fold respectively increase per standard deviation (SD) in liability to MD (**Figure 4A**). The point estimates for the reverse causality analyses were smaller, but it is not possible to make strong statements about the significance of the estimates because we needed to relax the significance threshold imposed to achieve a sufficient number of SNP instruments; these analyses should be revisited in when more genome-wide significant SNPs are identified. (**Figure 4A, Table S15**).

**Figure 4A.**
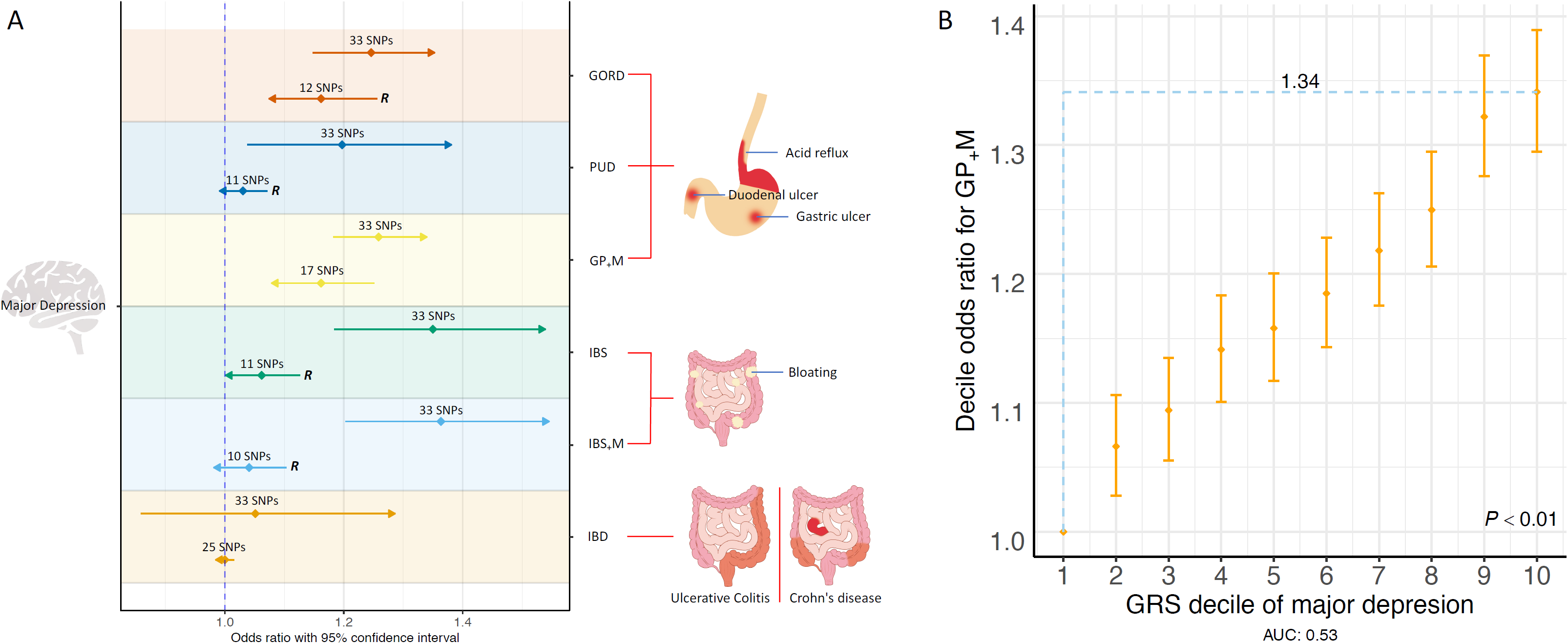
Mendelian Randomisation (MR) results between major depression (MD) and six digestion phenotypes. The left y axis represents MD while the right y axis represents six digestion phenotypes. The arrow for each horizontal line represents the direction from exposure trait to outcome trait relative to the y-axes labels. OR and 95% CI are represented as diamond and horizontal lines taking values from x axis. Each digestive phenotype corresponds to two horizontal lines. “***R***” on the right side of the horizontal line represents relaxation of significance threshold to obtain more genetic instrument (**Table S15**) and the number of the SNP instruments used in analysis are shown above the diamond. The common pathological characteristics or symptoms for these phenotype-related diseases are shown on the right side (but pathological characteristics or symptoms are not limited to these locations). **Figure 4B.** Genetic risk score (GRS) of major depression (MD) predicts odds ratio (OR) for GP_+_M. GRS from MD associated SNPs with P< 0.01 were converted to deciles (1 = lowest, 10 = highest). OR and 95% confidence intervals (CI, orange diamonds and bars) relative to decile 1 were estimated using logistic regression. The blue dashed lines represent that compared with the lowest decile, the highest decile have an OR of 1.34 for GP_+_M related disorders, mainly gastro-oesophageal reflux disease.

### Genetic risk score (GRS) prediction

Given the bidirectional statistically significant results between MD and GP_+_M, we used MD GWAS summary statistics (European ancestry, excluding the UKB cohort)^49^ to generate MD genetic risk scores and used these to predict GP_+_M risk (risk for GORD, PUD and likelihood for taking GORD, PUD medications) in the UKB. Participants in the UKB with a high genetic risk score for MD have a higher risk for GP_+_M-related disorders. The top decile of individuals ranked on genetic risk prediction for MD had an OR of 1.34 (95% CI: 1.29-1.39) for GP_+_M risk compared to the bottom decile (**Figure 4B** and **Table S17**). We also selected genome-wide significant SNPs associated with PUD in UKB to calculate GRS and predict peptic ulcer risk in GERA cohort. The top decile of individuals ranked on genetic risk prediction for PUD had an OR of 1.49 (95% CI: 1.16-1.92) for PUD risk compared to the bottom decile (**Figure S10**).

## Discussion

This study describes an analysis of four common digestion disorders using a single study cohort. We used the both the phenotypes and genotypes of up to 456,414 individuals to study the genetic contributions to GORD, PUD, IBS and IBD and the connection between these disorders with major depression. Our results give evidence for the following conclusions.

First, there is an increased risk for each disorder in full-siblings which suggests a familial contribution. If interpreted as only reflecting shared genetic factors, these generate estimates of heritability (**Table 1**): GORD (0.23), PUD (0.23), IBS (0.15) and IBD (0.59). Many studies report estimates of heritability for IBD using twin and family data, while GORD, PUD and IBS have been less studied. The reported heritability estimates for GORD (0.43^11^ and 0.31^50^), PUD (0.39^51^), IBS (0.48 for female^52^), CD (∼0.7-0.8)^14^ and UC (∼0.6-0.7)^14^ are still higher than our estimates, which could reflect different ascertainment biases in participant recruitment of UKB versus traditional genetic epidemiology studies.

Second, GWASs of GORD, PUD and IBS, together with medication derived related phenotypes, identified 27 quasi-independent loci. Some have not been reported by previous GWAS studies but have biological support for their mechanistic involvement. A Japanese cohort study^17^ identified only two SNPs associated with duodenal ulcer, rs2294008 and rs505922. In our PUD-associated SNPs, rs2976388, located in *PSCA* gene, is in high LD (*r*^2^ = 0.94) with rs2294008, while rs687621(P = 8.7E-8) is in high LD (*r*^2^ = 0.98) with rs505922 (blue dots highlighted in **Figure 2A**). These two loci are reported in Europeans here for the first time. These two loci are likely to be associated with duodenal ulcer development after *Helicobacter pylori* infection^17^. From published data, we also found that allele A of SNP rs2976388 is associated with increased *PSCA* expression (b_eQTL_ = 0.73, P_eQTL_ = 8.8E-41), and through SMR analysis, the expression of *PSCA* decreased risk for PUD (b_SMR_= −0.12, P_SMR_ = 4.8E-9). Decreased *PSCA* expression has been reported following *Helicobacter pylori* infection^53^, indicating negative regulation of *PSCA* expression by *Helicobacter pylori* infection. Other data sets recorded for *Helicobacter pylori* infection status are needed to explore this proposed relationship. Two other novel PUD-associated SNPs may also relate to genetic risk to *Helicobacter pylori* infection: rs147048677 (P = 1.6E-11) is a synonymous variant in the *MUC1* gene. A mouse model study^54^ has shown that Muc1 limits *Helicobacter pylori* colonization of gastric mucosa. rs681343, which we found to be statistically significant in both UKB discovery and GERA replication GWAS for PUD, is located in the *FUT2* gene and this gene has also been implicated in susceptibility to *Helicobacter pylori* infection^55^ in humans. A novel PUD-associated SNP, rs10500661 (P= 6.7E-10) is located in 7.2kb upstream of *CCKBR* (cholecystokinin B receptor) and this gene encodes a G-protein coupled receptor for both gastrin and cholecystokinin, regulatory peptides of the brain and gastrointestinal tract^56^. This gene, as shown in the GTEx^42^ portal (https://gtexportal.org/home/gene/CCKBR), is highly expressed in the brain frontal cortex (Brodmann Area (BA) 9) and stomach. Moreover, this gene is a therapeutic-effect target gene for proglumide (ATC code: A02BX06) to treat peptic ulcer, of which the mechanism is to inhibit gastrointestinal motility and reduce gastric acid secretions. In addition to these findings, itriglumide, an antagonist for the *CCKBR* protein, has been investigated as a potential treatment for anxiety and panic disorders^57^.

For GP_+_M, two of the significantly associated SNPs, rs6809836 and rs12462498, have been previously linked to Barrett’s esophagus and esophageal adenocarcinoma^58,59^ (**Table 2**). Another GORD-associated SNP, rs10891491, is located in the *NCAM1, TTC12* and *DRD2* gene region which has been linked to depressive symptoms. *DRD2* is therapeutic target gene for atypical antipsychotics, such as ziprasidone (ATC code: N05AE04)^60^. Blocking dopamine D2 receptors encoded by this gene in the chemoreceptor trigger zone relieves nausea and in the gastrointestinal tract increases motility^61^. A previous study reported that rs10512344 is the only one SNP genome-wide significantly associated (P = 3.6E-8) with IBS using UKB data^20^. Both the IBS (lead SNP: rs112243849, P = 7.5E-8) and IBS^+^M (lead SNP: rs7861675, P = 1.2E-7) phenotypes in our study reconfirmed the association between this locus and IBS at the genome-wide suggestive level. A detailed comparison with this study is provided in **Supplementary Note 3**.

Third, we provide direct genetic evidence that IBD is etiologically different to the other digestion phenotypes as illustrated by high genetic correlations among GORD, PUD, IBS, which all show low genetic correlations with IBD (**Figure 3B**). Both GORD and PUD are acid-related diseases; their high genetic correlation (*r*_*g*_ = 0.65, P_H0:rg=0_ = 6.5E-28) motivated the combination of GORD and PUD with medication taking cases. The high genetic relationship between GORD and IBS is not unexpected given that a highly comorbid relationship has been previously reported^62^. Orthogonal evidence for genetic differences between IBD and the other digestion phenotypes was provided by the partitioned SNP-based heritability analyses, which showed enrichment of GP_+_M-associated SNPs in genes expressed in the BA9 region of the brain cortex while those for IBD are enriched in blood and immune related tissues. Gene set enrichment analysis showed GP_+_M associated SNPs are enriched in neuron-related gene sets. We note that a limitation of our brain enrichment analysis is that our conclusions are limited by the availability of tissue specific gene expression data. The GTEx database does not report gene expression data for multiple cortical regions, so specificity to BA9 or frontal cortex (over and above other cortical regions) is not established. As discussed by Finucane *et al.*^41^, it is not possible to draw strong conclusions about the most likely causal tissue or cell type as only a subset of cell types are tested, we can only say tissues with similar gene expression profiles to BA9 of brain or BA9 itself may be relevant to GP_+_M. Given the non-availability of gene expression data from other human tissues, such as sympathetic, parasympathetic (vagus nerve)^23^ and enteric nervous system^22,24^, we cannot conduct key hypothesis based enrichment analyses. However, despite these limitations, our findings indicate that a genetic contribution to GP_+_M may highlight the potential link between the nervous system and oesophagus, stomach and duodenum though there is likely not just one causal tissue or cell type. Historically, vagotomy was used commonly to manage peptic ulcer diseases, as vagal stimulation promotes acid secretion^63^ (now successfully treated by H2 receptor agonists), indicating the clinical importance of the link between the nervous system and gastrointestinal tract. We note that we did observe a significant genetic correlation between IBS_+_M and published IBD European GWAS summary statistics^64^ (*r*_*g*_ = 0.23, SE = 0.05, P_H0:rg=0_ = 1.6E-5, the *r*_*g*_ between IBS and IBD was 0.21, SE = 0.05, P_H0:rg=0_ = 2.0E-4). However, this relative lower genetic correlation suggests that IBD is etiologically different from IBS.

Fourth, we conducted Mendelian Randomisation (MR) to investigate if this methodology provides evidence of a causal relationship between major depression and GI disorder phenotypes. The association between mental health and GORD has been addressed through observational studies^25,65^. For example^65^, a bidirectional association between GORD and depression was reported with risk factor roles for depression on GORD and for GORD on depression. The MR results using GORD, PUD and GP_+_M are qualitatively similar and so for discussion purposes we focus on the combined GP_+_M phenotype that identified a higher number genome-wide significant SNPs for use in the MR instruments. We found an OR of 1.26 (P = 1.1E-12) for GP_+_M per SD in liability to MD, which has direction and effect size estimates consistent with those previously reported between MD and drugs for GORD and PUD (OR: 1.23, P = 4.0E-6)^66^. However, the reverse MR analysis (the effect of GP_+_M on MD) is also significant (OR: 1.16, P = 8.0E-5). The bidirectional statistically significant results from GSMR usually include two interpretations, one is that there is the bidirectional causality and the other one is horizontal pleiotropy, including an indirect relationship through an intermediate endophenotype^67^. Hence, our conclusions must include these interpretations although we note that MR Egger intercept test suggest no horizontal pleiotropy. In terms of bidirectional causality, there are several possible explanations. First, there are the intrinsic links between major depression and GORD. Patients with psychological comorbidity often perceive low intensity oesophageal stimulation as being painful due to hypervigilance to these intra-oesophageal events^68^. Psychological factors can decrease the pressure of the lower oesophageal sphincter and change oesophageal motility^69^. The reflux symptom itself could result in depression if patients are constantly feeling upset about their condition^69^. Second, the use of medications could mediate the effect. Tricyclic antidepressants can lead to a decrease in lower oesophageal sphincter pressure and thus and increase in the number of reflux episodes (anticholinergic effect)^70^. Recently a study shows that use of proton-pump inhibitors (PPI) for acid-related disorders are associated with the subsequent risk of major depression disorder^71^. Further studies are needed to clarify this association. The connection between major depression and GORD is complex, potentially involving the interplay of multiple mechanisms, further studies from multi-disciplines are needed to understand the connection. While a causal relationship cannot be confirmed between major depression and digestion related disorders, consideration of clinical implications of a possible relationship is justified. When treating patients with MD, awareness of the digestion symptoms for GORD could help to decide if further interventions are needed. Also, these results may provide clues for screening psychological factors in GORD patients. Previous study^72^ shows that GORD patients who are also comorbid with psychologic distress are associated with more severe symptoms at baseline and more residual symptoms after PPI treatment. In those patients, treatment for the underlying psychological distress might improve the PPI response^73^.

Despite these interesting findings, our study has several limitations. First, the phenotype of GORD, PUD and IBS was a combination of self-reported illness, medication-use and clinical diagnosis record. There is a potential influence regarding self-reported accuracy and misdiagnosis. Given the existent co-reporting of some diagnoses, we conducted sensitivity analyses in which individuals recorded with more than one diagnosis were excluded, but these analyses did not impact our conclusions. Importantly, we note that the GWAS results from IBD derived from the UKB were highly consistent with results from published GWAS. Second, our study is conducted in the UK Biobank cohort, which while a large population study has recognised volunteer bias^74^. Third, we do not have replication data sets for GORD, GP_+_M and IBS_+_M genome-wide significant SNPs. Fourth, we do not have the *Helicobacter pylori* infection status and microbiome data in UKB, thus additional analyses on these factors cannot be further explored. Fifth, we note a recent a meta-analysis GWAS study for GORD^75^ using UKB cohort, however, there are differences with our study. Our study focus is on GWAS of multiple digestion disorders, not limited to GORD. Our study addresses the link between the digestive tract and nervous system while they focus on GORD-associated loci that also associated with Barrett’s oesophagus and oesophageal adenocarcinoma. We also explore the comorbidity relationship within digestion disorders and between each of digestion disorders with depression in UKB.

In summary, the study identified 27, mostly novel, independent SNPs associated with different digestion disorders, including SNPs at or near *MUC1, FUT2, PSCA* and *CCKBR* genes associated with peptic ulcer disease, for which previously established roles of these genes in *Helicobacter pylori* infection, response to counteract infection-related damage and gastric secretion support their involvement. Post-GWAS analyses highlighted the link between nervous system and the gastrointestinal tract, which may add biological insight into nervous system-gastrointestinal tract links and aetiology of relevant diseases. In addition to this, Mendelian Randomisation analyses imply potentially bi-directional causality (the risk of GORD in liability to major depression and the risk of major depression in liability to GORD) or pleiotropic effect between them. Taken together, our findings demonstrate the role of genetic variants in the aetiology of common digestion disorders and the link between depression and GORD.

## URLs

UK Biobank: https://www.ukbiobank.ac.uk/about-biobank-uk/; dbGaP: https://www.ncbi.nlm.nih.gov/gap/; LD Score Regression: https://github.com/bulik/ldsc/; LD Hub: http://ldsc.broadinstitute.org/ldhub/; GTEx: https://gtexportal.org/home/; SMR: https://cnsgenomics.com/software/smr/#Overview/; MAGMA: https://ctg.cncr.nl/software/magma/; GSMR: http://cnsgenomics.com/software/gcta/#GSMR/.

## Methods

### UK Biobank genotyping and quality control

The United Kingdom Biobank (UKB) cohort is a population-based volunteer longitudinal cohort consisting of ∼500,000 individuals recruited at 22 centres across the United Kingdom^76^. Genotype data from these individuals were imputed using the Haplotype Reference Consortium (HRC) and UK10K as the reference sample. A European ancestry subset (456,414 individuals, including 348,501 unrelated individuals) was identified by projecting the UKB participants onto the 1000 Genome Project principal components coordinates. Genotype probabilities were converted to hard-call genotypes using PLINK2^77^ (hard-call 0.1) and single nucleotide polymorphisms (SNPs) with minor allele count < 5, Hardy-Weinberg equilibrium test P value < 1.0E-5, missing genotype rate > 0.05, or imputation accuracy (Info) score < 0.3 were excluded.

### Phenotype definition

The UKB phenotypes used in analyses were derived from two categories: one is disease-diagnoses from the combination of self-reported non-cancer illness code (UKB data field 20002) and ICD10 codes from hospital admission records (UKB data fields: 41202 and 41204). The other category is medication-taking based on treatment/medication code (UKB data field 20003; see also **Table S2**). For the GORD disease-diagnoses phenotype (39,851 cases and 416,563 controls), participants were classified as cases if they had either a self-reported (code: 1138) or ICD10 (code: K210 and K219) record for GORD. The remaining individuals were assigned as controls. PUD disease-diagnoses cases are a combination of stomach ulcer cases (self-reported code: 1142, ICD10 code: K250-K257 and K259), duodenal ulcer cases (self-reported code: 1457, ICD10 code: K260-K267 and K269) and other site peptic ulcer cases (self-reported code: 1400, ICD10 code: K270-K277 and K279). The remaining individuals were PUD controls. There were 12,226 cases and 444,188 controls for PUD. In clinical practice, medications for PUD also have a therapeutic effect on GORD. Thus, we first identified 54,541 individuals taking medications that are mainly considered medications for GORD/PUD (**Table S2**) and further combined these medication-taking cases with diseases-diagnoses cases for GORD and PUD, leaving a total of 75,192 cases (35,005 male cases) and 381,222 controls (phenotype abbreviation: GP_+_M – for GORD, PUD and corresponding medications). For the IBS disease-diagnoses phenotype (14,994 cases and 441,420 controls), case status was assigned according to the self-reported (code: 1154) or ICD10 (code: K580 and K589) record. The remaining individuals were coded as IBS controls. We then combined individuals taking medications for IBS (4,363 cases) with IBS disease-diagnoses cases, giving a total of 16,518 cases (4,317 male cases) and 439,896 controls and defined as a IBS_+_M phenotype (abbreviation for combination of IBS and medications for IBS). Following Mowat *et al.*^78^, inflammatory bowel disease (IBD) disease-diagnoses cases are a combination of Crohn’s diseases (self-reported code: 1462, ICD10 code: K500, K501, K508, and K509), ulcerative colitis (self-reported code: 1463, ICD10 code: K510-K515, K518 and K519) and other inflammatory bowel disease (self-reported code: 1461), giving a total of 6,115 cases and 450,299 controls. Since the medications for IBD can also be used to treat other diseases (i.e. IBD medications are not specific), we did not incorporate the medication data for the IBD phenotype. The Supplementary Data 1 of Wu *et al.*^66^ provides UKB medication classification based on Anatomical Therapeutic Chemical (ATC) Classification System^60^ and we extracted medications for GORD/PUD (the first two ATC level: A02) and IBS (the first two ATC level: A03) (**Table S2**). For the validation of UKB PUD genome-wide significant SNPs, we used Genetic Epidemiology Research on Aging (GERA)^79^ PUD summary statistics from Zhu *et al.*^48^. The definition of PUD phenotypes from the GERA cohort is described in The Supplementary Table 4 of Zhu *et al.*^48^. Briefly, the total 61,847 individuals were divided into two groups: 1,004 peptic ulcer cases according to ICD9 code (531-534) and 60,843 controls, respectively.

### Comorbidity analyses

Comorbidity analyses, including comorbidity among the four digestive diseases and comorbidity between each of the GORD, PUD, IBS and IBD with depression phenotypes in the UKB, were conducted in 348,501 unrelated individuals. Among the four digestive diseases, for each two of the GORD, PUD, IBS and IBD cases (6 pairs in total), we first checked whether the number of overlapped individuals between case groups is statistically significantly larger than the overlap expected by chance. For each of the GORD, PUD, IBS and IBD disease (defined as the index disease), we conducted competitive comorbidity analyses to test among the other three diseases which disease is more prone to be comorbid with the index disease. Briefly, we calculated the proportion of the index disease cases in the other three diseases respectively and compared them in pairs by using two-proportions *Z* test. One prerequisite for two-proportion *Z* test is two samples are independent of each other. Given this, we removed the overlapped cases among these three diseases when calculating the proportion of the index disease cases. For comorbidity between each of the GORD, PUD, IBS and IBD with depression phenotypes, We first derived 8 depression phenotypes based on different data field (20002, 20216, 2090, 20440, 20442, 2100, 41202, 41204) and mental health online follow-up (data category: 138) from the UKB according to Cai *et al.*^47^. The details for depression phenotype definition were described in the ref^47^. Briefly, depression phenotypes were defined according to help seeking behaviour and symptoms, including seen general practice (GP) for nerves, anxiety, tension or depression (abbreviation: Gppsy), seen psychiatrist for nerves, anxiety, tension or depression (abbreviation: Psypsy), probable recurrent major depression or single probable major depression episode (abbreviation: DepAll), self-reported depression (abbreviation: SelfRepDep), ICD10 defined depression (abbreviation: ICD10Dep), DSM V clinical guideline defined major depression (abbreviation: LifetimeMDD), major depression recurrence (abbreviation: MDDRecur) and seen GP for depression but no cardinal symptoms (abbreviation: GpNoDep) phenotypes. We then checked the overlap individuals from each of the GORD, PUD, IBS and IBD with the 8 depression phenotypes. For each of 32 digestion-depression phenotype pairs, we tested whether the number of individuals who are both digestion phenotype case and depression phenotype case is statistically significantly different from the expected number. Bonferroni correction was used to account for multiple testing.

### Full-sibling risk and heritability estimation

To demonstrate the genetic component for the UKB case-control phenotypes of GORD, PUD, IBS and IBD, we estimated the increased risk of the disorders in full-siblings of those affected (We didn’t incorporate other relative pairs data given the limited sample size for disease cases), compared to the risk in all UKB individuals (disease risk). As described by Bycroft *et al.*^76^, 22,665 full-sibling pairs were inferred from the kinship coefficients estimated using KING^80^. Risk ratio, 95% confidence interval (CI) and corresponding P value were calculated using fmsb R package (https://cran.r-project.org/web/packages/fmsb/fmsb.pdf). After obtaining the full-sibling relative risks, we used liability distribution theory^81-83^ to estimate the heritability of each trait, under the assumption that the increased risk only reflects shared genetic factors. As a sensitivity analysis, we repeated analyses after excluding any individual recorded to have more than one disorder.

### Genome-wide association study (GWAS) analyses

We performed case-control GWAS analyses using BOLT-LMM^84^ with sex, age and 20 ancestry principal components (PCs) fitted as covariates. 543,919 SNPs generated by LD pruning (*r*^2^ < 0.9) from Hapmap3 SNPs were used to control for population structure and polygenic effects, including genetic relatedness between individuals. The effect size (*β*) from BOLT-LMM on the observed 0-1 scale were transformed to odds ratio (OR) using the following equation^85^: 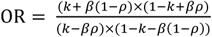, where *k* is the proportion of sample that are cases, and *ρ* is the allele frequency in the full UKB European cohort. The standard error (s.e.) for OR were then calculated based on the OR and P value from the initial GWAS using the formula 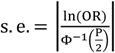. A total of 8,546,066 SNPs with minor allele frequency (MAF) > 0.01 were analysed. Quasi-independent trait-associated regions were generated through linkage disequilibrium (LD) clumping retaining the most associated SNP (lead SNP) in each region (PLINK (v1.90b)^77^ --clump-p1 5.0E-8 --clump-p2 5.0E-8 --clump-r2 0.01 --clump-kb 1000). In addition to the genome-wide significant SNPs identified through BOLT-LMM, the genotype data (8,546,066 SNPs with MAF > 0.01) of 348,501 unrelated European individuals were used to provide a LD reference. Due to the complexity of major histocompatibility complex (MHC) region (25Mb – 34Mb), only the most significant SNP across that region was reported. Regional visualisation plots were produced using LocusZoom^86^. The genomic inflation factor (λGC) was also reported for each phenotype. We used the GERA^79^ cohort PUD GWAS summary statistics^48^ for a validation look-up of the UKB PUD genome-wide significant SNPs. We also conducted pleiotropy (SNP associated with multiple traits) analysis. Briefly, we downloaded published GWAS associations from the GWAS Catalog^29^ on July 9 2019. For each of GORD, PUD, GP_+_M and IBS_+_M associated SNPs in our study (index SNP), we first selected SNPs from the GWAS Catalog within a ±1,000kb window size of the index SNP. We then selected the GWAS Catalog SNPs significantly associated (P < 5.0E-8) with either mental health-related traits or digestive diseases. We reported as a pleiotropic association if selected GWAS Catalog SNPs are in LD (*r*^2^ > 0.1) with the index SNP. Similarly, for IBD-associated SNPs in our study, we checked whether significant association (P < 5.0E-8) have been reported for inflammatory bowel diseases (including the subtypes) using the downloaded GWAS Catalog data.

### SNP-based heritability and genetic correlations of the six digestion phenotypes

Linkage Disequilibrium Score regression (LDSC)^30^ was used to estimate SNP-based heritability 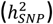 from the GWAS summary statistics. The 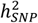 estimated on the observed scale were transformed to the liability scale taking the sample lifetime risk (proportion of sample that are cases) as the disease lifetime risk estimates. The summary statistics for each phenotype were filtered using the LDSC default file, w_hm3.snplist, with the default LD scores computed using 1000 Genomes European data (eur_w_ld_chr) as a reference. Genetic correlations (*r*_*g*_) between any two of the six UKB digestion phenotypes or each of the six phenotypes and six published psychiatric traits (attention deficit/hyperactivity disorder (ADHD)^31^, schizophrenia (SCZ)^32^, anxiety disorder^33^, posttraumatic stress disorder (PTSD)^34^, bipolar disorder (BIP)^35^ and autism spectrum disorder (ASD)^36^) were calculated using bivariate LDSC^87^. The *r*_*g*_ were also calculated between each of the six digestion phenotypes and 252 other traits using LD Hub^37^. As sensitivity analyses, we repeated analyses after excluding any individual recorded to have more than one disorder. Although removal of these individuals could make estimates of SNP-based heritability difficult to interpret, genetic correlations are more robust to such ascertainment^67^.

### Linking GWAS findings to gene expression

Following the estimation of 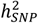, we partitioned the heritability by genomic features^88^. Briefly, in this method^88^, genetic variants are assigned into 53 functional categories using 24 publicly available annotation data sets, such as UCSC coding, UTRs, promoter and intronic regions^89^, conserved regions^90^ and functional genomic annotations constructed using ENCODE^91^ and Roadmap Epigenomics Consortium data^92^. The method evaluates the contribution of each functional category to the overall 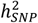 of a trait. A category is enriched for 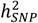 if the variants with high LD to that category have elevated *χ*^2^ statistics, compared to the expectation given the number of SNPs in the category. In another analysis, genetic variants were annotated to histone marks (H3K4me1, H3K4me3, H3K9ac and H3K27ac) by cell type specific classes and these annotations were allocated to 10 groups: adrenal and pancreas, central nervous system (CNS), cardiovascular, connective and bone, gastrointestinal, immune and hematopoietic, kidney, liver, skeletal muscle and other. We tested the enrichment of 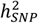 in tissues relevant to the 6 digestion phenotypes: the adrenal and pancreas, gastrointestinal, immune and hematopoietic and liver cell types. We also considered the CNS given the high *r*_*g*_ between the 5 of the six digestion phenotypes and depressive symptoms. We also used LDSC specific expressed genes (SEG)^41^ analysis to test the enrichment of 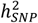 through gene expression derived cell-type specific annotations. First, given the strong contribution to GP_+_M 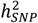 from the CNS and to IBD 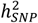 from the immune cell group, LDSC-SEG^41^ was applied to test the enrichment of 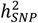 in 205 different tissues (53 from GTEx^42^ and 152 from Franke lab^43,44^). Second, given the observation that GORD, GP_+_M and IBS_+_M 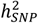 were enriched in the CNS, we also applied LDSC SEG to test the enrichment of 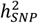 for GORD, GP_+_M and IBS_+_M in 13 brain regions using the multiple brain regions available in the GTEx study (https://gtexportal.org/home) data^42^ to identify specific brain regions implicated by the GWAS results for the three phenotypes. Bonferroni correction was used to account for multiple testing. We checked eQTL (expression quantitative trait loci, i.e. SNPs associated gene expression in different tissues) status for each genome-wide significant SNP using GTEx^42^ results for gastrointestinal tissue and brain tissues. Summary-data-based Mendelian Randomisation (SMR)^48^ was used to provide evidence for likely causal relationship between the trait-associated SNPs and gene expression. We used eQTLGen^93^ whole blood eQTL data since this is the largest eQTL data set and many eQTLs are shared across tissues^94^. To capture more tissue specific eQTL, we used GTEx^42^ eQTL data from 6 tissues: oesophagus-gastroesophageal junction, oesophagus mucosa, oesophagus muscularis, stomach, colon sigmoid, colon transverse tissue. We also used GTEx eQTL data from frontal cortex (BA9) tissue for GORD, PUD, GP_+_M, IBS and IBS_+_M given 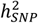 enrichment results. The Bonferroni corrected significance threshold was 0.05/155,059, where 155,059 is the number of total genes tested in SMR analyses. Because of its complexity, we do not report results of the MHC region (25Mb – 34Mb)^48^.

### Gene-based and gene-set enrichment analyses

MAGMA (v1.06)^46^ (Multi-marker Analysis of GenoMic Annotation) was used to test for gene-based association based on the SNP association results of the six digestion phenotypes. Gene length boundaries were defined as 35 kilobase (kb) upstream and 10 kb downstream from start and stop site, respectively, to include regulatory elements. The NCBI 37.3 build was used to assign the genetic variants to each gene. SNPs with MAF > 0.01 from 10,000 randomly sampled unrelated UKB European-ancestry individuals were used to provide a LD reference. A total of 18,402 genes were assessed for an association with each of the six digestion phenotypes with Bonferroni correction used to determine significance (α = 0.05/18402, P < 2.7E-6). We used the results obtained from gene-based analysis, together with curated gene sets (c2.all) and gene ontology sets (c5.bp, c5.cc, c5.mf) from MSigDB (v5.2)^95,96^ to conduct gene-set enrichment analyses. Competitive test P value for each gene set, as implemented in MAGMA, were computed taking gene size, density, minor allele count and gene-gene correlation into consideration^46^. False discovery rate (FDR)-adjusted P values for biological pathways for each of the six digestion phenotypes were generated using Benjamini and Hochberg’s method^97^ to account for multiple testing.

### Mendelian Randomisation

We applied the Generalised Summary-data-based Mendelian Randomisation (GSMR)^48^ method to explore the potentially causal effect of MD as an exposure on the six UKB digestion phenotypes as outcome traits (defined as forward direction). GSMR uses summary-level data to test for causal associations between a putative risk factor (exposure) and an outcome trait. Independent genome-wide significant SNPs from the MD GWAS (excluding UKB cohort)^49^ were used as the Mendelian Randomisation (MR) exposure instrument variables. The HEIDI outlier test^48^ was used to remove outlier pleiotropic genetic instruments associated with both exposure phenotype and outcome phenotype from the analysis. We also conducted reverse causation analysis (i.e. testing the opposite hypothesis that the six digestion phenotypes cause MD). However, GSMR guidelines advise the use of at least 10 independent lead SNPs as genetic instruments to achieve robust results. In order to test the effect of GORD (5 SNPs with P < 5.0E-8), PUD (4 SNPs), IBS (0 SNP) and IBS_+_M (3 SNPs) on MD, we relaxed the significance threshold to allow for at least 10 SNPs for each of the four phenotypes. The other parameters were set to software defaults. For comparison, we also conducted IVW-MR^98^, MR-Egger^99^, weighted median-MR^100^ and MR-PRESSO^101^ analyses following the STROBE-MR guideline^102^.

### Genetic risk score (GRS) prediction

We used MD GWAS summary statistics^49^ (European ancestry, excluding UKB cohort) as discovery data to predict GP_+_M risk (risk for GORD, PUD and likelihood for taking GORD, PUD drugs). The MD data SNPs were matched with the GP_+_M SNPs (7,156,547 SNPs), then LD pruned and “clumped”, discarding variants within 1,000kb of, and in *r*^2^ *≥* 0.1 with, another (more significant) marker using SNPs with MAF > 0.01 from 10,000 random sampled unrelated UKB European-ancestry individuals as the LD reference. GRS of GP_+_M sample individuals were generated for a range of MD GWAS summary statistics data association P value thresholds (5.0E-8, 1.0E-5, 1.0E-4, 1.0E-3, 1.0E-2, 0.05, 0.1, 0.5). For each discovery-target pair, three outcome variables were calculated. (1) The P value of case-control GRS difference from logistic regression. (2) Area under the receiver operator characteristic curve using R package pROC^103^, which can be interpreted as the probability of ranking a randomly chosen case higher than a randomly chosen control. (3) Odds ratio and 95% confidence interval for the 2^nd^ to 10^th^ GRS deciles group compared with 1^st^ decile. We also used UKB PUD GWAS summary statistics to calculate genetic risk score based on P value threshold 5E-8 for individuals from GERA cohort and conducted out-of-sample GRS PUD prediction for GERA individuals following same analyses above.

## Acknowledgements

We thank members of Program in Complex Trait Genomics group in The University of Queensland. This research was supported by the Australian National Health and Medical Research Council (Grant Number: 1113400, 1078901, 1078037). Yeda Wu thanks the University of Queensland Senate for the F.G. Meade Scholarship and UQ Research Training Scholarship. This study makes use of data from UK Biobank (ID: 12505) and we thank the UK Biobank participants and the UKB research teams for their generous contributions to generating an important research resource. We also use data from the Resource for Genetic Epidemiology Research on Adult Health and Aging (GERA: dbGaP phs000674.v2.p2) study which was supported by grant RC2 AG036607 from the National Institute of Health, grants from Robert Wood Johnson Foundation, the Ellison Medical Foundation, the Wayne and Gladys Valley Foundation, and Kaiser Permanente. We thank the Kaiser Permanente Medical Care Plan, Northern California Region (KPNC) members who have generously agreed to participate in the Kaiser Permanente Research Program on Genes, Environment and Health (RPGEH). We thank 23andMe for the use of GWAS summary statistics for major depression that include data from 23andMe.

## Author Contribution

N.R.W., P.M.V. and Y.W conceived and designed the experiment. Y.W. performed the analysis with assistance and guidance from G.K.M., E.M.B., J.S. contributed to data quality control of UKB data. Y.W., N.R.W. and P.M.V. wrote the manuscript with the participant of all authors.

## Data availability

Summary statistics are available at http://cnsgenomics.com/data.html. The data that support the findings of this study are available from UK Biobank (http://www.ukbiobank.ac.uk/about-biobank-uk/). Restrictions apply to the availability of these data, which were used under license for the current study (ID: 12505). Data are available for bona fide researchers upon application to the UK Biobank. We also used peptic ulcer disease GWAS summary statistics (https://cnsgenomics.com/data.html) from the Resource for the Genetic Epidemiology Research on Adult Health and Aging (GERA: dbGaP phs000674.v2.p2) study. We used GWAS summary statistics for major depression that include data from 23andMe. These data can be obtained by qualified researchers under an agreement with 23andMe that protects the privacy of the 23andMe participant 23andMe. Researchers can perform meta-analysis of 23andMe summary statistics and the other five-cohort results file, as described in Wray *et al.*^49^, to get major depression GWAS summary statistics (excluding UK Biobank cohort). The data for generating the figures are provided in the Supplementary Materials.

## Code availability

All the code for the analyses in this study are at http://cnsgenomics.com/data.html.

