## Supplementary notes, tables and figures for "Genome-wide association study of gastrointestinal disorders reinforces the link between the digestive tract and the nervous system"

### *Supplementary Note*

Supplementary Note 1

As a sensitivity analysis, SNP-based heritability ($h^2_{SNP}$) and genetic correlation ($r_g$) analyses were conducted for GORD, PUD, IBS and IBD using the phenotypes generated after excluding individuals with more than one of the four gastrointestinal (GI) disorders (defined as sensitivity analysis phenotypes). The number of overlapped individuals for GORD, PUD, IBS and IBD case are in **Figure S4A**. As expected, the $h^2_{SNP}$ estimates on the observed scale for these disorders were lower due to excluded case individuals but still significantly different from zero (**Figure S4B**). We then calculated the $r_g$ between these sensitivity analysis phenotypes with the original phenotypes, within sensitivity analysis phenotypes, between sensitivity analysis phenotypes with traits from LD Hub and six published mental disorder studies. Each of the sensitivity analysis phenotypes is highly genetically correlated with corresponding original phenotype, as expected given the part/whole relationship, but included for completeness (**Figure S4C**). The $r_g$ within sensitivity analysis phenotypes showed high $r_g$ among GORD, PUD, IBS and all low non-statistically significant $r_g$ with IBD. The $r_g$ between GORD and PUD is 0.42 (SE = 0.08, $P_{H0:rg=0}$ = 2.4E-7) and the $r_g$ between GORD and IBS is 0.47 (SE = 0.08, $P_{H0:rg=0}$ = 8.5E-10), which are lower to the original results. The $r_g$ between PUD and IBS is 0.25 (SE = 0.12, $P_{H0:rg=0}$ = 0.036), of which the original $r_g$ is 0.48 (SE = 0.10, $P_{H0:rg=0}$ = 8.0E-7). This difference is due to the sample size. As shown in **Figure S4D**, the number of overlapped individuals for PUD and IBS is 776, however, we over-removed 3,030 individuals for PUD and 3,048 individuals for IBS due to the overlap with the other two GI disorders (GORD and IBD, shown in **Figure S4A**). The number of total PUD and IBS cases is 12,226 and 14,994 and these over-removed individuals occupy ~1/4 for PUD cases and ~1/5 for IBS cases, resulting in reducing power to estimate the $r_g$. Thus we only removed the only 776 overlapped individuals for PUD and IBS and re-calculated the $r_g$ between PUD and IBS, as shown in **Figure S4E**. The $r_g$ is 0.38 (SE = 0.11, $P_{H0:rg=0}$ = 5.0E-4). GORD, PUD and IBS sensitivity analyses phenotypes showed statistically significantly $r_g$ with depressive symptoms while there is no statistically significant $r_g$ between IBD and depressive symptoms. The $h^2_{SNP}$ for sensitivity analyses phenotypes are in **Table S7**. The $r_g$ between sensitivity analysis phenotypes with the original phenotypes, within sensitivity analysis phenotypes and traits from the six published mental disorder studies are in **Table S9**. The $r_g$ between sensitivity analysis phenotypes with traits from LD Hub are in **Supplementary Data 4**.

Supplementary Note 2

Given the statistically significant results between MD and $GP_+M$ from bidirectional GSMR analyses, we also conducted bidirectional MR analyses between MD and $GP_+M$ using TwoSampleMR package. For each of MD and $GP_+M$ GWAS summary statistics, we first generated the quasi-independent loci using PLINK(v1.90b)[1] (--clump-p1 5.0E-8 –clump-p2 5.0E-8 –clump-r2 0.01 –clump-kb 1000) and the genotype data (8,545,066 SNPs with MAF > 0.01) of 348,501 unrelated European individuals as a LD reference. Only the most significant SNP across the MHC region was retained. For each of the genetic instrument, we extracted the allele, effect size, standard error and P value from the exposure GWAS summary statistics. We also extracted the corresponding information from the outcome GWAS summary statistics for these genetic instruments. If a SNP was unavailable in the outcome GWAS summary statistics, we identified proxy SNP with a minimum LD $r^2$ = 0.8. We performed harmonisation of the direction of effects between exposure and outcome association (action = 2). For each direction of potential influence, we combined MR estimates using inverse variance-weighted (IVW) analysis[2], which essentially translates to a weighted regression of SNP-outcome effects on SNP-exposure effects where the intercept is constrained to zero. The IVW method will return an unbiased estimate if there is no or balanced horizontal pleiotropy. To account for this, we compared results from IVW method with results from MR Egger[3] and weighted median method[4], from which the estimates are known to be relatively robust to horizontal pleiotropy, though at the cost of reduced statistical power. To assess robustness of significant results, we also conducted the MR Egger intercept test for horizontal pleiotropy. We also applied MR-PRESSO (Pleiotropy Residual Sum and Outlier)[5] to detect and correct for any outliers reflecting likely pleiotropic biases for all reported results. The IVW results showed bidirectionally statistically significant results, of which the pattern is similar as the GSMR results (**Figure S9**). The MR-Egger intercept test showed no statistical significance, suggesting that there is no horizontal pleiotropy (**Table S16**). There is no outliers being removed after MR-PRESSO analyses.

#### Supplementary Note 3

We noticed a previous study report that rs10512344 is the only one SNP which are genome-wide significantly associated (reported P = 3.6E-8) with female IBS using UKB data[6]. Both the IBS (lead SNP: rs112243849, P = 7.5E-8, **Figure S2A**) and $IBS^{+}M$ (lead SNP: rs7861675, P = 1.2E-7, **Figure 2D**) phenotypes in our study reconfirmed the association between this locus and IBS at the genome-wide suggestive level. That study stated that among those unrelated individuals with British ancestry in UKB, self-reported IBS individuals were selected as cases and the remaining as controls. The study further removed individuals with IBD and celiac disease for both cases and controls and IBS for controls using self-reported illness data (data field: 20002) and ICD10 main diagnoses (data field: 41002)[6]. In our analysis, we integrated whole European ancestry individuals with both self-reported IBS diagnoses, ICD10 main and secondary IBS diagnoses (data field: 41202 and 41204) and taking corresponding medications as cases while the others as controls (**Methods**) to gain more statistical power. This may lead to slight difference of statistically significant level for a particular locus but increased power to detect more SNP associations, though only three in our analysis as shown in **Figure 2D**.

#### *Supplementary Tables*

Table S1. Sensitivity analysis[1] of full-sibling relative risk and heritability estimation for GORD, PUD, IBS and IBD

| Digestion phenotypes | GORD | PUD | IBS | IBD |
|---|---|---|---|---|
| N case:N control[1] | 33527:416563 | 8420:444188 | 11170:441420 | 4742:450299 |
| N case/(N case + N control) | 0.074 | 0.019 | 0.025 | 0.010 |
| No. of male case:No. of female case | 15642:17885 | 4995:3425 | 2746:8424 | 2321:2421 |
| No. of male control:No of female control | 190500:226063 | 201807:242381 | 205006:236414 | 205871:244428 |
| Odds of being male case/Odds of being female case | 0.082:0.079 | 0.025:0.014 | 0.013:0.036 | 0.011:0.010 |
| Male:Female odds ratio for being case | 1.04 | 1.75 | 0.38 | 1.14 |
| No. of full-sibling pairs where both proband and full-sibling are cases | 356 | 24 | 42 | 22 |
| No. of full-sibling pairs where only the proband is a case | 2993 | 744 | 1083 | 432 |
| Full-sibling relative risk (95% confidence interval) | 1.43 (1.29-1.58) | 1.68 (1.13-2.49) | 1.51 (1.12-2.04) | 4.65 (3.09-7.00) |
| Heritability (95% confidence interval)[2] | 0.21 (0.15-0.28) | 0.19 (0.04-0.34) | 0.16 (0.04-0.29) | 0.53 (0.37-0.71) |

[1] The individuals recorded to have more than one disorder were removed.
[2] The corresponding lower and upper values of 95% confidence interval (CI) for risk in full-sibling were used to calculate the 95% CI for heritability estimation.

Table S2. Medications for gastro-oesophageal reflux disease (GORD), peptic ulcer disease (PUD) and irritable bowel syndrome (IBS) in UK Biobank

| Category | Coding | Active ingredient | ATC code[1] |
|---|---|---|---|
| Maalox tablet | 1140865358 | Aluminum Hydroxide, Magnesium Hydroxide, Simethicone | A02A |
| Mucogel suspension | 1140865368 | Aluminum Hydroxide, Magnesium Hydroxide | A02A |
| Co-magaldrox | 1140881318 | Aluminum Hydroxide, Magnesium Hydroxide | A02A |
| Magnesium trisilicate | 1140881324 | Magnesium Trisilicate | A02A |
| Milk of magnesia suspension | 1140881550 | Magnesium Hydroxide | A02AA |
| Magnesium carbonate | 1140881320 | Magnesium Carbonate | A02AA01 |
| Magnesium hydroxide | 1140881330 | Magnesium Hydroxide | A02AA04 |
| Rennie duo oral suspension | 1141166086 | Calcium Carbonate | A02AC01 |
| Asilone liquid | 1140881422 | Magnesium Oxide, Aluminum Hydroxide, Dimethicone | A02AF |
| Maalox plus suspension | 1140865366 | Aluminum Hydroxide, Magnesium Hydroxide | A02AF02 |
| Gastrocote liquid | 1140881414 | Sodium Bicarbonate, Aluminum Hydroxide, Alginic Acid, Magnesium Trisilicate | A02AX |
| Gastrocote s/f liquid | 1140928346 | Sodium Bicarbonate, Aluminum Hydroxide, Alginic Acid, Magnesium Trisilicate | A02AX |
| Peptac liquid | 1141168752 | Sodium Bicarbonate, Calcium Carbonate, Alginic Acid | A02AX |
| Gavilast-p 75mg tablet | 1141188426 | Ranitidine | A02BA |
| Cimetidine | 1140865426 | Cimetidine | A02BA01 |
| Tagamet 100 tablet | 1140909500 | Cimetidine | A02BA01 |
| Ranitidine | 1140879406 | Ranitidine | A02BA02 |
| Zantac 75 tablet | 1140916980 | Ranitidine | A02BA02 |
| Famotidine | 1140865608 | Famotidine | A02BA03 |
| Pepcid ac indigestion tablet | 1140909496 | Famotidine | A02BA03 |
| Nizatidine | 1140865618 | Nizatidine | A02BA04 |
| Misoprostol | 1140865628 | Misoprostol | A02BB01 |
| Omeprazole | 1140865634 | Omeprazole | A02BC01 |
| Losec 10mg capsule | 1140909578 | Omeprazole | A02BC01 |
| Pantoprazole | 1140929012 | Pantoprazole | A02BC02 |
| Protium 20mg e/c tablet | 1141164616 | Pantoprazole | A02BC02 |
| Lansoprazole | 1140864752 | Lansoprazole | A02BC03 |
| Zoton 15mg capsule | 1140923688 | Lansoprazole | A02BC03 |
| Rabeprazole sodium | 1141168584 | Rabeprazole | A02BC04 |

| | | | |
|---|---|---|---|
| Pariet 10mg e/c tablet | 1141168590 | Rabeprazole | A02BC04 |
| Esomeprazole | 1141177526 | Esomeprazole | A02BC05 |
| Nexium 20mg tablet | 1141177532 | Esomeprazole | A02BC05 |
| Acidex oral suspension | 1141172224 | Sodium Bicarbonate, Calcium Carbonate, Alginic Acid | A02BX |
| Sucralfate | 1140865536 | Sucralfate | A02BX02 |
| Antepsin 1g tablet | 1140865538 | Sucralfate | A02BX02 |
| Gaviscon liquid | 1140865354 | Sodium Bicarbonate, Calcium Carbonate, Alginic Acid | A02BX13 |
| Topal tablet | 1140865370 | Aluminum Hydroxide, Magnesium Carbonate, Alginic Acid | A02BX13 |
| Kolanticon gel | 1140865380 | Aluminum Hydroxide, Simethicone, Dicycloverine Hydrochloride, Light Magnesium Oxide | A03A |
| Mintec 0.2ml e/c capsule | 1140865418 | Peppermint Oil | A03A |
| Mebeverine hcl+ispaghula 135mg/3.5g/sachet granules | 1140865408 | Mebeverine Hydrochloride | A03AA04 |
| Mebeverine | 1140879428 | Mebeverine | A03AA04 |
| Colofac-100 tablet | 1141167334 | Mebeverine | A03AA04 |
| Dicyclomine hydrochloride | 1140865378 | Dicyclomine | A03AA07 |
| Merbentyl 10mg tablet | 1140865382 | Dicyclomine | A03AA07 |
| Dicycloverine | 1141194852 | Dicyclomine | A03AA07 |
| Pro-banthine 15mg tablet | 1140865330 | Propantheline | A03AB05 |
| Propantheline | 1140883480 | Propantheline | A03AB05 |
| Spasmonal 60mg capsule | 1140865336 | Alverine | A03AX08 |
| Alverine | 1140879424 | Alverine | A03AX08 |
| Spasmonal fibre granules 500g | 1141166264 | Alverine | A03AX08 |
| Alverine citrate+sterculia 0.5%/62% granules | 1140865338 | Alverine, Sterculia | A03AX58 |
| Atropine | 1140883494 | Atropine | A03BA01 |
| Hyoscine butylbromide | 1140865394 | Scopolamine Butylbromide | A03BB01 |
| Buscopan 10mg tablet | 1140865396 | Scopolamine Butylbromide | A03BB01 |

[1] The Supplementary Data 1 of Wu *et al.*[7] provides UKB medication classification based on Anatomical Therapeutic Chemical (ATC) Classification System[8] and we extracted medications for GORD/PUD (the first two ATC level: A02) and IBS (the first two ATC level: A03) as listed above.

Table S3. Genome-wide significant SNPs associated with inflammatory bowel disease (IBD) in UK Biobank

| Digestion Phenotypes | SNP | CHR. | BP | A1/A2 | A1 Frequency | OR | P | Reported SNP[1] | PMID |
|---|---|---|---|---|---|---|---|---|---|
| IBD | rs138753323 | 6 | 32173599 | T/C | 0.97 | 0.59 | 1.5E-36 | rs17207986 | 20848476 |
| | rs11581607 | 1 | 67707690 | G/A | 0.93 | 1.53 | 1.9E-22 | rs11209026 | 18758464 |
| | rs2836878 | 21 | 40465534 | G/A | 0.73 | 1.22 | 1.2E-20 | rs2836878 | 18758464 |
| | rs6017342 | 20 | 43065028 | A/C | 0.48 | 0.87 | 4.0E-15 | rs6017342 | 28067908 |
| | rs10737482 | 1 | 20173858 | T/C | 0.39 | 0.86 | 7.4E-15 | rs6426833 | 28067908 |
| | rs3024493 | 1 | 206943968 | C/A | 0.85 | 0.84 | 2.3E-13 | rs3024505 | 28067908 |
| | rs6671847 | 1 | 161478810 | G/A | 0.49 | 1.14 | 1.7E-12 | rs1801274 | 28067908 |
| | rs1297261 | 21 | 16812623 | T/C | 0.57 | 1.14 | 6.0E-12 | rs2823286 | 28067908 |
| | rs2212434 | 11 | 76281593 | C/T | 0.55 | 0.88 | 1.4E-11 | rs11236797 | 28067908 |
| | rs11403745 | 10 | 101282604 | C/CA | 0.48 | 1.13 | 2.3E-11 | rs4409764 | 28067908 |
| | rs72810950 | 10 | 90822640 | T/C | 0.90 | 1.24 | 1.4E-10 | - | - |
| | rs905634 | 1 | 200884985 | C/T | 0.69 | 1.13 | 4.4E-10 | rs7554511 | 28067908 |
| | rs10761659 | 10 | 64445564 | A/G | 0.46 | 0.89 | 1.2E-09 | rs10761659 | 28067908 |
| | rs2066847 | 16 | 50763778 | G/GC | 0.98 | 0.69 | 1.3E-09 | rs2066847 | 20570966 |
| | 9:5057580 | 9 | 5057580 | CTTT/C | 0.56 | 0.89 | 1.8E-09 | rs10758669 | 23128233 |
| | rs12128452 | 1 | 20143706 | T/G | 0.61 | 1.12 | 2.3E-09 | rs3806308 | 19122664 |
| | rs34893377 | 3 | 49515972 | T/TA | 0.69 | 0.89 | 3.0E-09 | rs9836291 | 26192919 |
| | rs1521186 | 1 | 151784547 | G/A | 0.55 | 0.90 | 3.5E-09 | - | - |
| | rs138182176 | 5 | 40434755 | A/AAATAT | 0.39 | 0.90 | 1.0E-08 | rs1992661 | 26974007 |
| | rs10799837 | 1 | 20135612 | G/A | 0.43 | 1.11 | 1.2E-08 | rs1317209 | 20228799 |
| | rs34926204 | 7 | 107488726 | T/C | 0.56 | 1.11 | 1.6E-08 | rs6466198 | 26974007 |
| | rs35788599 | 12 | 68476749 | G/C | 0.62 | 0.90 | 1.9E-08 | rs11614178 | 26974007 |
| | rs2816972 | 1 | 200085004 | A/G | 0.11 | 0.84 | 3.6E-08 | rs2816958 | 26974007 |

[1] The associations between SNPs from "Reported SNP" column and IBD have been reported by other studies (corresponding PMID column). These SNPs are either same as IBD-associated SNPs in our UK Biobank analysis or in linkage disequilibrium with our IBD-associated SNPs in UK Biobank. For the detailed statistics for those reported SNPs, please refer to **Supplementary Data 2**.

Table S4. GERA peptic ulcer GWAS summary statistics for SNPs associated with UKB peptic ulcer diseases (P < 5e-8).

| SNP | CHR. | BP | Nearest gene[1] | UKB statistics | | | | GERA statistics | | | |
|---|---|---|---|---|---|---|---|---|---|---|---|
| | | | | A1/A2 | A1 frequency | OR[2] | P | A1/A2 | A1 frequency | OR[2] | P |
| rs681343 | 19 | 49206462 | *FUT2* | C/T | 0.49 | 0.91 | 6.9E-13 | C/T | 0.51 | 0.84 | 5.0E-4 |
| rs147048677 | 1 | 155161794 | *MUC1* | C/T | 0.94 | 0.84 | 1.6E-11 | C/T | 0.95 | 0.88 | 0.20 |
| rs2976388 | 8 | 143760256 | *JRK; PSCA* | G/A | 0.58 | 1.09 | 7.4E-11 | G/A | 0.57 | 1.08 | 0.12 |
| rs10500661 | 11 | 6273744 | *CNGA4; CCKBR* | T/C | 0.80 | 0.91 | 6.7E-10 | T/C | 0.79 | 0.90 | 0.08 |

[1] SNPs are annotated if they are within a gene region or there are only a few genes around. For full gene location please refer to the locus zoom plot (**Supplementary Data 1**).

[2] Odds ratio (OR) is for risk of A1 allele compared to A2 allele.

Table S5. SNP-based heritability estimates parameters from LD score regression

| Digestion Phenotypes | $\lambda_{GC}$ | Intercept (s.e.) | h2_observed (s.e.) | Population lifetime risk[1] | h2_liability (s.e.) |
|---|---|---|---|---|---|
| GORD | 1.2 | 1.02 (0.008) | 0.02 (0.002) | 0.087 | 0.08 (0.005) |
| PUD | 1.1 | 1.01 (0.006) | 0.01 (0.001) | 0.027 | 0.05 (0.008) |
| $GP_{+}M$ | 1.4 | 1.04 (0.008) | 0.04 (0.002) | 0.165 | 0.10 (0.004) |
| IBS | 1.1 | 1.01 (0.007) | 0.01 (0.001) | 0.033 | 0.06 (0.008) |
| $IBS_{+}M$ | 1.1 | 1.01 (0.008) | 0.01 (0.001) | 0.036 | 0.07 (0.008) |
| IBD | 1.1 | 1.03 (0.008) | 0.01 (0.002) | 0.013 | 0.12 (0.017) |

[1] We used the proportion of sample that are cases as the estimates of population lifetime.
Abbreviation: s.e.: standard error

Table S6. LDSC genetic correlation estimates between each pair of the six digestion phenotypes (see **Figure 3B**)

| Trait1 | Trait2 | rg | se | p | h2 (obs) | h2 se (obs) | h2 int | h2 int se | gcov int | gcov int se |
|---|---|---|---|---|---|---|---|---|---|---|
| GORD | GORD | 1.00 | 0.000 | 0.0E+00 | 0.02 | 0.002 | 1.02 | 0.008 | 1.02 | 0.008 |
| GORD | PUD | 0.65 | 0.059 | 6.5E-28 | 0.01 | 0.001 | 1.01 | 0.007 | 0.10 | 0.005 |
| GORD | GP+M | 0.97 | 0.011 | 0.0E+00 | 0.04 | 0.002 | 1.04 | 0.008 | 0.72 | 0.007 |
| GORD | IBS | 0.61 | 0.057 | 1.5E-26 | 0.01 | 0.001 | 1.01 | 0.007 | 0.07 | 0.005 |
| GORD | IBS+M | 0.62 | 0.052 | 3.6E-32 | 0.01 | 0.001 | 1.01 | 0.008 | 0.08 | 0.005 |
| GORD | IBD | 0.14 | 0.061 | 2.2E-02 | 0.01 | 0.002 | 1.03 | 0.008 | 0.01 | 0.005 |
| PUD | GORD | 0.65 | 0.059 | 6.5E-28 | 0.02 | 0.002 | 1.02 | 0.008 | 0.10 | 0.005 |
| PUD | PUD | 1.00 | 0.000 | 0.0E+00 | 0.01 | 0.001 | 1.01 | 0.007 | 1.01 | 0.007 |
| PUD | GP+M | 0.75 | 0.042 | 5.8E-72 | 0.04 | 0.002 | 1.04 | 0.008 | 0.38 | 0.005 |
| PUD | IBS | 0.48 | 0.097 | 8.0E-07 | 0.01 | 0.001 | 1.01 | 0.007 | 0.03 | 0.005 |
| PUD | IBS+M | 0.50 | 0.093 | 5.0E-08 | 0.01 | 0.001 | 1.01 | 0.008 | 0.04 | 0.005 |
| PUD | IBD | 0.17 | 0.094 | 7.1E-02 | 0.01 | 0.002 | 1.03 | 0.008 | 0.02 | 0.005 |
| GP+M | GORD | 0.97 | 0.011 | 0.0E+00 | 0.02 | 0.002 | 1.02 | 0.008 | 0.72 | 0.007 |
| GP+M | PUD | 0.75 | 0.042 | 5.8E-72 | 0.01 | 0.001 | 1.01 | 0.007 | 0.38 | 0.005 |
| GP+M | GP+M | 1.00 | 0.000 | 0.0E+00 | 0.04 | 0.002 | 1.04 | 0.008 | 1.04 | 0.008 |
| GP+M | IBS | 0.64 | 0.052 | 1.7E-34 | 0.01 | 0.001 | 1.01 | 0.007 | 0.09 | 0.006 |
| GP+M | IBS+M | 0.64 | 0.048 | 4.5E-40 | 0.01 | 0.001 | 1.01 | 0.008 | 0.11 | 0.006 |
| GP+M | IBD | 0.17 | 0.050 | 8.0E-04 | 0.01 | 0.002 | 1.03 | 0.008 | 0.03 | 0.006 |
| IBS | GORD | 0.61 | 0.057 | 1.5E-26 | 0.02 | 0.002 | 1.02 | 0.008 | 0.07 | 0.005 |
| IBS | PUD | 0.48 | 0.097 | 8.0E-07 | 0.01 | 0.001 | 1.01 | 0.007 | 0.03 | 0.005 |
| IBS | GP+M | 0.64 | 0.052 | 1.7E-34 | 0.04 | 0.002 | 1.04 | 0.008 | 0.09 | 0.006 |
| IBS | IBS | 1.00 | 0.000 | 0.0E+00 | 0.01 | 0.001 | 1.01 | 0.007 | 1.01 | 0.007 |
| IBS | IBS+M | 1.00 | 0.004 | 0.0E+00 | 0.01 | 0.001 | 1.01 | 0.008 | 0.96 | 0.007 |
| IBS | IBD | 0.17 | 0.078 | 2.5E-02 | 0.01 | 0.002 | 1.03 | 0.008 | 0.04 | 0.005 |
| IBS+M | GORD | 0.62 | 0.052 | 3.6E-32 | 0.02 | 0.002 | 1.02 | 0.008 | 0.08 | 0.005 |
| IBS+M | PUD | 0.50 | 0.093 | 5.0E-08 | 0.01 | 0.001 | 1.01 | 0.007 | 0.04 | 0.005 |
| IBS+M | GP+M | 0.64 | 0.048 | 4.5E-40 | 0.04 | 0.002 | 1.04 | 0.008 | 0.11 | 0.006 |
| IBS+M | IBS | 1.00 | 0.004 | 0.0E+00 | 0.01 | 0.001 | 1.01 | 0.007 | 0.96 | 0.007 |
| IBS+M | IBS+M | 1.00 | 0.000 | 0.0E+00 | 0.01 | 0.001 | 1.01 | 0.008 | 1.01 | 0.008 |
| IBS+M | IBD | 0.21 | 0.073 | 4.0E-03 | 0.01 | 0.002 | 1.03 | 0.008 | 0.04 | 0.005 |
| IBD | GORD | 0.14 | 0.061 | 2.2E-02 | 0.02 | 0.002 | 1.02 | 0.008 | 0.01 | 0.005 |
| IBD | PUD | 0.17 | 0.094 | 7.1E-02 | 0.01 | 0.001 | 1.01 | 0.007 | 0.02 | 0.005 |
| IBD | GP+M | 0.17 | 0.050 | 8.0E-04 | 0.04 | 0.002 | 1.04 | 0.008 | 0.03 | 0.006 |
| IBD | IBS | 0.17 | 0.078 | 2.5E-02 | 0.01 | 0.001 | 1.01 | 0.007 | 0.04 | 0.005 |
| IBD | IBS+M | 0.21 | 0.073 | 4.0E-03 | 0.01 | 0.001 | 1.01 | 0.008 | 0.04 | 0.005 |
| IBD | IBD | 1.00 | 0.000 | 0.0E+00 | 0.01 | 0.002 | 1.03 | 0.008 | 1.03 | 0.008 |

Table S7. SNP-based heritability estimates parameters from LD score regression for sensitivity analyses (see **Figure S4**)

| Digestion Phenotypes | $\lambda_{GC}$ | Intercept (s.e.) | h2_observed (s.e.) | Population lifetime risk[1] | h2_liability (s.e.) |
|---|---|---|---|---|---|
| GORD (RO) | 1.15 | 1.01 (0.008) | 0.020 (0.001) | 0.074 | 0.068 (0.005) |
| PUD (RO) | 1.05 | 1.01 (0.006) | 0.006 (0.001) | 0.019 | 0.052 (0.010) |
| PUD (RTO) | 1.05 | 1.01 (0.007) | 0.007 (0.001) | 0.025 | 0.049 (0.008) |
| IBS (RO) | 1.05 | 1.01 (0.007) | 0.007 (0.001) | 0.025 | 0.052 (0.009) |
| IBS (RTO) | 1.10 | 1.01 (0.007) | 0.010 (0.001) | 0.031 | 0.059 (0.008) |
| IBD (RO) | 1.05 | 1.02 (0.008) | 0.009 (0.002) | 0.010 | 0.118 (0.020) |

[1] We used the proportion of sample that are cases as the estimates of population lifetime.
Abbreviation: s.e.: standard error; RO: remove individuals with more than one of GORD, PUD, IBS and IBD disorders; RTO: remove the overlapped individuals with both PUD and IBS

Table S8. LDSC genetic correlation estimates between each of the six digestion phenotypes and 6 psychiatric traits (see **Figure 3B**)

| Trait1 | Trait2 | rg | se | p | h2 (obs) | h2 s.e. (obs) | h2 int | h2 int s.e. | gcov int | gcov int s.e. |
|---|---|---|---|---|---|---|---|---|---|---|
| GORD[1] | ADHD | 0.49 | 0.042 | 6.7E-32 | 0.24 | 0.016 | 1.03 | 0.010 | 0.00 | 0.006 |
| GORD | SCZ | 0.02 | 0.028 | 4.5E-01 | 0.42 | 0.015 | 1.05 | 0.012 | 0.01 | 0.006 |
| GORD | Anxiety | 0.33 | 0.134 | 1.3E-02 | 0.06 | 0.026 | 1.00 | 0.007 | 0.01 | 0.005 |
| GORD | PTSD | 0.27 | 0.128 | 3.6E-02 | 0.05 | 0.022 | 0.99 | 0.007 | 0.00 | 0.005 |
| GORD | BIP | -0.05 | 0.037 | 1.7E-01 | 0.34 | 0.015 | 1.02 | 0.008 | 0.00 | 0.006 |
| GORD | ASD | -0.01 | 0.044 | 8.3E-01 | 0.20 | 0.015 | 1.00 | 0.008 | 0.01 | 0.006 |
| PUD | ADHD | 0.54 | 0.067 | 9.0E-16 | 0.24 | 0.016 | 1.03 | 0.010 | 0.00 | 0.006 |
| PUD | SCZ | 0.14 | 0.044 | 1.1E-03 | 0.42 | 0.015 | 1.05 | 0.012 | 0.00 | 0.006 |
| PUD | Anxiety | 0.22 | 0.203 | 2.7E-01 | 0.06 | 0.026 | 1.00 | 0.007 | 0.01 | 0.005 |
| PUD | PTSD | 0.44 | 0.192 | 2.3E-02 | 0.05 | 0.022 | 0.99 | 0.007 | 0.00 | 0.004 |
| PUD | BIP | 0.04 | 0.053 | 4.8E-01 | 0.34 | 0.015 | 1.02 | 0.008 | 0.00 | 0.005 |
| PUD | ASD | 0.03 | 0.071 | 6.4E-01 | 0.20 | 0.015 | 1.00 | 0.008 | 0.00 | 0.005 |
| $GP_{+}M$ | ADHD | 0.56 | 0.035 | 2.4E-58 | 0.24 | 0.016 | 1.03 | 0.010 | 0.01 | 0.008 |
| $GP_{+}M$ | SCZ | 0.03 | 0.023 | 2.0E-01 | 0.42 | 0.015 | 1.05 | 0.012 | 0.01 | 0.006 |
| $GP_{+}M$ | Anxiety | 0.40 | 0.134 | 3.2E-03 | 0.06 | 0.026 | 1.00 | 0.007 | 0.01 | 0.005 |
| $GP_{+}M$ | PTSD | 0.35 | 0.112 | 2.0E-03 | 0.05 | 0.022 | 0.99 | 0.007 | 0.00 | 0.005 |
| $GP_{+}M$ | BIP | -0.01 | 0.031 | 7.2E-01 | 0.34 | 0.015 | 1.02 | 0.008 | 0.00 | 0.006 |
| $GP_{+}M$ | ASD | 0.06 | 0.038 | 9.5E-02 | 0.20 | 0.015 | 1.00 | 0.008 | 0.00 | 0.006 |
| IBS | ADHD | 0.32 | 0.064 | 5.3E-07 | 0.24 | 0.016 | 1.03 | 0.010 | 0.00 | 0.007 |
| IBS | SCZ | 0.15 | 0.046 | 8.0E-04 | 0.42 | 0.015 | 1.05 | 0.012 | -0.01 | 0.007 |
| IBS | Anxiety | 0.29 | 0.175 | 9.7E-02 | 0.06 | 0.026 | 1.00 | 0.007 | 0.00 | 0.005 |
| IBS | PTSD | 0.41 | 0.208 | 4.7E-02 | 0.05 | 0.022 | 0.99 | 0.007 | 0.00 | 0.005 |
| IBS | BIP | 0.06 | 0.050 | 2.5E-01 | 0.34 | 0.015 | 1.02 | 0.008 | 0.01 | 0.005 |
| IBS | ASD | 0.24 | 0.068 | 5.0E-04 | 0.20 | 0.015 | 1.00 | 0.008 | 0.00 | 0.005 |
| $IBS_{+}M$ | ADHD | 0.32 | 0.059 | 5.3E-08 | 0.24 | 0.016 | 1.03 | 0.010 | 0.00 | 0.006 |
| $IBS_{+}M$ | SCZ | 0.15 | 0.042 | 4.0E-04 | 0.42 | 0.015 | 1.05 | 0.012 | 0.00 | 0.007 |
| $IBS_{+}M$ | Anxiety | 0.33 | 0.166 | 4.6E-02 | 0.06 | 0.026 | 1.00 | 0.007 | 0.00 | 0.005 |
| $IBS_{+}M$ | PTSD | 0.41 | 0.200 | 4.1E-02 | 0.05 | 0.022 | 0.99 | 0.007 | 0.00 | 0.005 |
| $IBS_{+}M$ | BIP | 0.06 | 0.046 | 2.1E-01 | 0.34 | 0.015 | 1.02 | 0.008 | 0.01 | 0.005 |
| $IBS_{+}M$ | ASD | 0.23 | 0.067 | 6.0E-04 | 0.20 | 0.015 | 1.00 | 0.008 | 0.00 | 0.006 |
| IBD | ADHD | 0.00 | 0.059 | 9.7E-01 | 0.24 | 0.016 | 1.03 | 0.010 | 0.01 | 0.006 |
| IBD | SCZ | 0.01 | 0.039 | 7.2E-01 | 0.42 | 0.015 | 1.05 | 0.012 | 0.00 | 0.006 |
| IBD | Anxiety | 0.28 | 0.156 | 6.7E-02 | 0.06 | 0.026 | 1.00 | 0.007 | 0.00 | 0.004 |
| IBD | PTSD | 0.16 | 0.160 | 3.2E-01 | 0.05 | 0.022 | 0.99 | 0.007 | 0.00 | 0.005 |
| IBD | BIP | 0.01 | 0.054 | 8.7E-01 | 0.34 | 0.015 | 1.02 | 0.008 | 0.00 | 0.005 |
| IBD | ASD | -0.05 | 0.060 | 4.5E-01 | 0.20 | 0.015 | 1.00 | 0.008 | 0.00 | 0.005 |

[1] Cells highlighted with yellow represent that genetic correlation are still significant after Bonferroni correction.

Table S9. LDSC genetic correlation estimates between sensitivity analyses phenotypes and original phenotypes, within sensitivity analyses phenotypes and between sensitivity analyses phenotypes and 6 psychiatric traits (see **Figure S4**)

| **Trait1** | **Trait2** | **rg** | **se** | **p** | **gcov int** | **gcov int se** |
|---|---|---|---|---|---|---|
| GORD (RO) | GORD | 0.99 | 0.004 | 0.0E+00 | 0.940 | 0.008 |
| GORD (RO) | PUD | 0.62 | 0.073 | 1.9E-17 | -0.033 | 0.005 |
| GORD (RO) | IBS | 0.58 | 0.063 | 2.6E-20 | -0.042 | 0.005 |
| GORD (RO) | IBD | 0.09 | 0.068 | 1.9E-01 | -0.030 | 0.005 |
| GORD (RO) | GORD (RO) | 1.00 | 0.000 | 0.0E+00 | 1.014 | 0.008 |
| GORD (RO) | PUD (RO) | 0.42 | 0.081 | 2.4E-07 | -0.041 | 0.005 |
| GORD (RO) | IBS (RO) | 0.47 | 0.077 | 8.5E-10 | -0.047 | 0.005 |
| GORD (RO) | IBD (RO) | -0.03 | 0.074 | 7.0E-01 | -0.029 | 0.005 |
| GORD (RO) | ADHD | 0.49 | 0.046 | 1.1E-25 | -0.002 | 0.006 |
| GORD (RO) | SCZ | -0.01 | 0.029 | 6.3E-01 | 0.013 | 0.006 |
| GORD (RO) | Anxiety | 0.28 | 0.137 | 3.9E-02 | 0.011 | 0.005 |
| GORD (RO) | PTSD | 0.19 | 0.138 | 1.7E-01 | -0.001 | 0.005 |
| GORD (RO) | BIP | -0.07 | 0.038 | 8.2E-02 | 0.001 | 0.006 |
| GORD (RO) | ASD | -0.01 | 0.048 | 8.2E-01 | 0.008 | 0.006 |
| PUD (RO) | GORD | 0.42 | 0.075 | 1.7E-08 | -0.039 | 0.005 |
| PUD (RO) | PUD | 0.93 | 0.026 | 4.1E-279 | 0.843 | 0.006 |
| PUD (RO) | IBS | 0.31 | 0.099 | 1.5E-03 | -0.020 | 0.005 |
| PUD (RO) | IBD | 0.11 | 0.105 | 2.9E-01 | -0.017 | 0.005 |
| PUD (RO) | GORD (RO) | 0.42 | 0.081 | 2.4E-07 | -0.041 | 0.005 |
| PUD (RO) | PUD (RO) | 1.00 | 0.000 | 0.0E+00 | 1.007 | 0.006 |
| PUD (RO) | IBS (RO) | 0.25 | 0.118 | 3.6E-02 | -0.018 | 0.005 |
| PUD (RO) | IBD (RO) | 0.03 | 0.120 | 8.2E-01 | -0.016 | 0.005 |
| PUD (RO) | ADHD | 0.45 | 0.077 | 7.1E-09 | 0.001 | 0.006 |
| PUD (RO) | SCZ | 0.08 | 0.050 | 9.3E-02 | 0.002 | 0.006 |
| PUD (RO) | Anxiety | 0.20 | 0.221 | 3.7E-01 | 0.005 | 0.005 |
| PUD (RO) | PTSD | 0.40 | 0.212 | 5.7E-02 | 0.000 | 0.004 |
| PUD (RO) | BIP | 0.00 | 0.061 | 9.5E-01 | 0.000 | 0.005 |
| PUD (RO) | ASD | 0.03 | 0.075 | 7.0E-01 | 0.000 | 0.005 |
| IBS (RO) | GORD | 0.50 | 0.077 | 7.0E-11 | -0.046 | 0.005 |
| IBS (RO) | PUD | 0.40 | 0.120 | 1.0E-03 | -0.022 | 0.005 |
| IBS (RO) | IBS | 1.00 | 0.017 | 0.0E+00 | 0.873 | 0.007 |
| IBS (RO) | IBD | 0.09 | 0.098 | 3.6E-01 | -0.014 | 0.005 |
| IBS (RO) | GORD (RO) | 0.47 | 0.077 | 8.5E-10 | -0.047 | 0.005 |
| IBS (RO) | PUD (RO) | 0.25 | 0.118 | 3.6E-02 | -0.018 | 0.005 |
| IBS (RO) | IBS (RO) | 1.00 | 0.000 | 0.0E+00 | 1.006 | 0.007 |
| IBS (RO) | IBD (RO) | 0.00 | 0.105 | 9.6E-01 | -0.013 | 0.005 |

| | | | | | | |
|---|---|---|---|---|---|---|
| IBS (RO) | ADHD | 0.26 | 0.073 | 3.0E-04 | 0.001 | 0.006 |
| IBS (RO) | SCZ | 0.16 | 0.055 | 2.7E-03 | -0.006 | 0.007 |
| IBS (RO) | Anxiety | 0.22 | 0.203 | 2.8E-01 | 0.002 | 0.005 |
| IBS (RO) | PTSD | 0.32 | 0.234 | 1.8E-01 | 0.004 | 0.005 |
| IBS (RO) | BIP | 0.06 | 0.056 | 3.0E-01 | 0.006 | 0.005 |
| IBS (RO) | ASD | 0.30 | 0.077 | 1.0E-04 | -0.001 | 0.005 |
| IBD (RO) | GORD | 0.01 | 0.068 | 8.4E-01 | -0.033 | 0.005 |
| IBD (RO) | PUD | 0.05 | 0.110 | 6.6E-01 | -0.020 | 0.006 |
| IBD (RO) | IBS | 0.12 | 0.084 | 1.5E-01 | -0.017 | 0.005 |
| IBD (RO) | IBD | 1.02 | 0.013 | 0.0E+00 | 0.905 | 0.008 |
| IBD (RO) | GORD (RO) | -0.03 | 0.074 | 7.0E-01 | -0.029 | 0.005 |
| IBD (RO) | PUD (RO) | 0.03 | 0.120 | 8.2E-01 | -0.016 | 0.005 |
| IBD (RO) | IBS (RO) | 0.00 | 0.105 | 9.6E-01 | -0.013 | 0.005 |
| IBD (RO) | IBD (RO) | 1.00 | 0.000 | 0.0E+00 | 1.022 | 0.008 |
| IBD (RO) | ADHD | -0.05 | 0.062 | 4.3E-01 | 0.006 | 0.006 |
| IBD (RO) | SCZ | 0.04 | 0.040 | 3.1E-01 | -0.003 | 0.006 |
| IBD (RO) | Anxiety | 0.22 | 0.163 | 1.8E-01 | 0.000 | 0.005 |
| IBD (RO) | PTSD | 0.03 | 0.178 | 8.7E-01 | 0.001 | 0.005 |
| IBD (RO) | BIP | 0.01 | 0.056 | 8.6E-01 | 0.000 | 0.005 |
| IBD (RO) | ASD | -0.07 | 0.066 | 2.9E-01 | 0.003 | 0.005 |

Table S10. Significant SNP-based heritability enrichment for GORD, GP$_+$M, IBS$_+$M and IBD of functional annotation based on the variants within each category after Bonferroni correction (see **Figure S5**)

| Trait | Category | Prop._SNPs | Prop. h2 | Prop. h2 s.e. | Enrichment | Enrichment s.e. | Enrichment p |
|---|---|---|---|---|---|---|---|
| GORD | Conserved | 0.03 | 0.40 | 0.081 | 15.53 | 3.109 | 2.6E-06 |
| GORD | Conserved (extend 500) | 0.33 | 0.66 | 0.073 | 1.98 | 0.220 | 1.3E-05 |
| GORD | DHS (extend 500) | 0.50 | 0.90 | 0.095 | 1.81 | 0.190 | 5.9E-05 |
| GORD | H3K4me1 (extend 500) | 0.61 | 0.91 | 0.068 | 1.49 | 0.112 | 6.1E-05 |
| GORD | H3K4me3 | 0.13 | 0.43 | 0.068 | 3.19 | 0.512 | 1.7E-05 |
| GP$_+$M | Conserved | 0.03 | 0.38 | 0.046 | 14.41 | 1.776 | 9.2E-13 |
| GP$_+$M | Conserved (extend 500) | 0.33 | 0.64 | 0.053 | 1.92 | 0.159 | 2.8E-08 |
| GP$_+$M | DHS (extend 500) | 0.50 | 0.92 | 0.070 | 1.85 | 0.140 | 1.3E-08 |
| GP$_+$M | Fetal DHS | 0.08 | 0.36 | 0.071 | 4.29 | 0.843 | 1.2E-04 |
| GP$_+$M | H3K27ac | 0.39 | 0.52 | 0.031 | 1.32 | 0.080 | 1.1E-04 |
| GP$_+$M | H3K4me1 (extend 500) | 0.61 | 0.90 | 0.043 | 1.47 | 0.070 | 6.0E-10 |
| GP$_+$M | H3K9ac | 0.13 | 0.37 | 0.053 | 2.92 | 0.419 | 4.6E-06 |
| GP$_+$M | H3K9ac (extend 500) | 0.23 | 0.46 | 0.050 | 1.98 | 0.216 | 1.7E-05 |
| GP$_+$M | Intron (extend 500) | 0.40 | 0.48 | 0.020 | 1.20 | 0.051 | 1.1E-04 |
| GP$_+$M | Super Enhancer | 0.17 | 0.25 | 0.019 | 1.51 | 0.112 | 1.1E-05 |
| GP$_+$M | Super Enhancer (extend 500) | 0.17 | 0.27 | 0.018 | 1.60 | 0.106 | 6.9E-08 |
| IBS$_+$M | Conserved | 0.03 | 0.56 | 0.153 | 21.55 | 5.871 | 8.8E-05 |
| IBD | H3K27ac | 0.39 | 0.95 | 0.103 | 2.43 | 0.264 | 6.0E-09 |
| IBD | H3K27ac (extend 500) | 0.42 | 1.04 | 0.110 | 2.47 | 0.260 | 2.2E-08 |
| IBD | Repressed (extend 500) | 0.72 | 0.20 | 0.092 | 0.28 | 0.128 | 1.1E-09 |
| IBD | Super Enhancer | 0.17 | 0.49 | 0.068 | 2.89 | 0.402 | 1.1E-06 |
| IBD | Super Enhancer (extend 500) | 0.17 | 0.59 | 0.062 | 3.43 | 0.362 | 5.7E-11 |

Table S11. SNP-based heritability enrichment for the six digestion phenotypes of cell group functional annotations (see **Figure 3C**)

| Trait | Category | Prop. SNPs | Prop. h2 | Prop. h2 s.e. | Enrichment | Enrichment s.e. | Enrichment p |
|---|---|---|---|---|---|---|---|
| GORD | Adrenal Pancreas | 0.09 | 0.27 | 0.055 | 2.87 | 0.584 | 1.7E-03 |
| GORD[1] | Central Nervous System | 0.15 | 0.41 | 0.053 | 2.75 | 0.355 | 3.9E-06 |
| GORD | Gastrointestinal | 0.17 | 0.29 | 0.067 | 1.73 | 0.397 | 7.1E-02 |
| GORD | Immune | 0.23 | 0.37 | 0.064 | 1.59 | 0.274 | 3.2E-02 |
| GORD | Liver | 0.07 | 0.15 | 0.041 | 2.06 | 0.568 | 6.2E-02 |
| PUD | Adrenal Pancreas | 0.09 | 0.35 | 0.132 | 3.75 | 1.409 | 6.2E-02 |
| PUD | Central Nervous System | 0.15 | 0.59 | 0.144 | 3.94 | 0.970 | 2.4E-03 |
| PUD | Gastrointestinal | 0.17 | 0.58 | 0.161 | 3.43 | 0.960 | 1.5E-02 |
| PUD | Immune | 0.23 | 0.61 | 0.164 | 2.61 | 0.703 | 2.2E-02 |
| PUD | Liver | 0.07 | 0.28 | 0.110 | 3.87 | 1.529 | 5.8E-02 |
| GP+M | Adrenal Pancreas | 0.09 | 0.24 | 0.036 | 2.61 | 0.380 | 3.0E-05 |
| GP+M | Central Nervous System | 0.15 | 0.41 | 0.035 | 2.73 | 0.236 | 3.1E-12 |
| GP+M | Gastrointestinal | 0.17 | 0.26 | 0.040 | 1.57 | 0.239 | 1.9E-02 |
| GP+M | Immune | 0.23 | 0.38 | 0.042 | 1.62 | 0.180 | 6.6E-04 |
| GP+M | Liver | 0.07 | 0.16 | 0.027 | 2.26 | 0.381 | 9.1E-04 |
| IBS | Adrenal Pancreas | 0.09 | 0.35 | 0.138 | 3.78 | 1.470 | 5.5E-02 |
| IBS | Central Nervous System | 0.15 | 0.62 | 0.128 | 4.15 | 0.860 | 1.4E-04 |
| IBS | Gastrointestinal | 0.17 | 0.33 | 0.151 | 1.96 | 0.900 | 3.0E-01 |
| IBS | Immune | 0.23 | 0.46 | 0.159 | 1.96 | 0.683 | 1.6E-01 |
| IBS | Liver | 0.07 | 0.29 | 0.099 | 3.96 | 1.377 | 3.1E-02 |
| IBS+M | Adrenal Pancreas | 0.09 | 0.35 | 0.116 | 3.71 | 1.238 | 2.5E-02 |
| IBS+M | Central Nervous System | 0.15 | 0.61 | 0.108 | 4.09 | 0.723 | 7.2E-06 |
| IBS+M | Gastrointestinal | 0.17 | 0.36 | 0.125 | 2.16 | 0.748 | 1.2E-01 |
| IBS+M | Immune | 0.23 | 0.54 | 0.145 | 2.32 | 0.621 | 3.2E-02 |
| IBS+M | Liver | 0.07 | 0.25 | 0.088 | 3.47 | 1.213 | 4.3E-02 |
| IBD | Adrenal Pancreas | 0.09 | 0.48 | 0.113 | 5.09 | 1.211 | 5.8E-04 |
| IBD | Central Nervous System | 0.15 | 0.35 | 0.118 | 2.35 | 0.791 | 8.6E-02 |
| IBD | Gastrointestinal | 0.17 | 0.72 | 0.142 | 4.28 | 0.850 | 1.0E-04 |
| IBD | Immune | 0.23 | 1.05 | 0.146 | 4.50 | 0.625 | 1.8E-08 |
| IBD | Liver | 0.07 | 0.27 | 0.084 | 3.74 | 1.157 | 1.8E-02 |

[1] Cells highlighted with yellow represent that tissue are still significant after Bonferroni correction.

Table S12. Statistically significant results of partitioning SNP-based heritability of $GP_+M$ and IBD to 205 tissues/cell types using LDSC cell type specific expressed genes analysis (see **Figure 3D**)

| Trait | Name | Tissue category for figures | Coefficient | Coefficient s.e. | Coefficient P value | -log10(p) |
|---|---|---|---|---|---|---|
| IBD | Leukocytes | Blood/Immune | 4.1E-09 | 9.3E-10 | 6.1E-06 | 5.22 |
| IBD | Mononuclear Leukocytes | Blood/Immune | 4.2E-09 | 1.1E-09 | 6.7E-05 | 4.17 |
| IBD | Blood Cells | Blood/Immune | 3.5E-09 | 9.1E-10 | 5.5E-05 | 4.26 |
| $GP_+M$ | Brain Frontal Cortex (BA9) | CNS | 2.1E-09 | 5.7E-10 | 8.0E-05 | 4.10 |

Table S13. Results of partitioning SNP-based heritability of six digestion phenotypes to GTEx brain expression data using LDSC cell type specific expressed genes analysis (see **Figure 3E**)

| Trait | Name | Coefficient | Coefficient s.e. | Coefficient P value |
|---|---|---|---|---|
| GORD[1] | Cortex | 1.8E-09 | 5.7E-10 | 9.2E-04 |
| GORD | Cerebellum | 1.4E-09 | 6.9E-10 | 2.5E-02 |
| GORD | Cerebellar Hemisphere | 1.1E-09 | 6.3E-10 | 4.2E-02 |
| GORD | Frontal Cortex (BA9) | 8.2E-10 | 4.8E-10 | 4.3E-02 |
| GORD | Anterior cingulate cortex (BA24) | 7.2E-10 | 5.0E-10 | 7.4E-02 |
| GORD | Amygdala | 1.3E-11 | 5.7E-10 | 4.9E-01 |
| GORD | Caudate (basal ganglia) | -1.1E-10 | 5.3E-10 | 5.8E-01 |
| GORD | Hippocampus | -2.1E-10 | 5.4E-10 | 6.5E-01 |
| GORD | Putamen (basal ganglia) | -2.3E-10 | 5.4E-10 | 6.7E-01 |
| GORD | Hypothalamus | -2.7E-10 | 5.0E-10 | 7.0E-01 |
| GORD | Nucleus accumbens (basal ganglia) | -4.7E-10 | 4.8E-10 | 8.4E-01 |
| GORD | Substantia nigra | -9.9E-10 | 5.3E-10 | 9.7E-01 |
| GORD | Spinal cord (cervical c-1) | -1.2E-09 | 5.5E-10 | 9.9E-01 |
| GP$_{+}$M | Cortex | 2.5E-09 | 6.9E-10 | 1.2E-04 |
| GP$_{+}$M | Frontal Cortex (BA9) | 2.0E-09 | 6.5E-10 | 8.3E-04 |
| GP$_{+}$M | Cerebellar Hemisphere | 1.4E-09 | 7.1E-10 | 2.4E-02 |
| GP$_{+}$M | Anterior cingulate cortex (BA24) | 1.1E-09 | 6.1E-10 | 3.1E-02 |
| GP$_{+}$M | Cerebellum | 1.1E-09 | 7.3E-10 | 6.7E-02 |
| GP$_{+}$M | Nucleus accumbens (basal ganglia) | 2.6E-10 | 6.2E-10 | 3.4E-01 |
| GP$_{+}$M | Amygdala | -3.2E-11 | 7.1E-10 | 5.2E-01 |
| GP$_{+}$M | Hippocampus | -3.6E-10 | 6.7E-10 | 7.1E-01 |
| GP$_{+}$M | Spinal cord (cervical c-1) | -6.1E-10 | 7.4E-10 | 7.9E-01 |
| GP$_{+}$M | Caudate (basal ganglia) | -5.7E-10 | 6.7E-10 | 8.0E-01 |
| GP$_{+}$M | Hypothalamus | -6.0E-10 | 6.1E-10 | 8.4E-01 |
| GP$_{+}$M | Substantia nigra | -1.4E-09 | 6.6E-10 | 9.8E-01 |
| GP$_{+}$M | Putamen (basal ganglia) | -1.3E-09 | 6.1E-10 | 9.9E-01 |
| IBS$_{+}$M | Frontal Cortex (BA9) | 9.0E-10 | 4.6E-10 | 2.4E-02 |
| IBS$_{+}$M | Cortex | 7.9E-10 | 5.1E-10 | 5.9E-02 |
| IBS$_{+}$M | Cerebellum | 7.2E-10 | 5.3E-10 | 8.9E-02 |
| IBS$_{+}$M | Cerebellar Hemisphere | 5.8E-10 | 4.9E-10 | 1.2E-01 |
| IBS$_{+}$M | Amygdala | 4.7E-10 | 4.5E-10 | 1.5E-01 |
| IBS$_{+}$M | Anterior cingulate cortex (BA24) | 4.1E-10 | 4.5E-10 | 1.8E-01 |
| IBS$_{+}$M | Spinal cord (cervical c-1) | 1.2E-10 | 4.1E-10 | 3.9E-01 |
| IBS$_{+}$M | Putamen (basal ganglia) | 1.3E-10 | 4.7E-10 | 3.9E-01 |
| IBS$_{+}$M | Hippocampus | 3.4E-11 | 4.3E-10 | 4.7E-01 |
| IBS$_{+}$M | Substantia nigra | -3.9E-11 | 4.2E-10 | 5.4E-01 |
| IBS$_{+}$M | Caudate (basal ganglia) | -3.0E-10 | 4.5E-10 | 7.5E-01 |
| IBS$_{+}$M | Nucleus accumbens (basal ganglia) | -3.4E-10 | 4.4E-10 | 7.8E-01 |
| IBS$_{+}$M | Hypothalamus | -1.2E-09 | 4.7E-10 | 9.9E-01 |

[1] Cells highlighted with yellow represent that tissue are still significant after Bonferroni correction.

Table S14. Results[1] of SMR analysis for the 3 digestion traits using GTEx and eQTLGen data

| eQTL Data | Trait | Tissue | Probe ID | Probe Chr. | Probe base pair position | Gene | Top SNP | A1/ A2 | Freq | $P_{GWAS}$ | $P_{eQTL}$ | $b_{SMR}$ | $P_{SMR}$ | $P_{HEIDI}$ | HEIDI pass |
|---|---|---|---|---|---|---|---|---|---|---|---|---|---|---|---|
| GTEx | PUD | Stomach | ENSG00000 167653.4 | 8 | 143757934 | *PSCA* | rs2976388 | A/G | 0.42 | 7.4E-11 | 8.8E-41 | -0.12 | 4.8E-09 | 0.49 | Yes |
| | | | ENSG00000 176920.10 | 19 | 49204217 | *FUT2* | rs601338 | A/G | 0.51 | 8.8E-13 | 2.8E-20 | -0.33 | 1.6E-08 | 0.83 | Yes |
| | IBD | Colon Sigmoid | ENSG00000 257582.1 | 10 | 101288520 | *RP11-129J12.2* | rs7085798 | A/C | 0.52 | 7.8E-11 | 2.1E-20 | 0.18 | 1.0E-07 | 0.05 | Yes |
| | | Colon Transverse | ENSG00000 257582.1 | 10 | 101288520 | *RP11-129J12.2* | rs1548962 | C/G | 0.52 | 5.4E-11 | 7.0E-22 | 0.22 | 6.0E-08 | 0.16 | Yes |
| eQTL Gen | PUD | Whole blood | ENSG00000 142233 | 19 | 49170501 | *NTN5* | rs569970 | T/C | 0.47 | 7.7E-11 | 4.4E-193 | 0.35 | 2.1E-10 | 0.01 | No |
| | GP$_{+}$M | | ENSG00000 139531 | 12 | 56395694 | *SUOX* | rs1873914 | C/G | 0.42 | 2.0E-07 | 0 | 0.08 | 2.4E-07 | 0.40 | Yes |
| | IBD | | ENSG00000 170128 | 1 | 200842694 | *GPR25* | rs296545 | A/G | 0.30 | 4.6E-09 | 4.2E-58 | -0.81 | 3.7E-08 | 0.0002 | No |
| | | | ENSG00000 233276 | 3 | 49395321 | *GPX1* | rs11130203 | A/G | 0.31 | 7.4E-09 | 4.8E-35 | -0.99 | 1.6E-07 | 0.01 | Yes |
| | | | ENSG00000 164062 | 3 | 49716415 | *APEH* | rs11718165 | G/A | 0.30 | 3.4E-08 | 5.9E-231 | 0.36 | 5.3E-08 | 0.05 | Yes |
| | | | ENSG00000 187796 | 9 | 139262244 | *CARD9* | rs11794847 | A/G | 0.42 | 2.9E-07 | 0 | 0.17 | 3.1E-07 | 0.38 | Yes |

[1] All listed results are significant after Bonferroni correction at $P < 0.05/155,059$.

Table S15. Mendelian Randomisation analysis results of investigating the causality hypothesis between major depression and six digestion phenotypes (see **Figure 4A**)

| Trait1 | Trait2 | Direction Trait1 -> Trait 2[1] | | | | Direction Trait2 -> Trait 1[2] | | | | |
|---|---|---|---|---|---|---|---|---|---|---|
| | | $b_{xy}$[3] | se | p | No. of SNPs | $b_{xy}$[3] | se | p | No. of SNPs | Significance threshold |
| Major depression | GORD | 0.22 | 0.042 | 1.4E-07 | 33 | 0.15[4] | 0.036 | 3.4E-05 | 12 | 5.0E-07 |
| Major depression | PUD | 0.18 | 0.073 | 1.0E-02 | 33 | 0.03 | 0.022 | 1.8E-01 | 11 | 1.0E-06 |
| Major depression | GP+M | 0.23 | 0.032 | 1.1E-12 | 33 | 0.15 | 0.038 | 8.0E-05 | 17 | 5.0E-08 |
| Major depression | IBS | 0.30 | 0.067 | 6.2E-06 | 33 | 0.06 | 0.026 | 2.4E-02 | 11 | 2.0E-06 |
| Major depression | IBS+M | 0.31 | 0.064 | 2.0E-06 | 33 | 0.04 | 0.027 | 1.5E-01 | 10 | 2.0E-06 |
| Major depression | IBD | 0.05 | 0.103 | 6.0E-01 | 33 | 0.003 | 0.008 | 7.4E-01 | 25 | 5.0E-08 |

[1] The direction represents using trait 1 as exposure to investigate the causality hypothesis on trait 2.
[2] The direction represents using trait 2 as exposure to investigate the causality hypothesis on trait 1.
[3] The unit represents per standard deviation change in liability to the exposure trait.
[4] Yellow highlighted cells indicate use of a relaxed significance threshold for genetic instrument inclusion in the Trait2 -> Trait1 analysis, specified in the significance threshold column.

Table S16. MR results for the relationship between major depression and GP$_{+}$M

| Exposure | Outcome | Method | No. of SNPs | b | se | P | Egger intercept (se) | Egger intercept P |
|---|---|---|---|---|---|---|---|---|
| MD | GP$_{+}$M | Inverse variance weighted[1] | 28 | 0.22[2] | 0.04 | 8.9E-7 | - | - |
| MD | GP$_{+}$M | MR Egger[1] | 28 | 0.05[2] | 0.18 | 7.9E-1 | 0.006 (0.006) | 3.4E-1 |
| MD | GP$_{+}$M | Weighted median[1] | 28 | 0.17[2] | 0.05 | 1.6E-3 | - | - |
| GP$_{+}$M | MD | Inverse variance weighted[1] | 14 | 0.12[2] | 0.05 | 9.8E-3 | - | - |
| GP$_{+}$M | MD | MR Egger[1] | 14 | 0.32[2] | 0.36 | 3.9E-1 | -0.007 (0.013) | 6.0E-1 |
| GP$_{+}$M | MD | Weighted median[1] | 14 | 0.11[2] | 0.06 | 5.0E-2 | - | - |

[1] No MR-PRESSO (Pleiotropy Residual Sum and Outlier) outliers were detected.

[2] The beta is the original value from the corresponding MR analysis.

Abbreviation: MR: mendelian randomization; GP$_{+}$M: phenotype for gastro-oesophageal reflux disease, peptic ulcer disease with the combination of taking corresponding medications.

Table S17. Genetic risk score for major depression[1] predicts $GP_{+}M$ (gastro-oesophageal reflux disease (GORD), peptic ulcer disease (PUD) in combination with medications for GORD and PUD) risk

| Clumping Range | Sample Prevalence (P) | Sample Size (N) | P value of case-control GRS difference | AUC | $ORD_{10}$[2] | $ORD_{10}$ CI[3] |
|---|---|---|---|---|---|---|
| 0-5.0E-08 | 0.165 | 456414 | 1.1E-11 | 0.508 | 1.08 | 1.04-1.12 |
| 0-1.0E-05 | 0.165 | 456414 | 1.1E-38 | 0.514 | 1.19 | 1.15-1.24 |
| 0-1.0E-04 | 0.165 | 456414 | 3.1E-59 | 0.518 | 1.28 | 1.23-1.32 |
| 0-1.0E-03 | 0.165 | 456414 | 2.0E-82 | 0.522 | 1.31 | 1.27-1.36 |
| 0-1.0E-02 | 0.165 | 456414 | 4.8E-109 | 0.525 | 1.34 | 1.29-1.39 |
| 0-5.0E-02 | 0.165 | 456414 | 3.4E-78 | 0.522 | 1.31 | 1.27-1.36 |
| 0-1.0E-01 | 0.165 | 456414 | 1.6E-82 | 0.522 | 1.29 | 1.25-1.34 |
| 0-5.0E-01 | 0.165 | 456414 | 1.0E-77 | 0.522 | 1.28 | 1.23-1.32 |

[1] The GWAS summary statistics data for major depression are from Wray *et al*.[9] (European ancestry and UK Biobank participants were excluded).

[2] The odds ratio of $GP_{+}M$ risk for participants with genetic risk score at 10$^{th}$ decile compared with participants with genetic risk score at 1$^{st}$ decile.

[3] 95% confidence interval for the odds ratio at 10$^{th}$ decile.

### *Supplementary Figures*

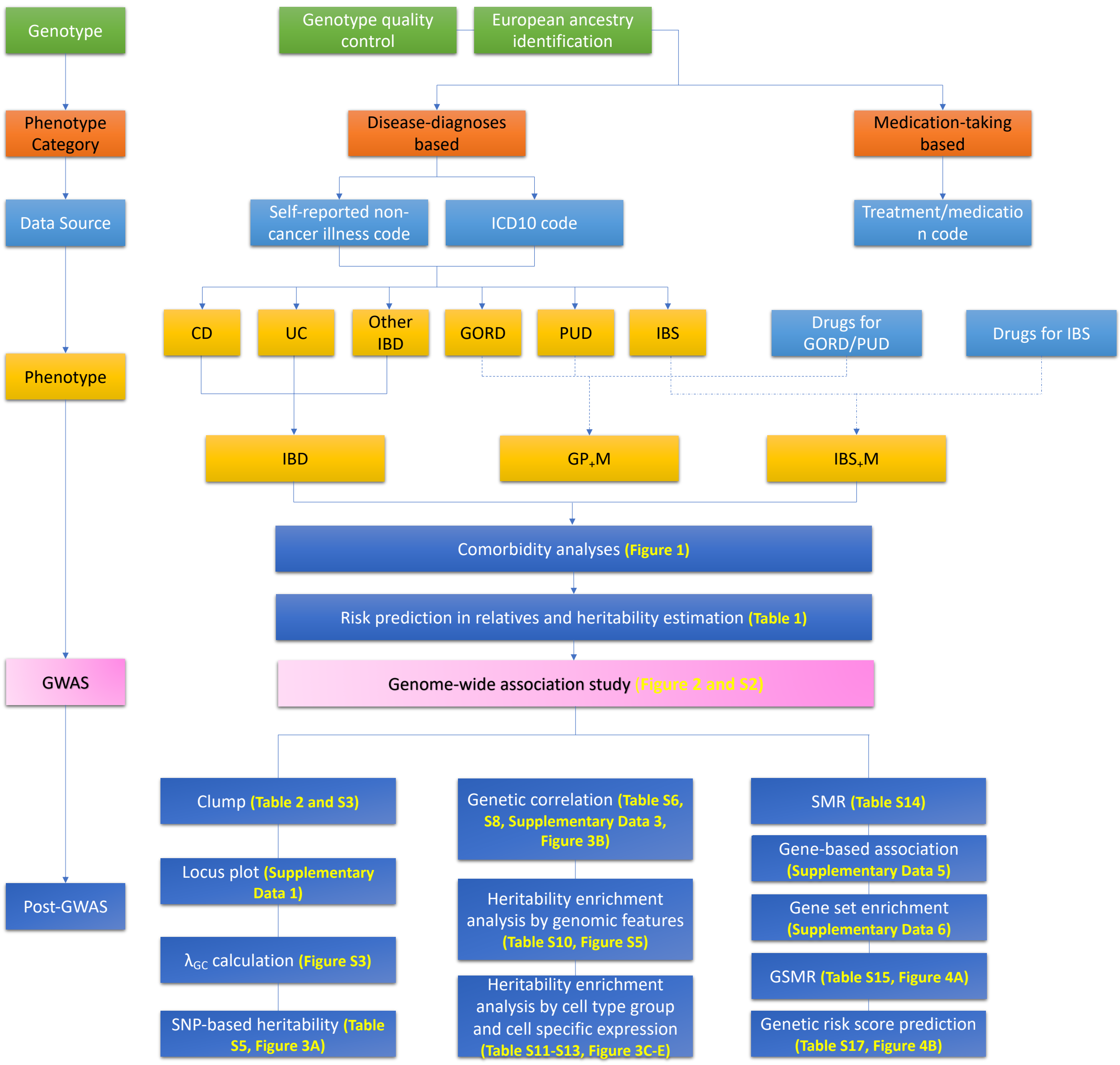

**Figure S1**. Full workflow of the study.

#### A IBS

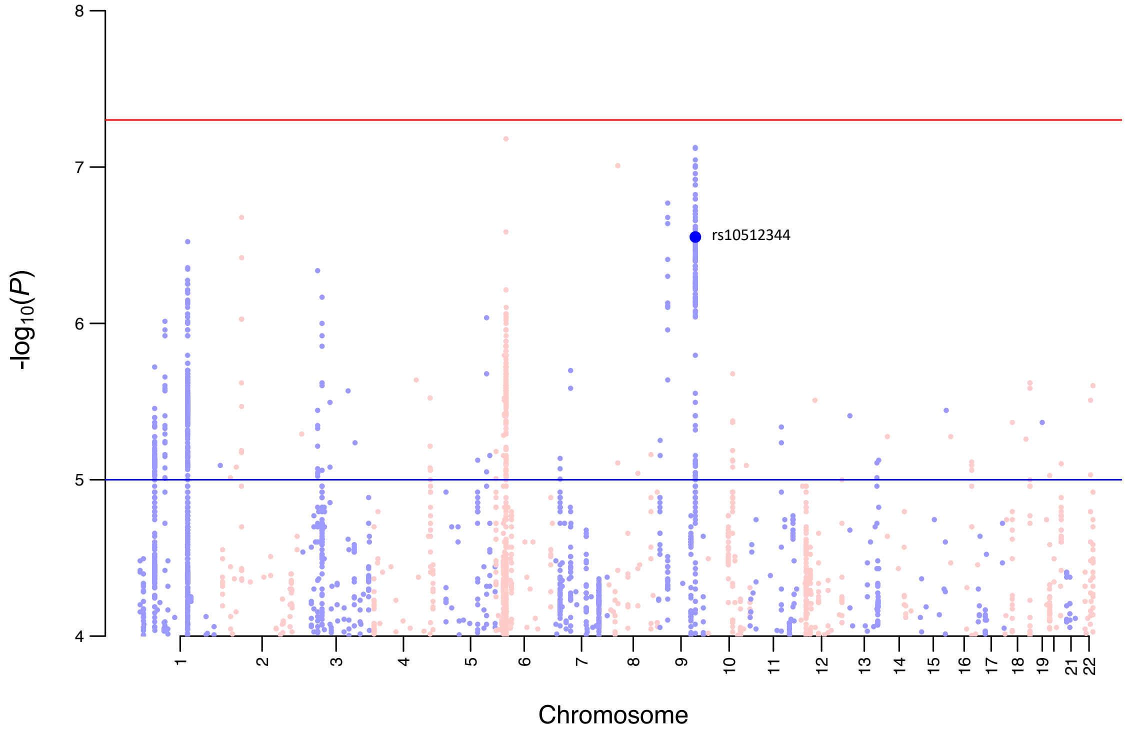

#### B IBD

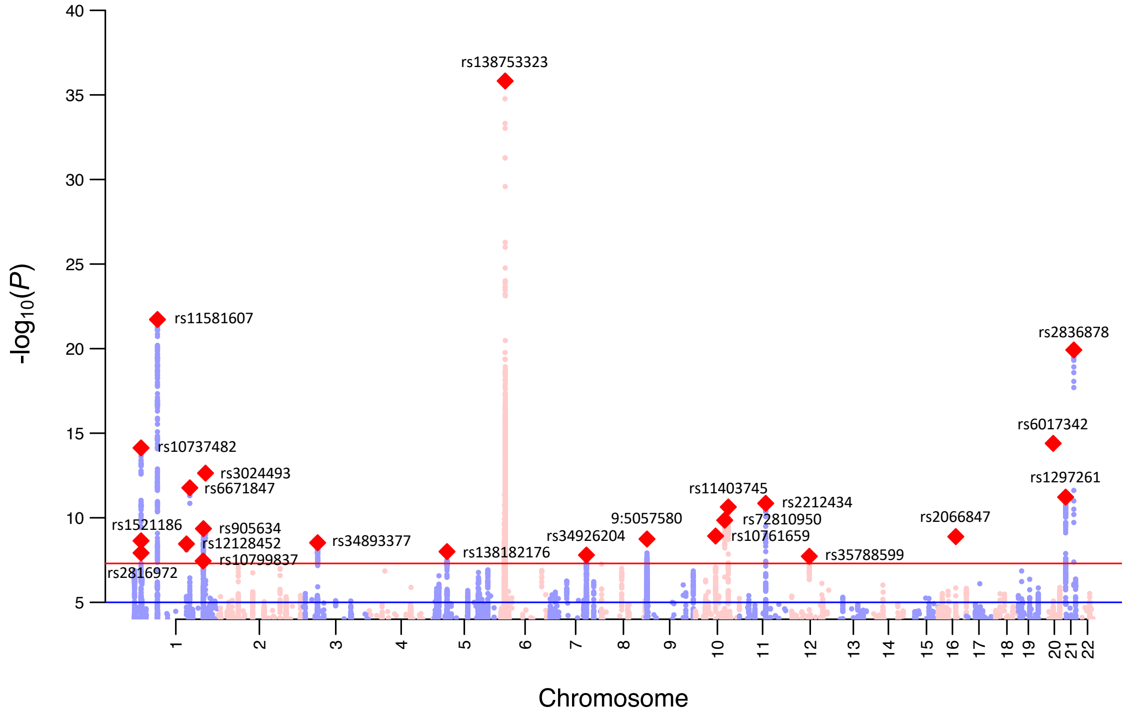

**Figure S2**. Manhattan plots for IBS and IBD. SNPs with red diamond represent genome-wide statistically significant independent loci (P < 5.0E-8) for each trait. For IBS (Panel A), rs10512344 (dot highlighted with blue colour) has been reported as associated with female IBS in UKB[6].

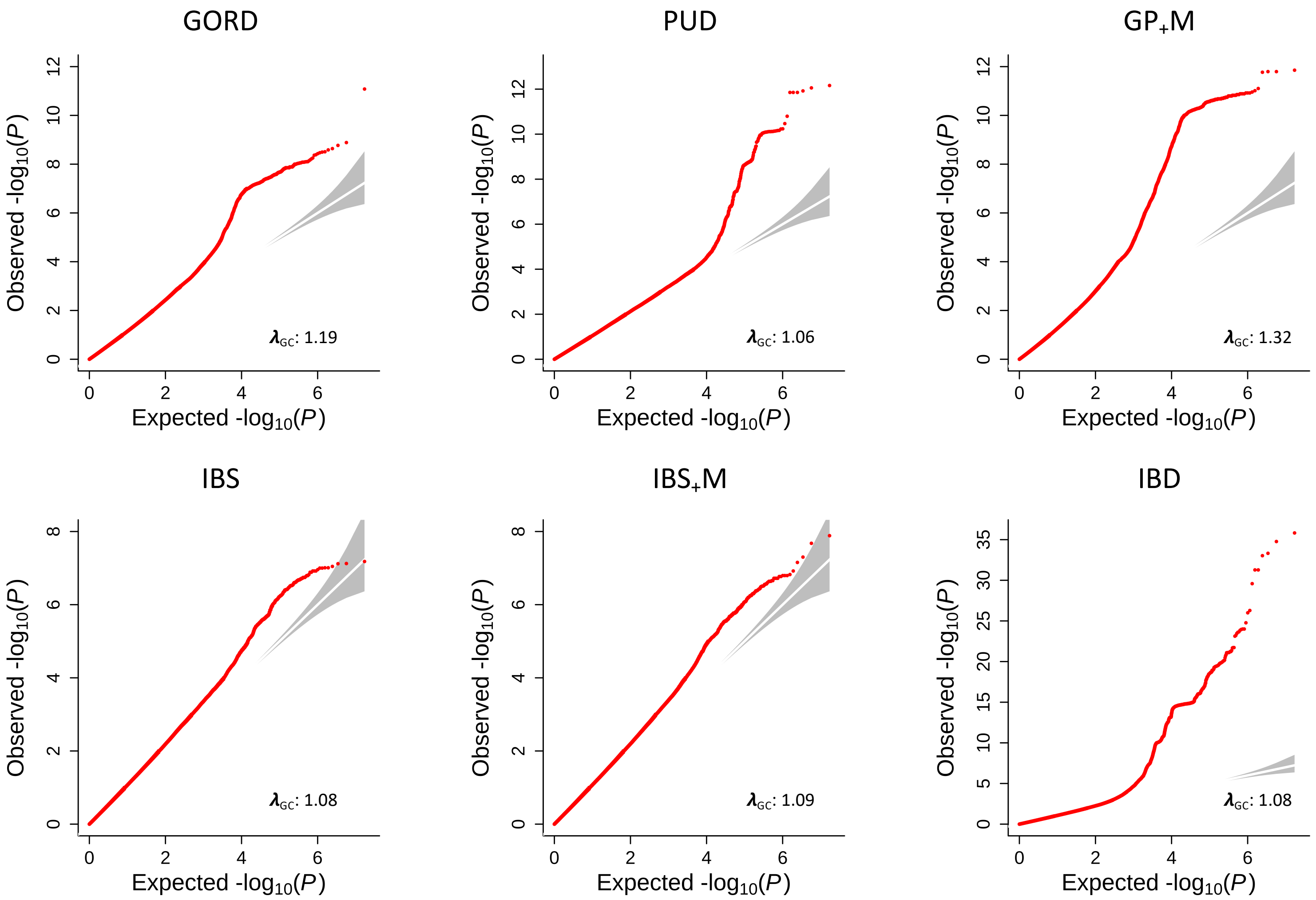

**Figure S3**. Quantile-Quantile (Q-Q) plots for the six digestion phenotypes.

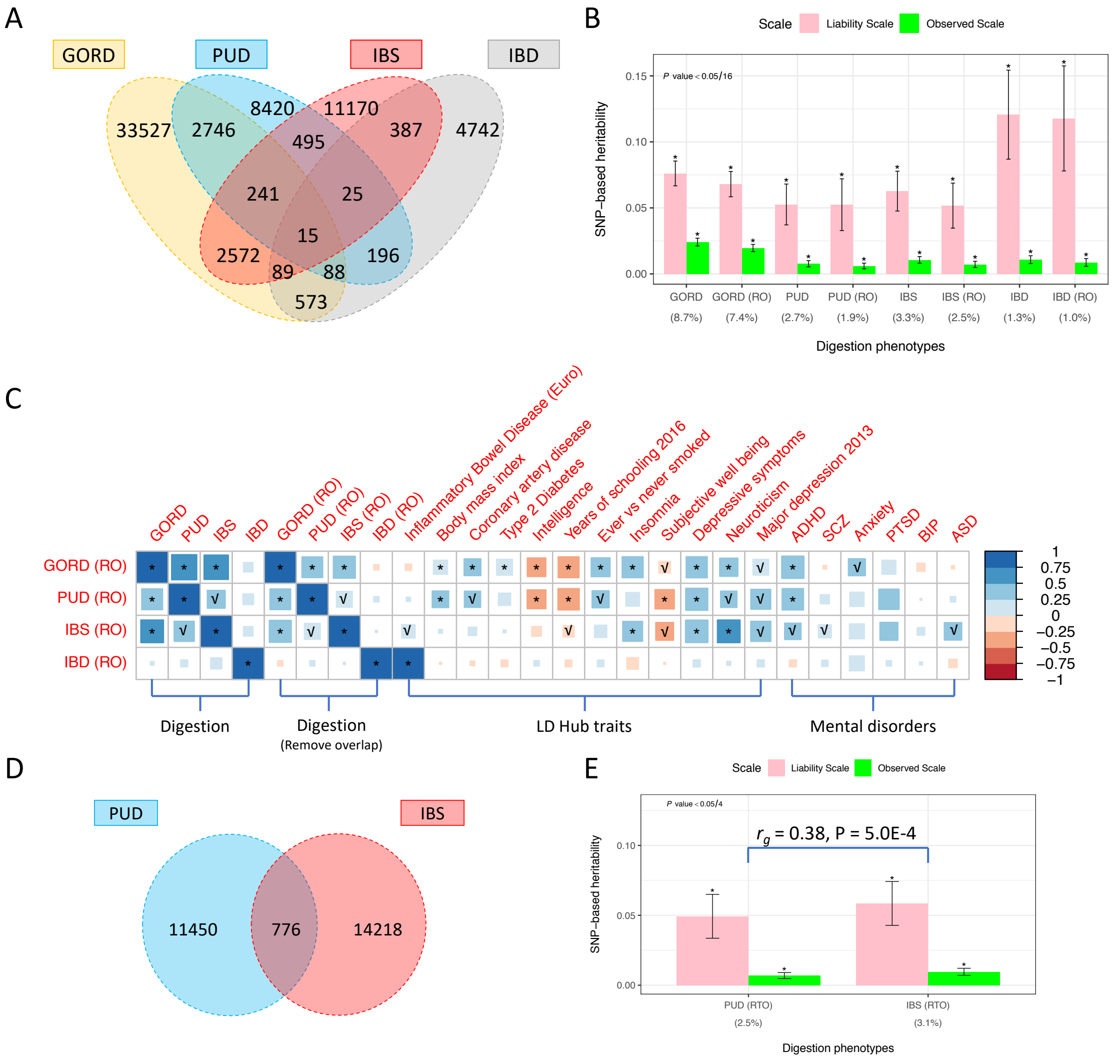

**Figure S4**. SNP-based heritability and genetic correlation for GORD, PUD, IBS and IBD after removing the overlapped individuals. **Panel A**. Venn diagram for the number of overlapped individuals among GORD, PUD, IBS and IBD cases. **Panel B**. Comparison of SNP-based heritability on observed and liability scale for GORD, PUD, IBS and IBD between the original phenotypes and phenotypes generated after removing individuals with more than one disorder (defined as sensitivity analysis phenotypes). “RO” represents removing overlapped individuals with more than one disorder. We took sample risk, i.e. the proportion cases in the UKB cohort, as the population lifetime risk to calculate the SNP-based heritability on the liability scale for each digestion phenotype; the sample risk percentage is shown below x axis. **Panel C**. Genetic correlation between sensitivity analysis phenotypes with the original phenotypes, within sensitivity analysis phenotypes, between sensitivity analysis phenotypes with traits from LD Hub and six published mental disorder studies. “*” represents that genetic correlation estimate was still significant after Bonferroni correction (P < 0.05/(4*4+4*4+4*258+4*6)) while “√” represents the P value for genetic correlation estimates < 0.05. **Panel D**. The number of overlapped individuals between PUD and IBS cases. **Panel E**. SNP-based heritability on observed and liability scale for PUD and IBS phenotypes generated after removing the 776 individuals with both PUD and IBS. “RTO” represents removing the 776 individuals with both PUD and IBS. The sample risk percentage, as shown below axis, was used as the population lifetime risk to calculate the SNP-based heritability on the liability scale.

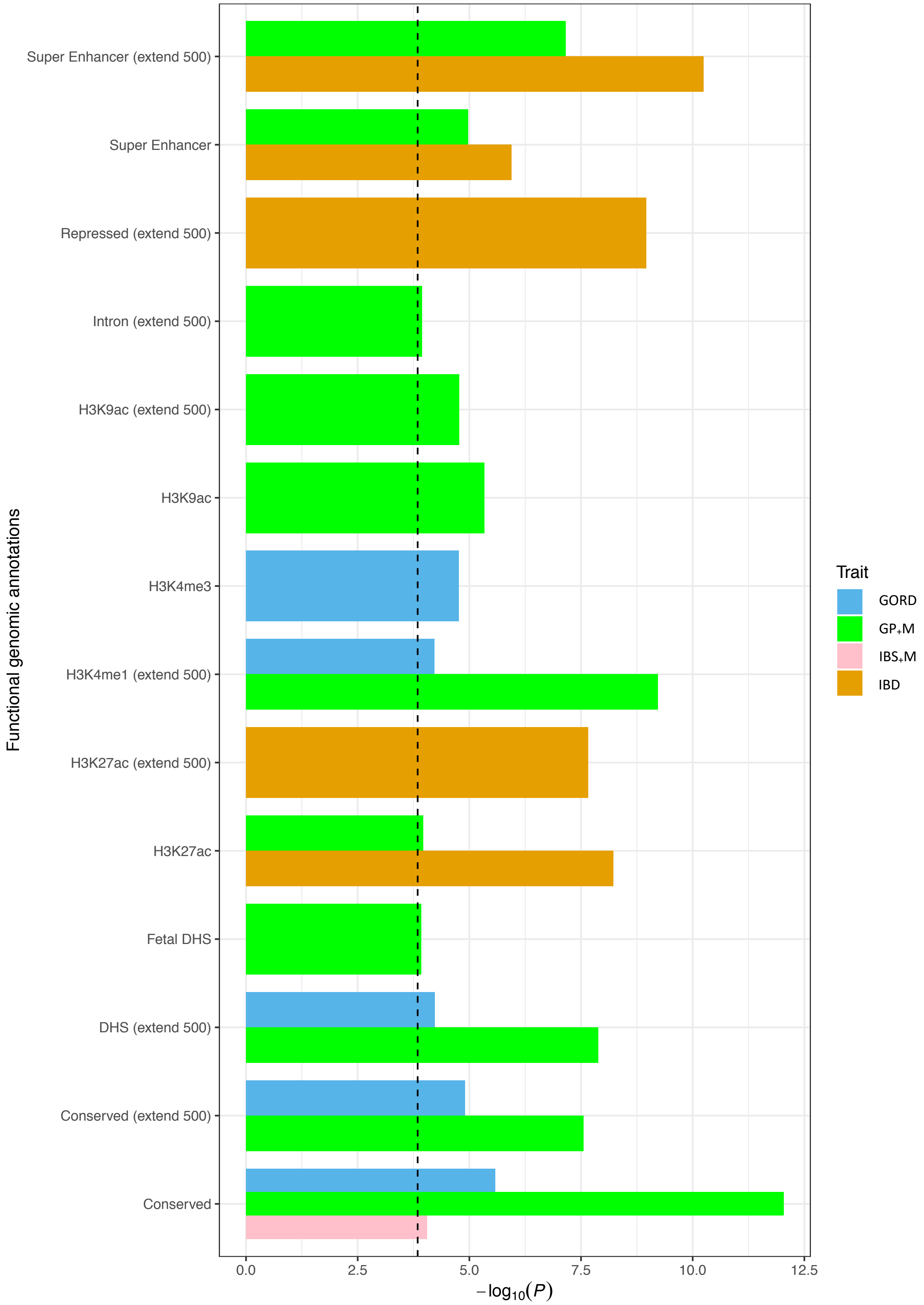

**Figure S5**. Significant heritability enrichment for GORD, GP+M, IBS+M and IBD of functional annotation based on the variants within each category after Bonferroni correction (P < 0.05/(53*6+5*6)).

| Category | Group | Phenotype | Depression phenotype | GORD | | PUD | | IBS | | IBD | | |
|---|---|---|---|---|---|---|---|---|---|---|---|---|
| Minimal phenotyping | Help seeking | Gppsy | Cases: 118,986 | Cases: 29,363 | Controls: 316,659 | Cases: 8,951 | Controls: 337,071 | Cases: 11,218 | Controls: 334,804 | Cases: 4,570 | | Controls: 341,452 |
| | | | Controls: 227,036 | 13,241 | 105,745 | 3,795 | 115,191 | 6,461 | 112,525 | 1,639 | *N* | 117,347 |
| | | | | 16,122 | 210,914 | 5,156 | 221,880 | 4,757 | 222,279 | 2,931 | | 224,105 |
| | | Psypsy | Cases: 40,270 | Cases: 29,455 | Controls: 317,404 | Cases: 8,974 | Controls: 337,885 | Cases: 11,242 | Controls: 335,617 | Cases: 4,575 | | Controls: 342,284 |
| | | | Controls: 306,589 | 4,850 | 35,420 | 1,538 | 38,732 | 2,400 | 37,870 | 596 | *N* | 39,674 |
| | | | | 24,605 | 281,984 | 7,436 | 299,153 | 8,842 | 297,747 | 3,979 | | 302,610 |
| | Cardinal Symptom | DepAll | Cases: 22,418 | Cases: 6,851 | Controls: 76,669 | Cases: 1,948 | Controls: 81,572 | Cases: 2,275 | Controls: 81,245 | Cases: 982 | | Controls: 82,538 |
| | | | Controls: 61,102 | 2,358 | 20,060 | 641 | 21,777 | 990 | 21,428 | 271 | *N* | 22,147 |
| | | | | 4,493 | 56,609 | 1,307 | 59,795 | 1,285 | 59,817 | 711 | | 60,391 |
| | Self-report | SelfRepDep | Cases: 19,919 | Cases: 27,726 | Controls: 232,613 | Cases: 8,459 | Controls: 251,880 | Cases: 10,955 | Controls: 249,384 | Cases: 4,416 | | Controls: 255,923 |
| | | | Controls: 240,420 | 2,788 | 17,131 | 808 | 19,111 | 1,472 | 18,447 | 292 | *N* | 19,627 |
| | | | | 24,938 | 215,482 | 7,651 | 232,769 | 9,483 | 230,937 | 4,124 | | 236,296 |
| EMR | EMR | ICD10Dep | Cases: 9,949 | Cases: 28,298 | Controls: 242,222 | Cases: 8,656 | Controls: 261,864 | Cases: 10,397 | Controls: 260,123 | Cases: 4,437 | | Controls: 266,083 |
| | | | Controls: 260,571 | 1,968 | 7,981 | 637 | 9,312 | 909 | 9,040 | 232 | | 9,717 |
| | | | | 26,330 | 234,241 | 8,019 | 252,552 | 9,488 | 251,083 | 4,205 | | 256,366 |
| Strictly defined | DSM V | LifetimeMDD | Cases: 16,946 | Cases: 4,547 | Controls: 62,274 | Cases: 1,234 | Controls: 65,587 | Cases: 1,765 | Controls: 65,056 | Cases: 754 | | Controls: 66,067 |
| | | | Controls: 49,875 | 1,520 | 15,426 | 385 | 16,561 | 826 | 16,120 | 237 | | 16,709 |
| | | | | 3,027 | 46,848 | 849 | 49,026 | 939 | 48,936 | 517 | | 49,358 |
| | | MDDRecur | Cases: 10,217 | Cases: 3,995 | Controls: 56,004 | Cases: 1,088 | Controls: 58,911 | Cases: 1,502 | Controls: 58,497 | Cases: 669 | | Controls: 59,330 |
| | | | Controls: 49,782 | 975 | 9,242 | 244 | 9,973 | 567 | 9,650 | 153 | | 10,064 |
| | | | | 3,020 | 46,762 | 844 | 48,938 | 935 | 48,847 | 516 | | 49,266 |
| No MDD | No Symptom | GpNoDepr | Cases: 9,066 | Cases: 4,468 | Controls: 56,339 | Cases: 1,299 | Controls: 59,508 | Cases: 1,273 | Controls: 59,534 | Cases: 705 | | Controls: 60,102 |
| | | | Controls: 51,741 | 965 | 8,101 | 272 | 8,794 | 389 | 8,677 | 107 | *N* | 8,959 |
| | | | | 3,503 | 48,238 | 1,027 | 50,714 | 884 | 50,857 | 598 | | 51,143 |

**Figure S6.** Comorbidity relationship between eight depression phenotypes and each of the GORD, PUD, IBS and IBD. Eight depression phenotypes were derived according to Cai *el al.*[10] and are grouped into four categories: minimal phenotyping, EMR (electronic medical record), strictly defined major depression disorder (MDD) and No MDD. The corresponding case and control number of the eight depression phenotypes are on the left of orange dashed line. For each of the GORD, PUD, IBS and IBD, the corresponding overlapped case and control number with eight depression phenotypes were calculated, as shown above the blue dashed line. We also listed the number of individuals who were cases for both depression and digestion phenotypes, only one phenotype and neither of them, as shown below the corresponding overlapped case and control number. The "*N*" represents no statistical significance after Bonferroni correction ($P < 0.05/(6+4*8)$). Abbreviation: Seen general practice (GP) for nerves, anxiety, tension or depression (Gppsy); Seen psychiatrist for nerves, anxiety, tension or depression (Psypsy); Probable recurrent major depression or single probable major depression episode (DepAll); Self-reported depression (SelfRepDep); ICD10 defined depression (ICD10Dep); DSM-V clinical guideline defined major depression (LifetimeMDD); Major depression recurrence (MDDRecur); Seen GP for depression but no cardinal symptoms (GpNoDep); Gastro-oesophageal reflux disease (GORD); Peptic ulcer disease (PUD); Irritable bowel syndrome (IBS) and Inflammatory bowel disease (IBD).

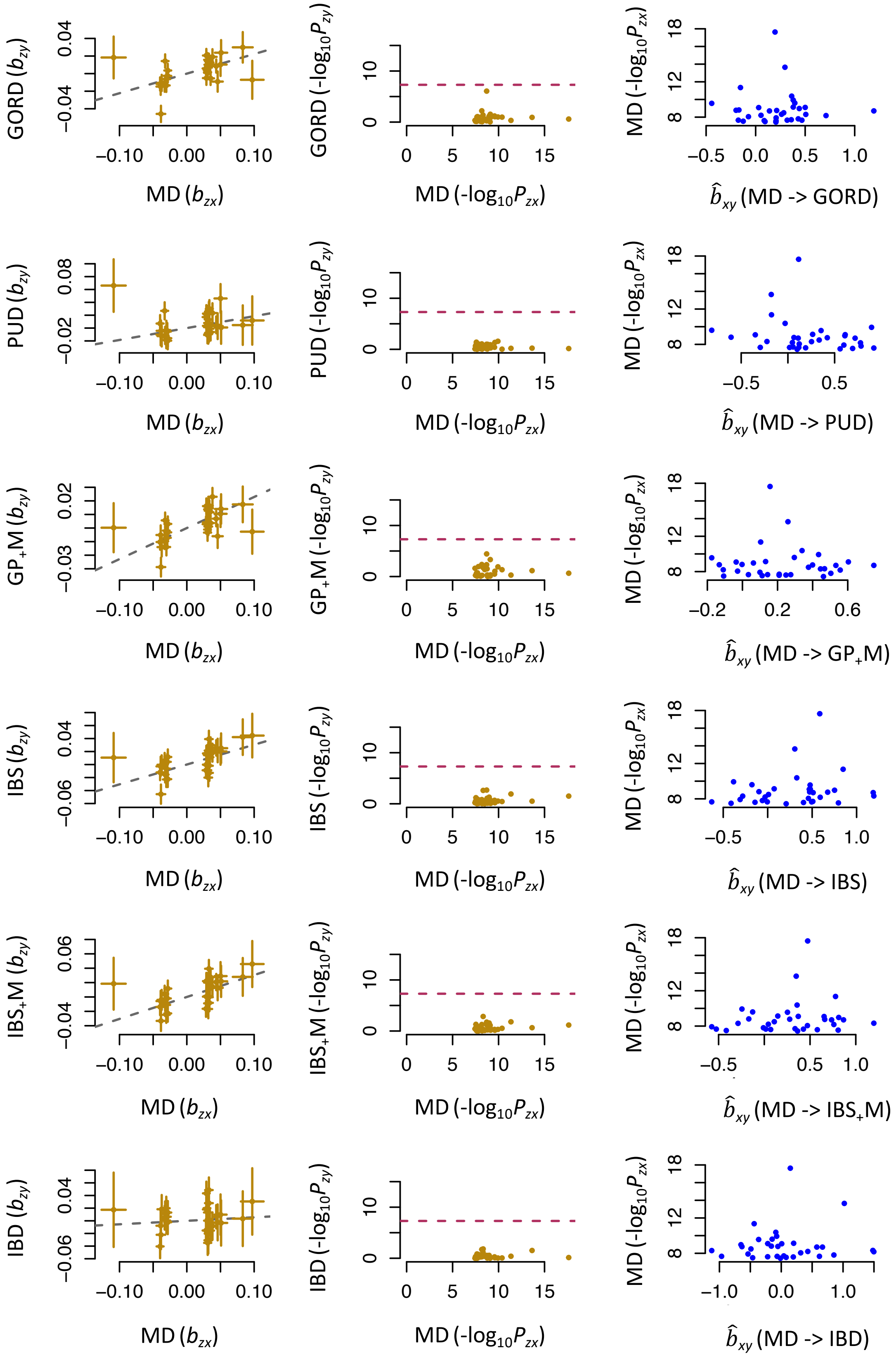

**Figure S7**. Plots of the effect sizes of exposure SNPs (x-axis) vs. outcome SNPs (y-axis) (first column) and association P values of all the genetic instruments from GWAS for major depression (MD) vs. those for each of the six digestion phenotypes (second column). Shown in the third column are the plots of $b_{xy}$ vs. GWAS P value of MD at each genetic variant. All the exposure SNPs were obtained after HEIDI outlier test.

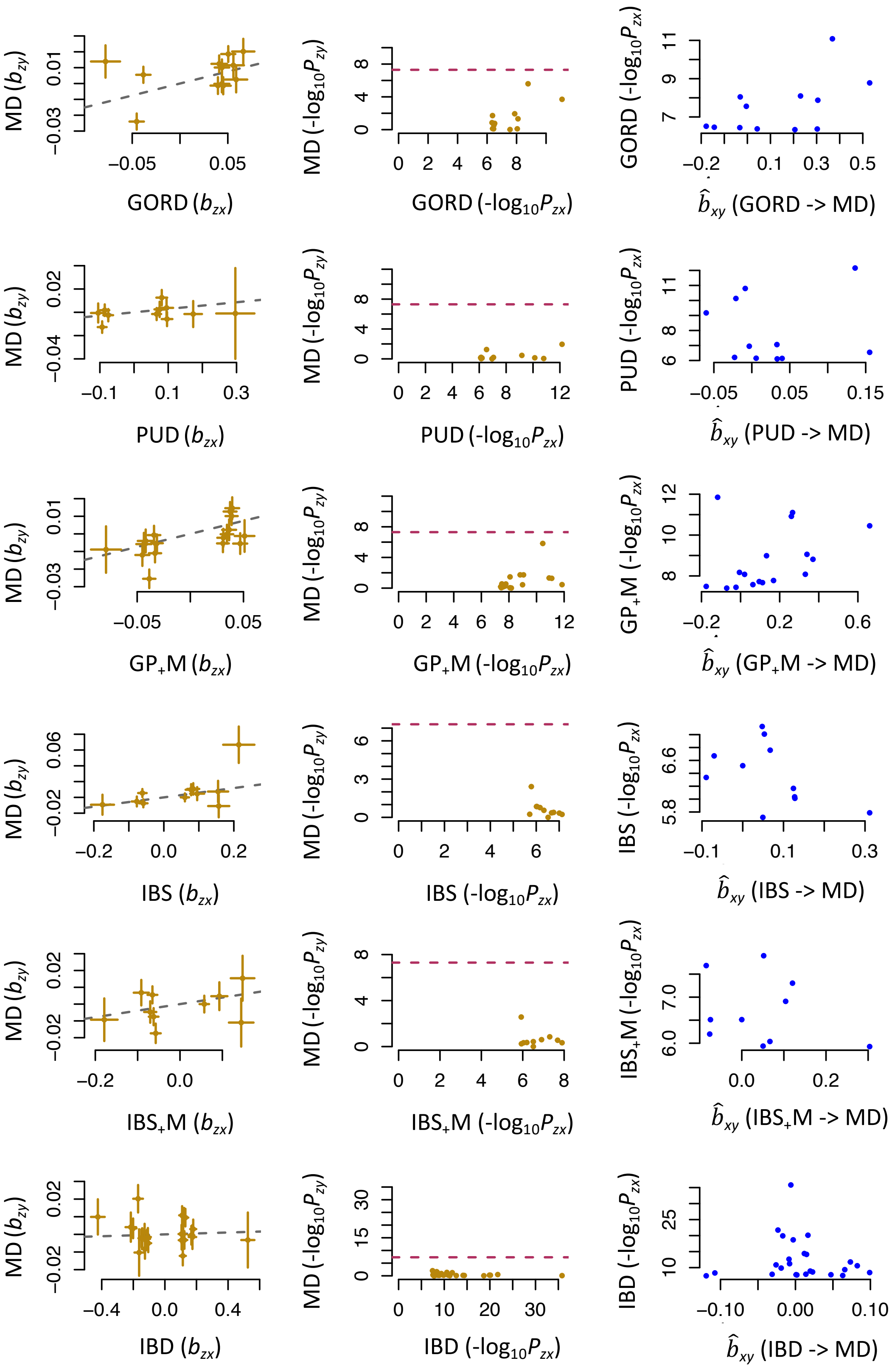

**Figure S8**. Plots of the effect sizes of exposure SNPs (x-axis) vs. outcome SNPs (y-axis) (first column) and association P values of all the genetic instruments from GWAS for each of the six digestion phenotypes vs. those for major depression (MD) (second column). Shown in the third column are the plots of $b_{xy}$ vs. GWAS P value of the six digestion phenotypes at each genetic variant. All the exposure SNPs were obtained after HEIDI outlier test. For GORD, PUD, IBS and IBS$_+$M, we relaxed the significance threshold to obtain more SNP instruments.

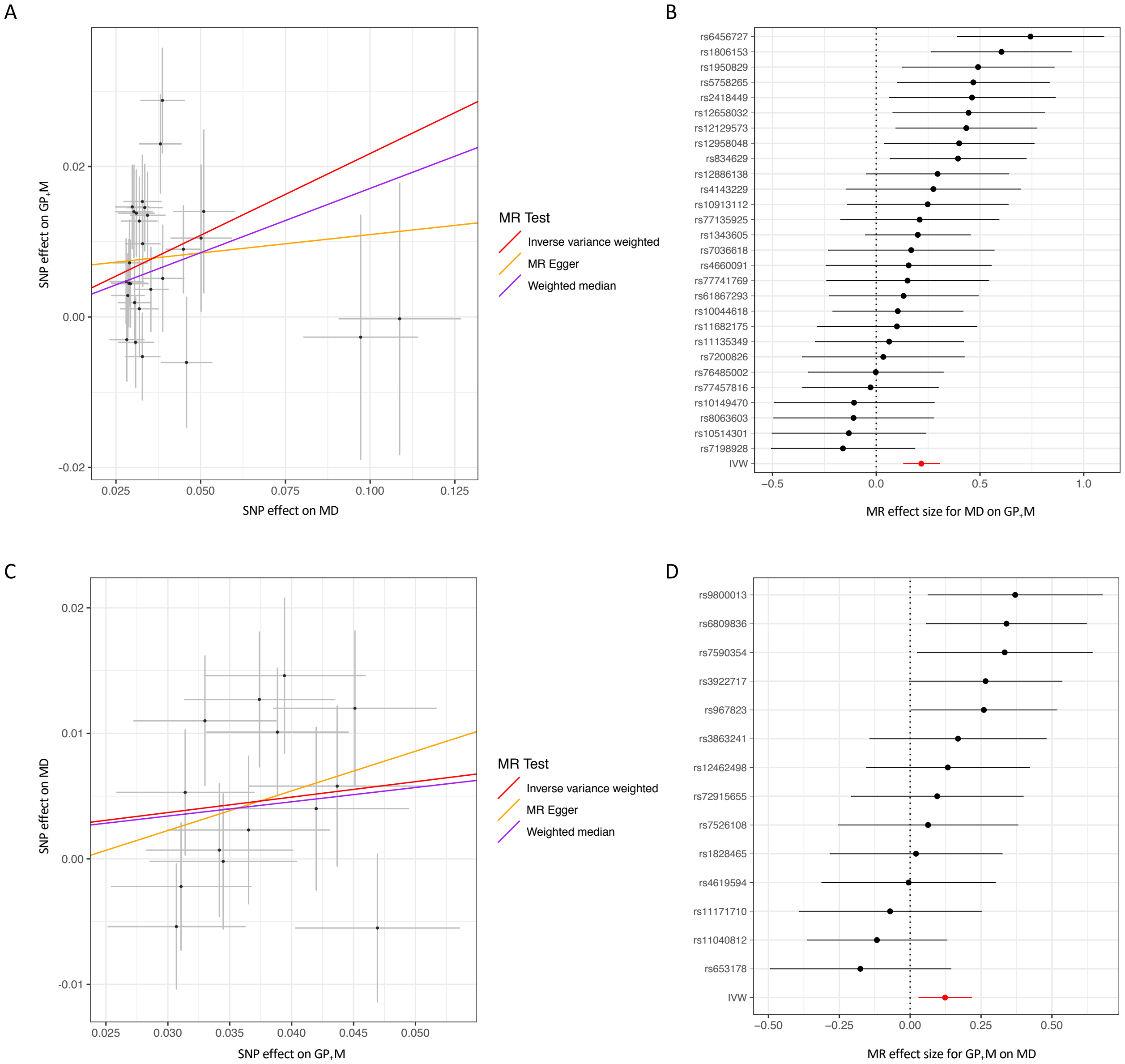

**Figure S9.** Mendelian Randomization (MR) plots between major depression and GP$_+$M. **Panel A.** Scatterplot of single-nucleotide polymorphism (SNP) potential effects of major depression on GP$_+$M, with the slope of each line corresponding to estimated MR effect per method. **Panel B.** Forest plot of individual and combined SNP MR-estimated effect sizes of major depression on GP$_+$M. The raw effect sizes with 95% confidence interval are presented as the dot and horizontal line. **Panel C.** Scatterplot of SNP potential effects of GP$_+$M on major depression, with the slope of each line corresponding to estimated MR effect per method. **Panel D.** Forest plot of individual and combined SNP MR-estimated effect sizes of GP$_+$M on major depression. The raw effect sizes with 95% confidence interval are presented as the dot and horizontal line.

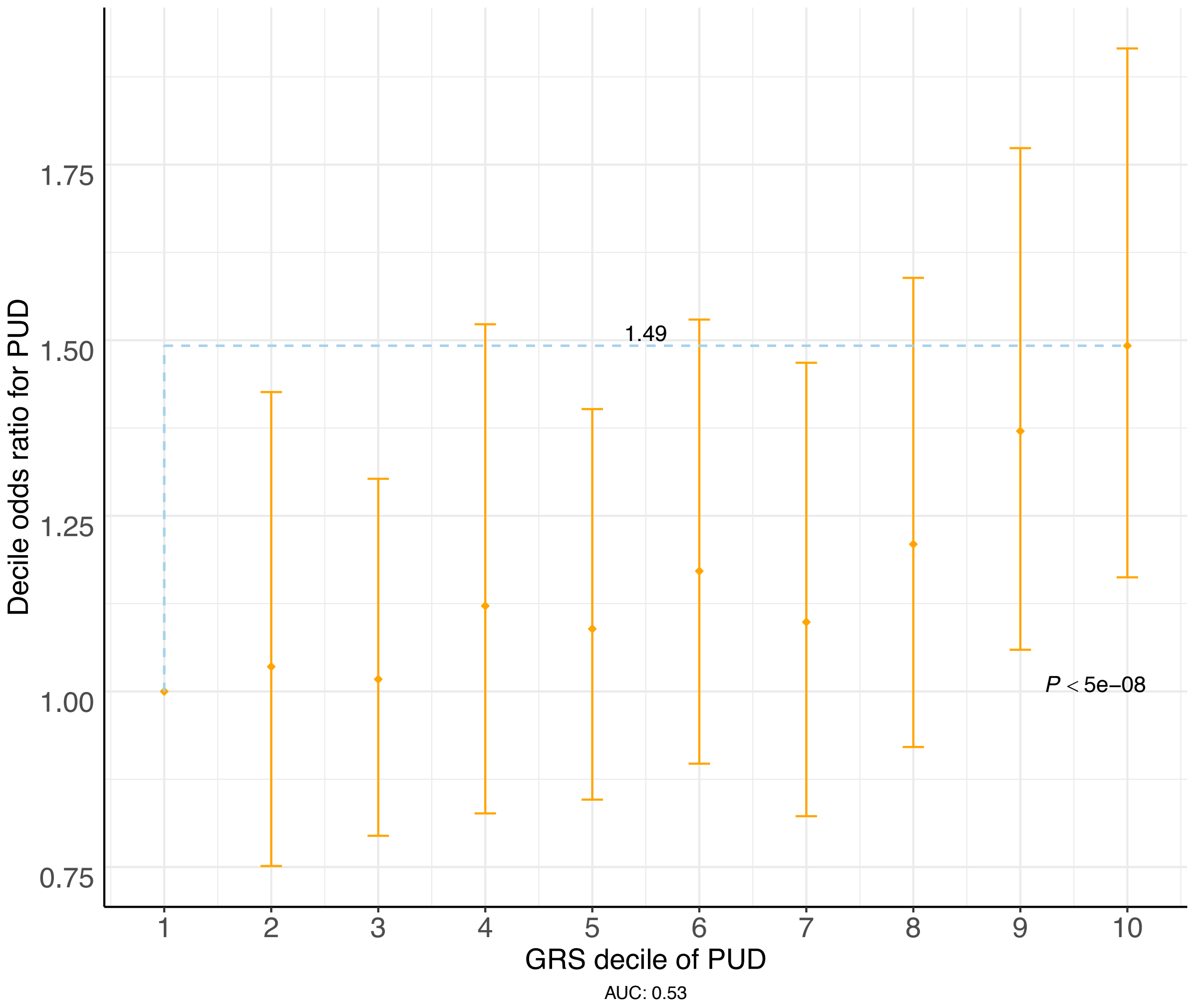

**Figure S10.** Genetic risk score (GRS) of peptic ulcer disease (PUD) predicts odds ratio (OR) for PUD in individuals from GERA cohort. GRS of individuals from GERA cohort were calculated based on PUD associated SNPs with P <5.0E-8 from UKB and converted to deciles (1 = lowest, 10 = highest). OR and 95% confidence intervals (CI, orange diamonds and bars) relative to decile 1 were estimated using logistic regression. The blue dashed lines represent that compared with the lowest decile, the highest decile have an OR of 1.49 for PUD. The number of PUD cases and controls from GERA cohort were 1,004 and 60,843, respectively. The P value for case-control PUD GRS difference from GERA cohort is 6.7E-5.
