## Supplementary data 1 for "Genome-wide association study of gastrointestinal disorders reinforces the link between the digestive tract and the nervous system"

Regional association plots of the 50 regions with genome-wide significant loci associated with the 5 digestion phenotypes in UK Biobank

### Gastro-oesophageal Reflux Disease (GORD)

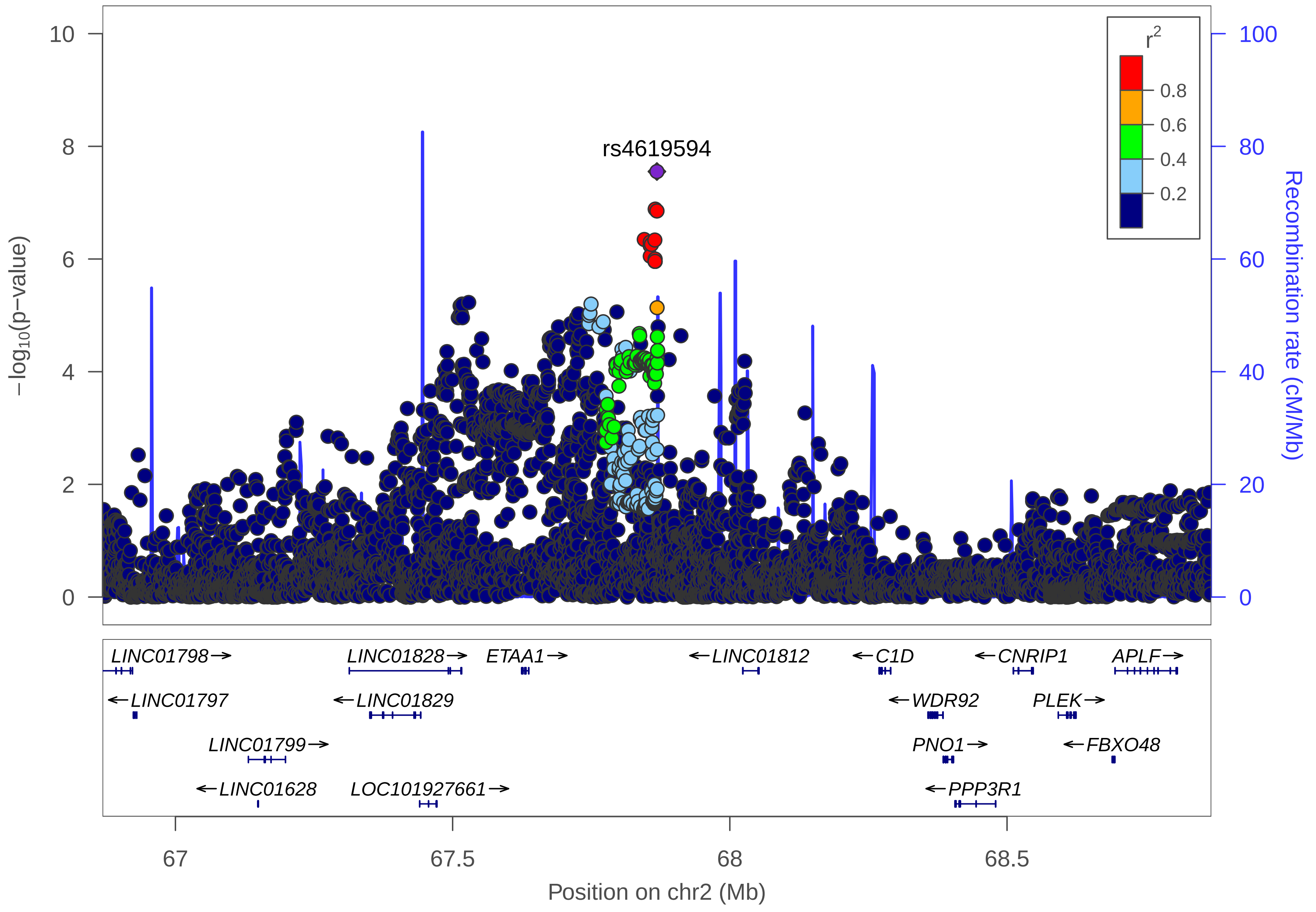
rs4619594
r2
0.8
0.6
0.4
0.2
−log10(p−value)
Recombination rate (cM/Mb)
0
2
4
6
8
10
0
20
40
60
80
100
LINC01798→
LINC01828→
ETAA1→
←LINC01812
←C1D
←CNRIP1
APLF→
←LINC01797
←LINC01829
←WDR92
PLEK→
LINC01799→
PNO1→
←FBXO48
←LINC01628
LOC101927661→
←PPP3R1
67
67.5
68
68.5
Position on chr2 (Mb)

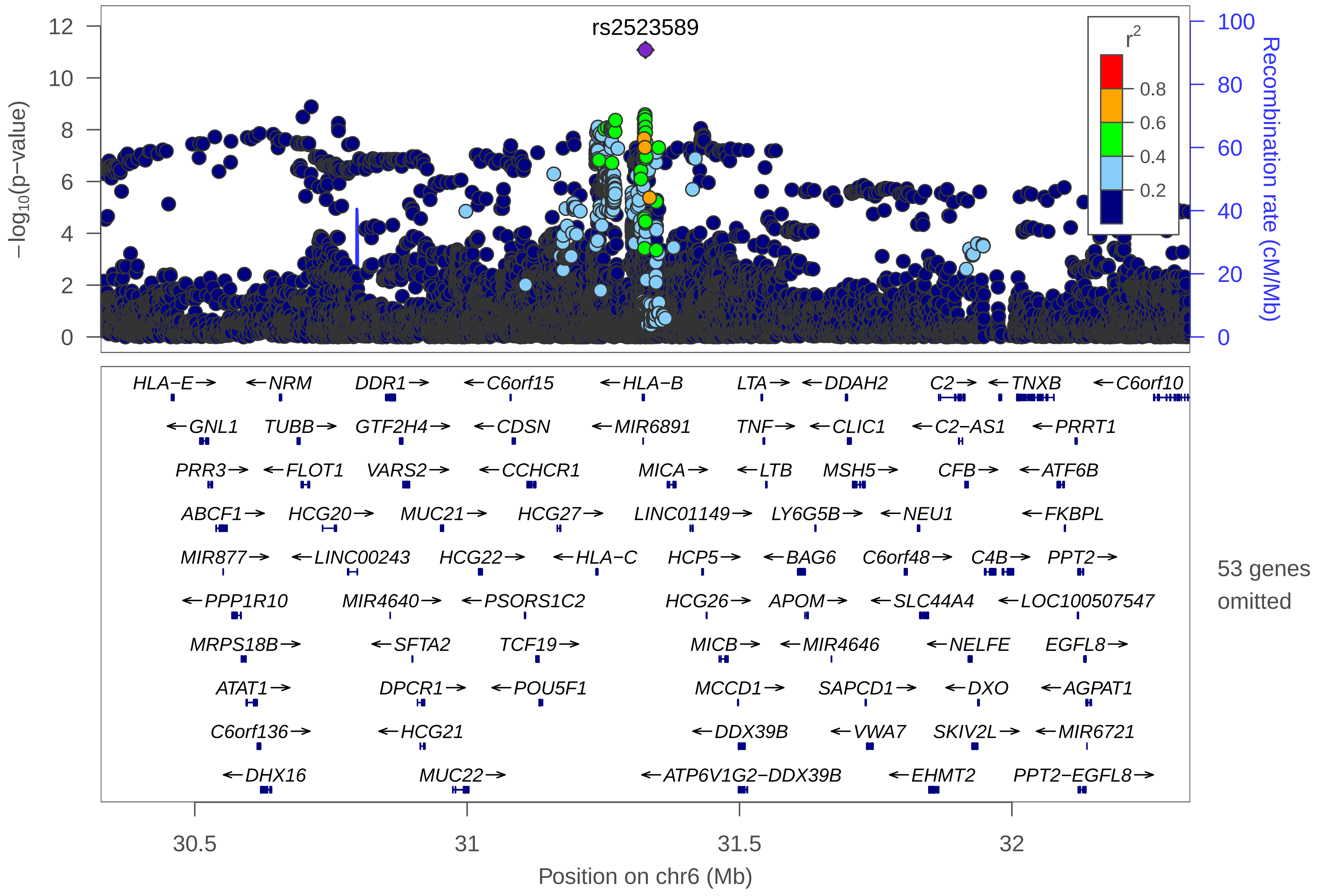
rs2523589
−log10(p−value)
Recombination rate (cM/Mb)
r2
0.8
0.6
0.4
0.2
0
2
4
6
8
10
12
20
40
60
80
100
HLA−E→
←NRM
DDR1→
←C6orf15
←HLA−B
LTA→
←DDAH2
C2→
←TNXB
←C6orf10
←GNL1
TUBB→
GTF2H4→
←CDSN
←MIR6891
TNF→
←CLIC1
←C2−AS1
←PRRT1
PRR3→
←FLOT1
VARS2→
←CCHCR1
MICA→
←LTB
MSH5→
CFB→
←ATF6B
ABCF1→
HCG20→
MUC21→
HCG27→
LINC01149→
LY6G5B→
←NEU1
←FKBPL
MIR877→
←LINC00243
HCG22→
←HLA−C
HCP5→
←BAG6
C6orf48→
C4B→
PPT2→
←PPP1R10
MIR4640→
←PSORS1C2
HCG26→
APOM→
←SLC44A4
←LOC100507547
MRPS18B→
←SFTA2
TCF19→
MICB→
←MIR4646
←NELFE
EGFL8→
ATAT1→
DPCR1→
←POU5F1
MCCD1→
SAPCD1→
←DXO
←AGPAT1
C6orf136→
←HCG21
←DDX39B
←VWA7
SKIV2L→
←MIR6721
←DHX16
MUC22→
←ATP6V1G2−DDX39B
←EHMT2
PPT2−EGFL8→
53 genes omitted
30.5
31
31.5
32
Position on chr6 (Mb)

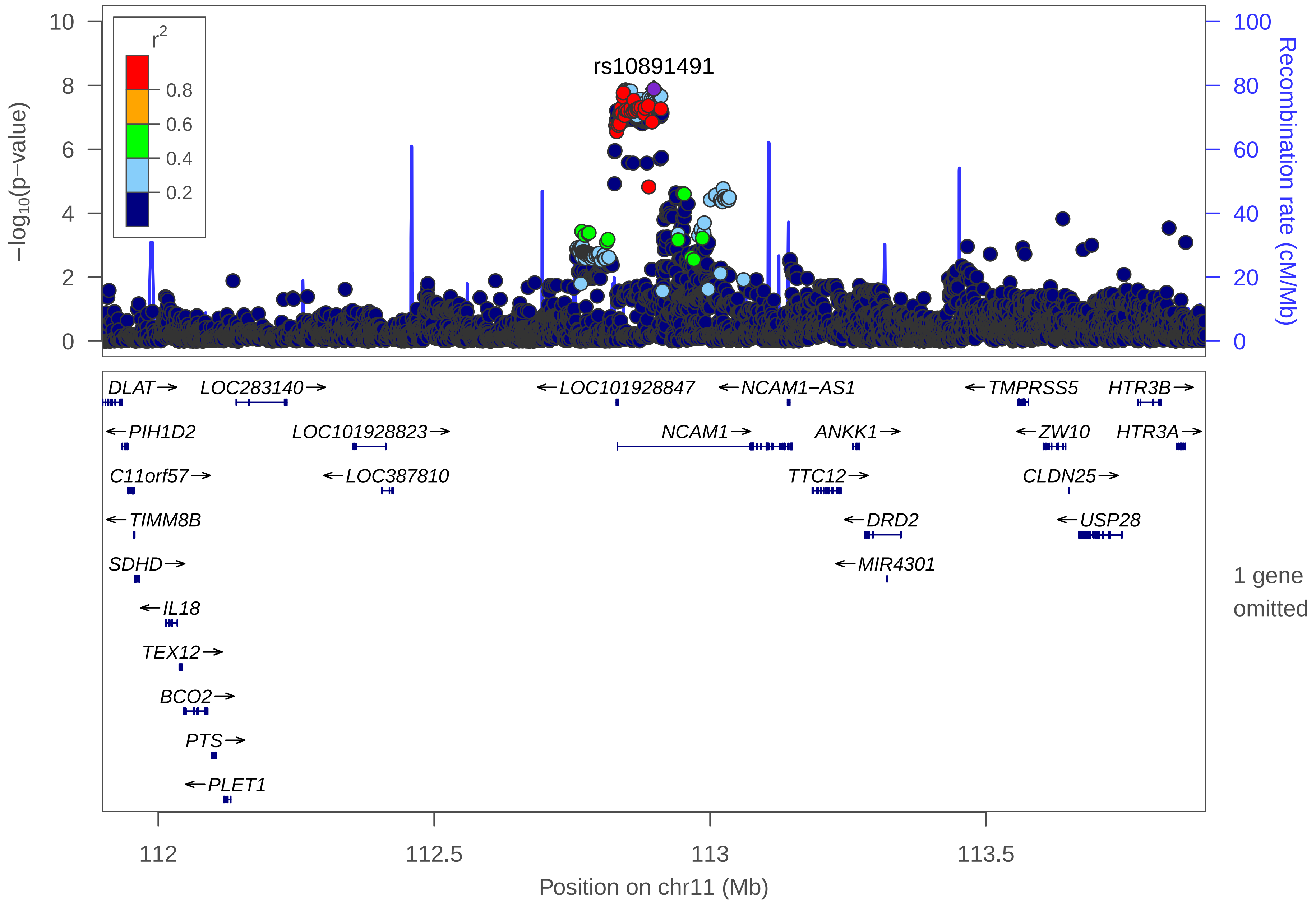

rs10891491
r2
0.8
0.6
0.4
0.2
−log10(p−value)
Recombination rate (cM/Mb)
0
2
4
6
8
10
0
20
40
60
80
100
DLAT
LOC283140
LOC101928847
NCAM1−AS1
TMPRSS5
HTR3B
PIH1D2
LOC101928823
NCAM1
ANKK1
ZW10
HTR3A
C11orf57
LOC387810
TTC12
CLDN25
TIMM8B
DRD2
USP28
SDHD
MIR4301
IL18
TEX12
BCO2
PTS
PLET1
1 gene omitted
112
112.5
113
113.5
Position on chr11 (Mb)

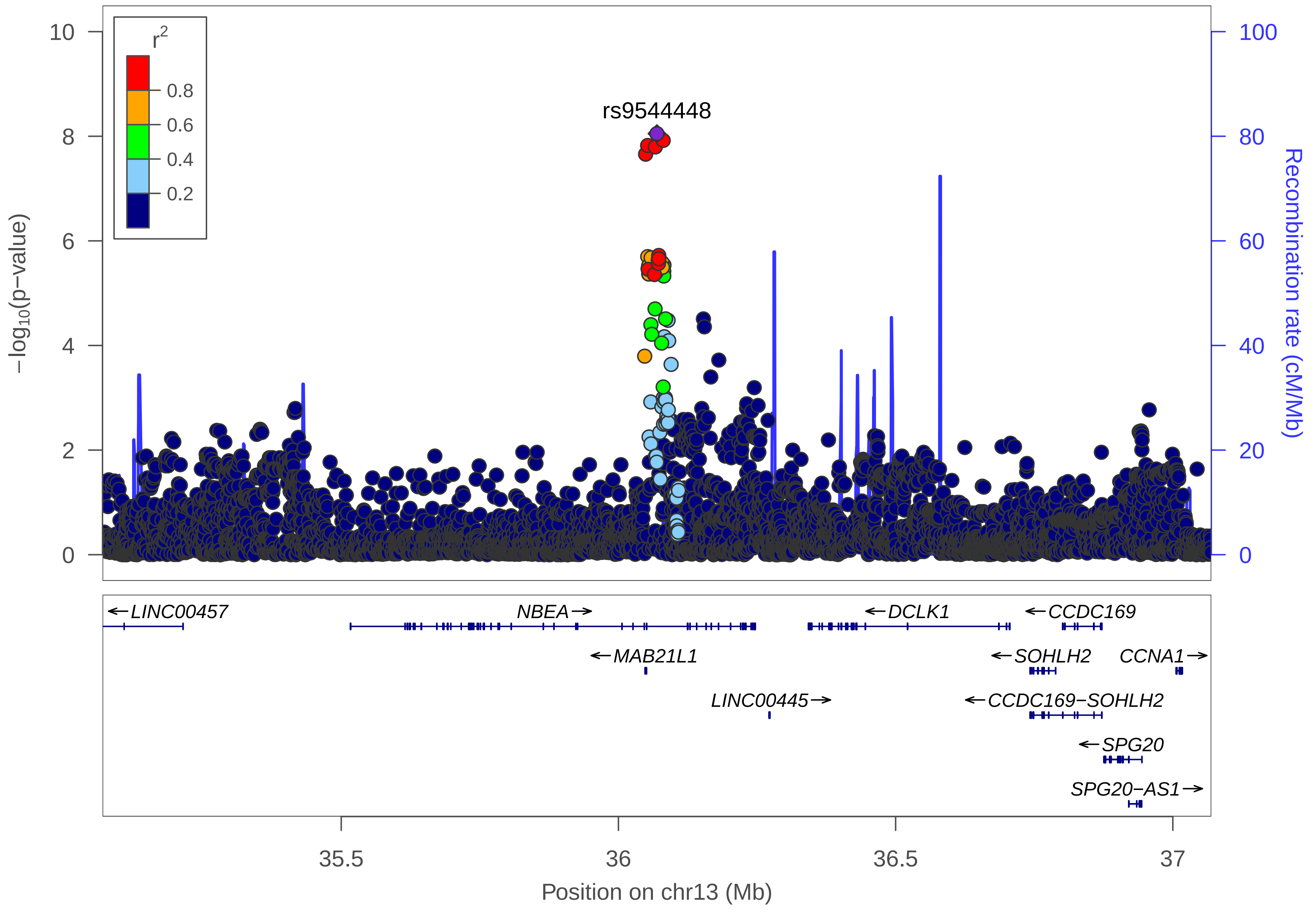

r2
0.8
0.6
0.4
0.2
rs9544448
10
8
6
4
2
0
−log10(p−value)
100
80
60
40
20
0
Recombination rate (cM/Mb)
←LINC00457
NBEA→
←DCLK1
←CCDC169
←MAB21L1
←SOHLH2
CCNA1→
LINC00445→
←CCDC169−SOHLH2
←SPG20
SPG20−AS1→
35.5
36
36.5
37
Position on chr13 (Mb)

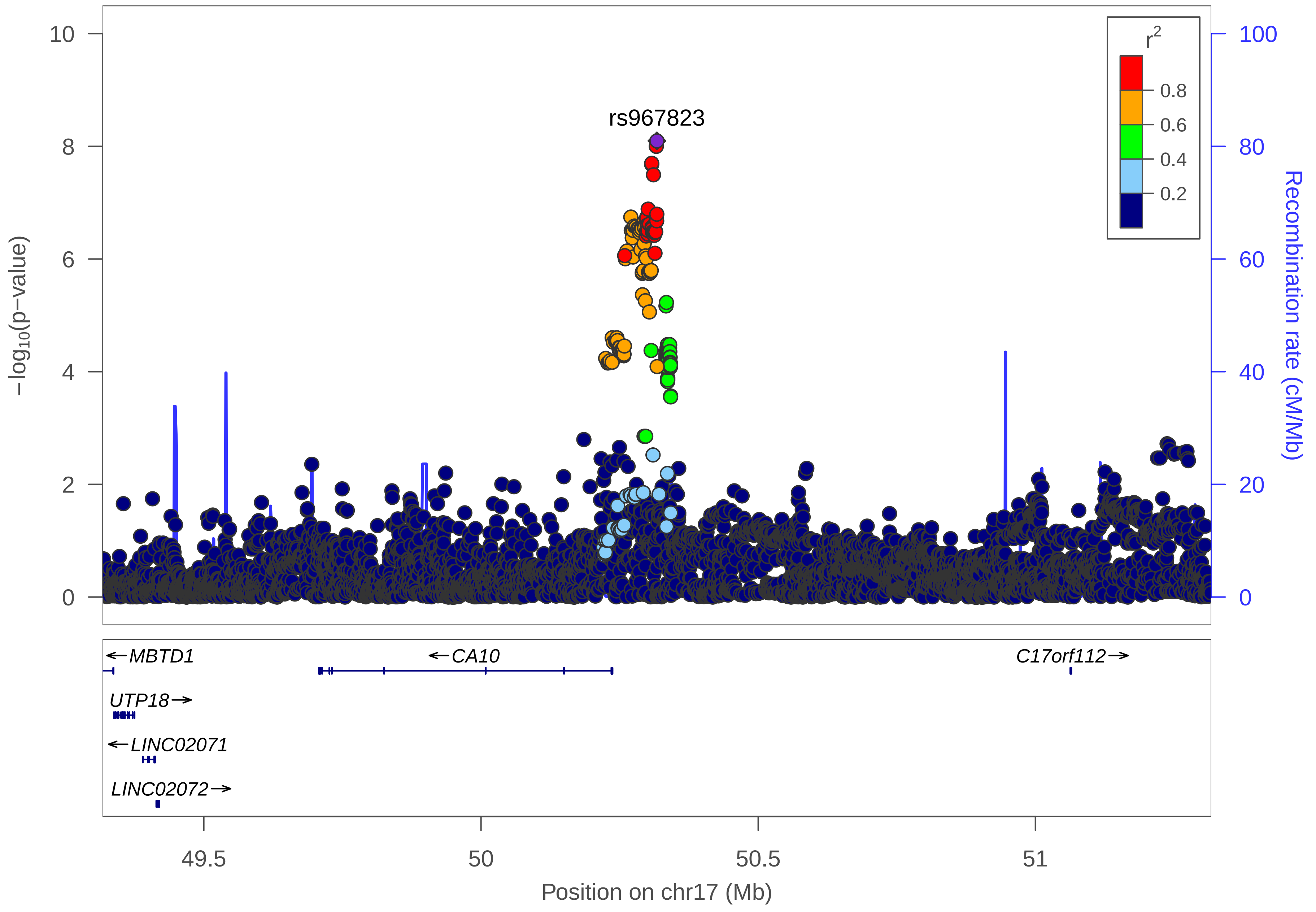
rs967823
r2
0.8
0.6
0.4
0.2
−log10(p−value)
Recombination rate (cM/Mb)
0
2
4
6
8
10
0
20
40
60
80
100
MBTD1
CA10
C17orf112
UTP18
LINC02071
LINC02072
49.5
50
50.5
51
Position on chr17 (Mb)

### Peptic Ulcer Disease (PUD)

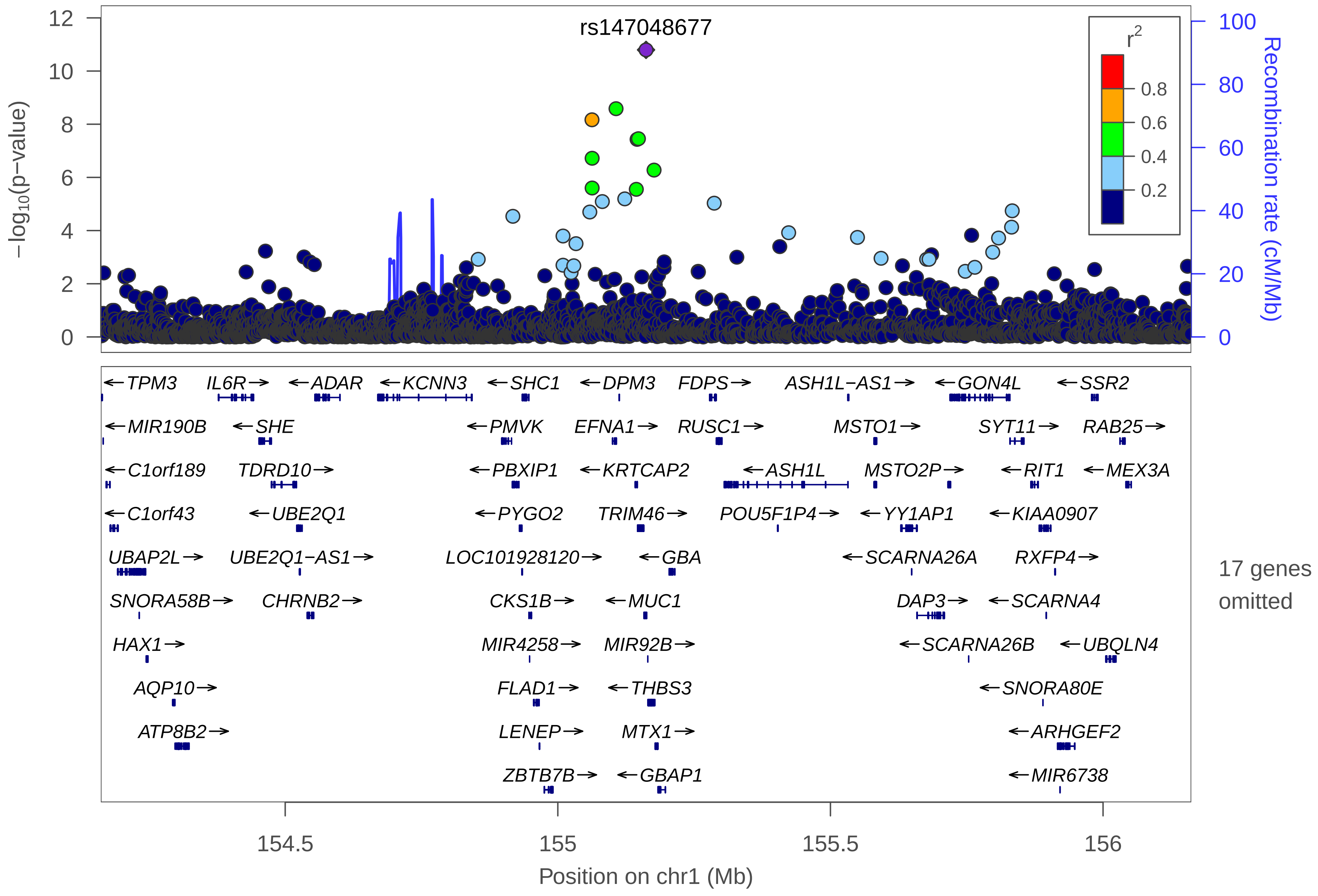

rs147048677
−log10(p−value)
Recombination rate (cM/Mb)
r2
0.8
0.6
0.4
0.2
0
2
4
6
8
10
12
0
20
40
60
80
100
←TPM3
IL6R→
←ADAR
←KCNN3
←SHC1
←DPM3
FDPS→
ASH1L−AS1→
←GON4L
←SSR2
←MIR190B
←SHE
←PMVK
EFNA1→
RUSC1→
MSTO1→
SYT11→
RAB25→
←C1orf189
TDRD10→
←PBXIP1
←KRTCAP2
←ASH1L
MSTO2P→
←RIT1
←MEX3A
←C1orf43
←UBE2Q1
←PYGO2
TRIM46→
POU5F1P4→
←YY1AP1
←KIAA0907
UBAP2L→
UBE2Q1−AS1→
LOC101928120→
←GBA
←SCARNA26A
RXFP4→
SNORA58B→
CHRNB2→
CKS1B→
←MUC1
DAP3→
←SCARNA4
HAX1→
MIR4258→
MIR92B→
←SCARNA26B
←UBQLN4
AQP10→
FLAD1→
←THBS3
←SNORA80E
ATP8B2→
LENEP→
MTX1→
←ARHGEF2
ZBTB7B→
←GBAP1
←MIR6738
17 genes omitted
154.5
155
155.5
156
Position on chr1 (Mb)

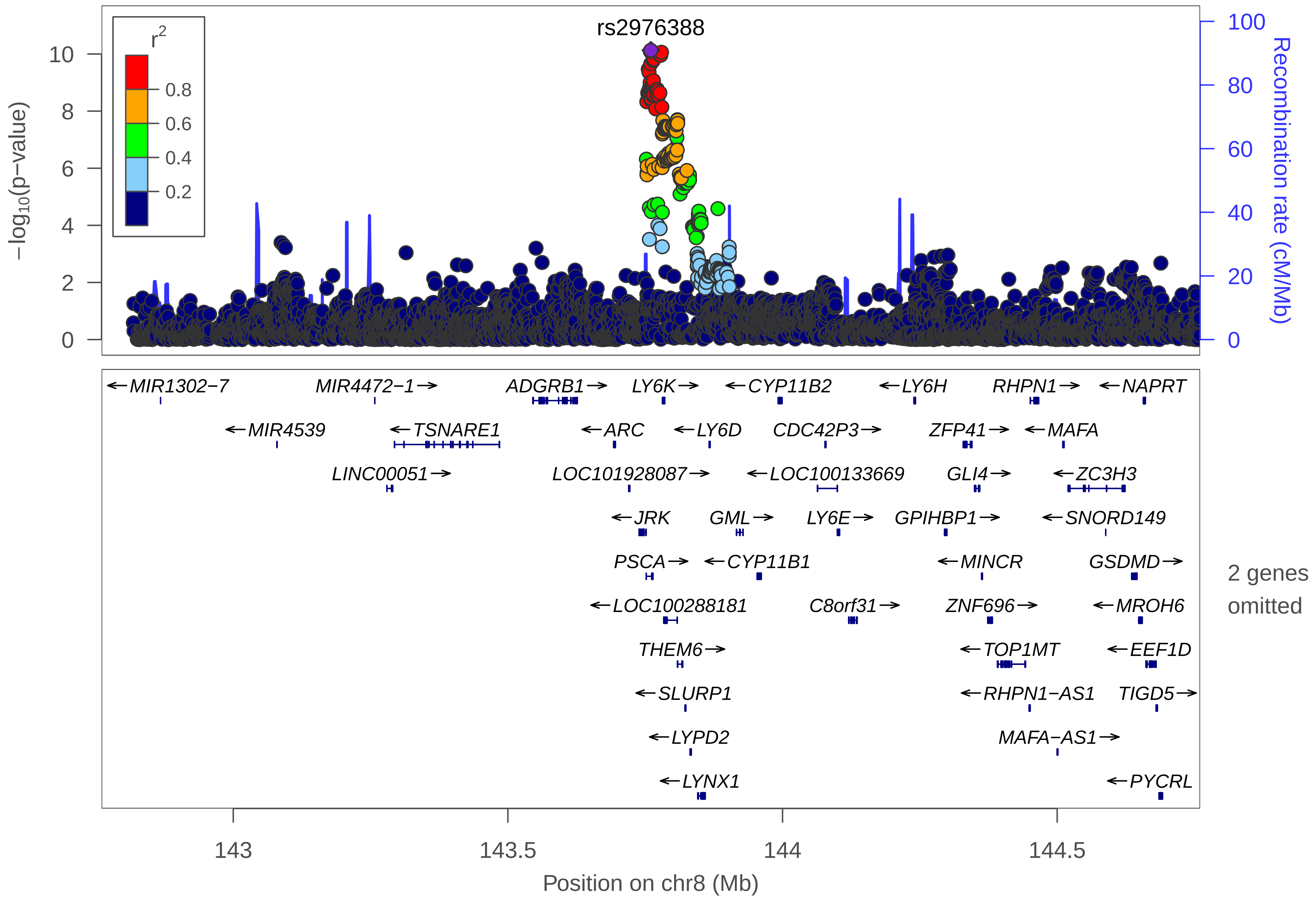
rs2976388
r2
0.8
0.6
0.4
0.2
−log10(p−value)
Recombination rate (cM/Mb)
MIR1302−7
MIR4472−1
ADGRB1
LY6K
CYP11B2
LY6H
RHPN1
NAPRT
MIR4539
TSNARE1
ARC
LY6D
CDC42P3
ZFP41
MAFA
LINC00051
LOC101928087
LOC100133669
GLI4
ZC3H3
JRK
GML
LY6E
GPIHBP1
SNORD149
PSCA
CYP11B1
MINCR
GSDMD
LOC100288181
C8orf31
ZNF696
MROH6
THEM6
TOP1MT
EEF1D
SLURP1
RHPN1−AS1
TIGD5
LYPD2
MAFA−AS1
LYNX1
PYCRL
2 genes omitted
143
143.5
144
144.5
Position on chr8 (Mb)

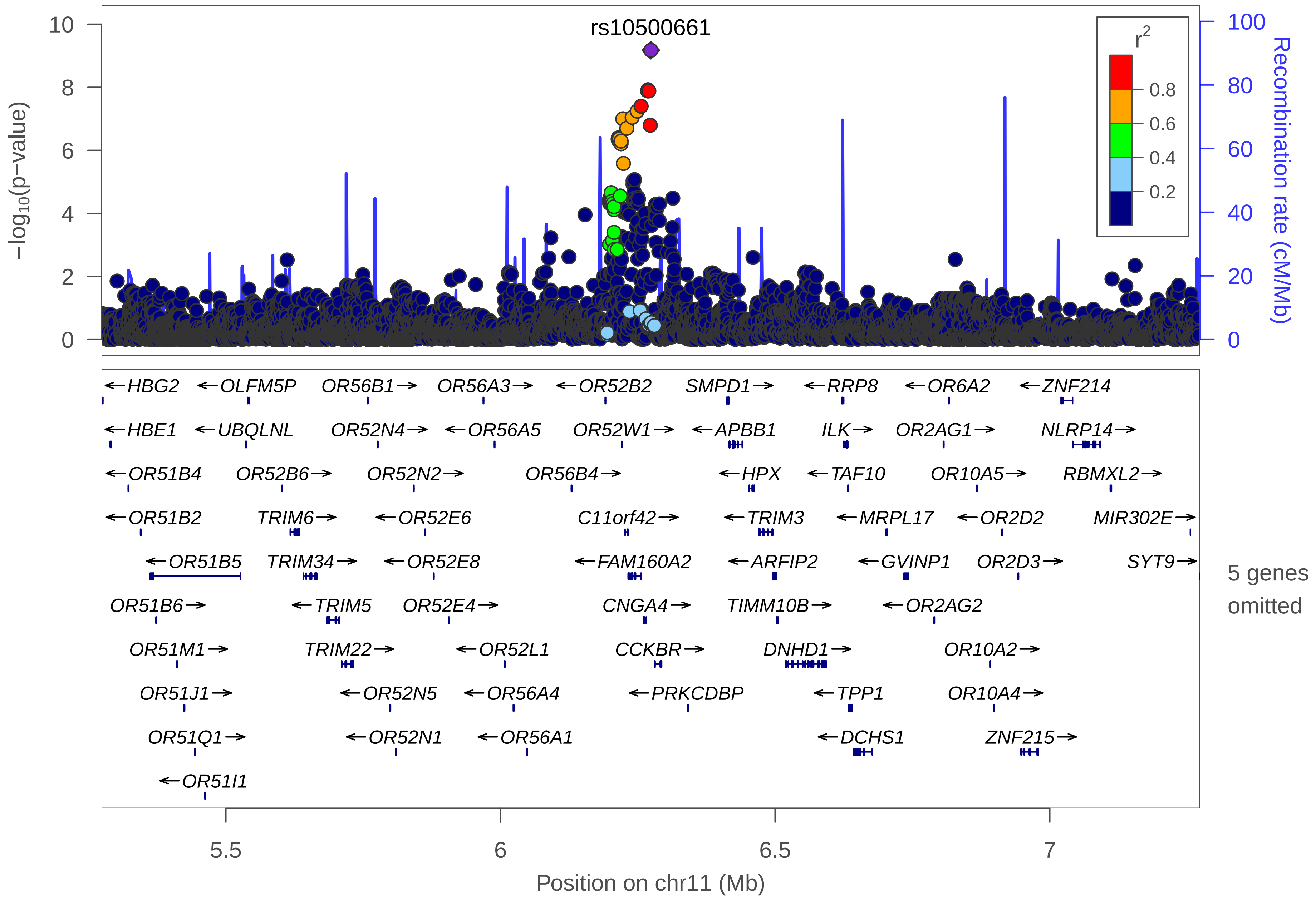

rs10500661
r2
0.8
0.6
0.4
0.2
-log10(p-value)
Recombination rate (cM/Mb)
10
8
6
4
2
0
100
80
60
40
20
0
←HBG2
←OLFM5P
OR56B1→
OR56A3→
←OR52B2
SMPD1→
←RRP8
←OR6A2
←ZNF214
←HBE1
←UBQLNL
OR52N4→
←OR56A5
OR52W1→
←APBB1
ILK→
OR2AG1→
NLRP14→
←OR51B4
OR52B6→
OR52N2→
OR56B4→
←HPX
←TAF10
OR10A5→
RBMXL2→
←OR51B2
TRIM6→
←OR52E6
C11orf42→
←TRIM3
←MRPL17
←OR2D2
MIR302E→
←OR51B5
TRIM34→
←OR52E8
←FAM160A2
←ARFIP2
←GVINP1
OR2D3→
SYT9→
OR51B6→
←TRIM5
OR52E4→
CNGA4→
TIMM10B→
←OR2AG2
OR51M1→
TRIM22→
←OR52L1
CCKBR→
DNHD1→
OR10A2→
OR51J1→
←OR52N5
←OR56A4
←PRKCDBP
←TPP1
OR10A4→
OR51Q1→
←OR52N1
←OR56A1
←DCHS1
ZNF215→
←OR51I1
5 genes omitted
5.5
6
6.5
7
Position on chr11 (Mb)

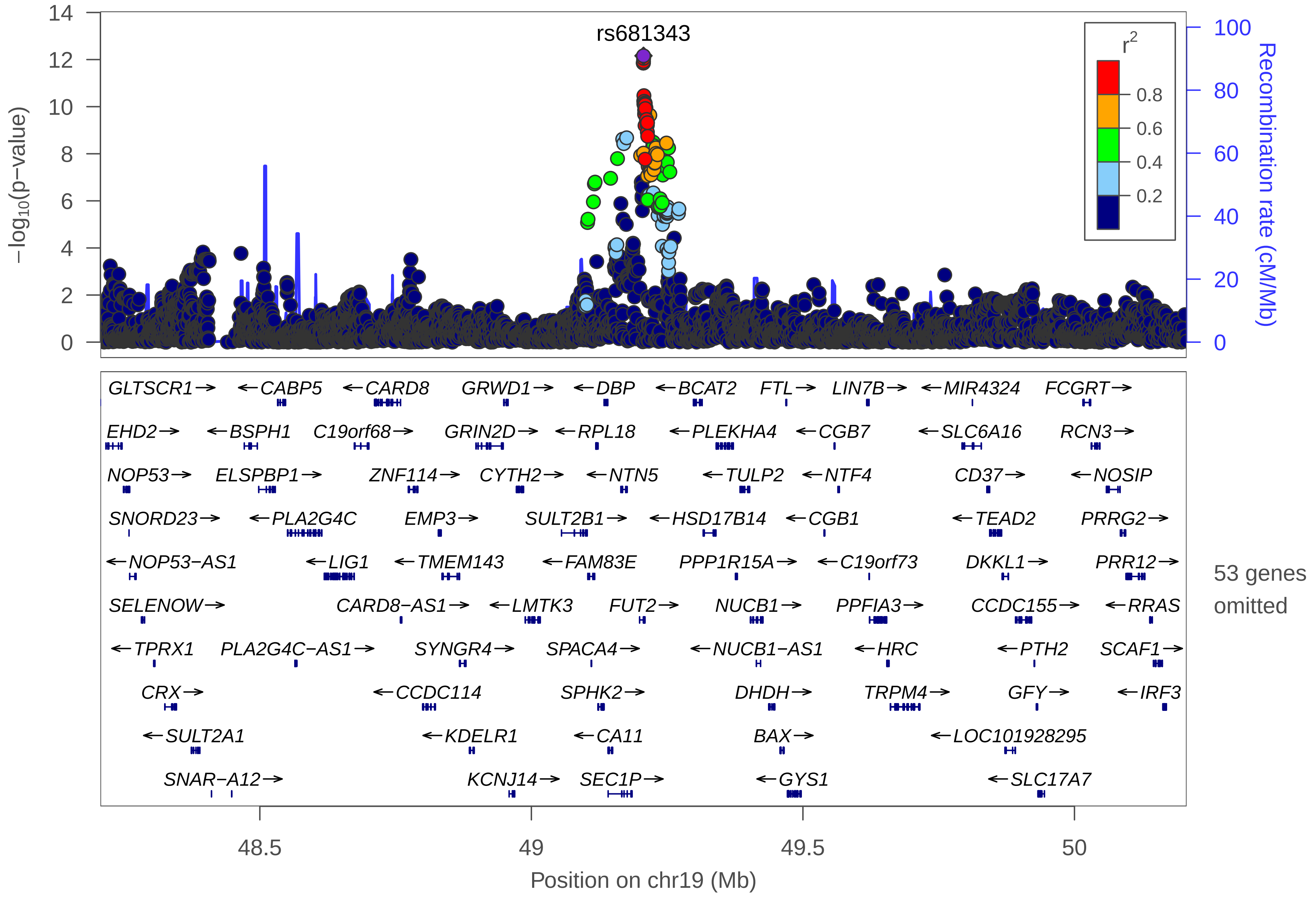
rs681343
r2
0.8
0.6
0.4
0.2
−log10(p−value)
Recombination rate (cM/Mb)
0
2
4
6
8
10
12
14
0
20
40
60
80
100
GLTSCR1→
←CABP5
←CARD8
GRWD1→
←DBP
←BCAT2
FTL→
LIN7B→
←MIR4324
FCGRT→
EHD2→
←BSPH1
C19orf68→
GRIN2D→
←RPL18
←PLEKHA4
←CGB7
←SLC6A16
RCN3→
NOP53→
ELSPBP1→
ZNF114→
CYTH2→
←NTN5
←TULP2
←NTF4
CD37→
←NOSIP
SNORD23→
←PLA2G4C
EMP3→
SULT2B1→
←HSD17B14
←CGB1
←TEAD2
PRRG2→
←NOP53−AS1
←LIG1
←TMEM143
←FAM83E
PPP1R15A→
←C19orf73
DKKL1→
PRR12→
SELENOW→
CARD8−AS1→
←LMTK3
FUT2→
NUCB1→
PPFIA3→
CCDC155→
←RRAS
←TPRX1
PLA2G4C−AS1→
SYNGR4→
SPACA4→
←NUCB1−AS1
←HRC
←PTH2
SCAF1→
CRX→
←CCDC114
SPHK2→
DHDH→
TRPM4→
GFY→
←IRF3
←SULT2A1
←KDELR1
←CA11
BAX→
←LOC101928295
SNAR−A12→
KCNJ14→
SEC1P→
←GYS1
←SLC17A7
53 genes omitted
48.5
49
49.5
50
Position on chr19 (Mb)

### Gastro-oesophageal Reflux Disease (GORD), Peptic Ulcer Disease (PUD) and corresponding medications (GP+M)

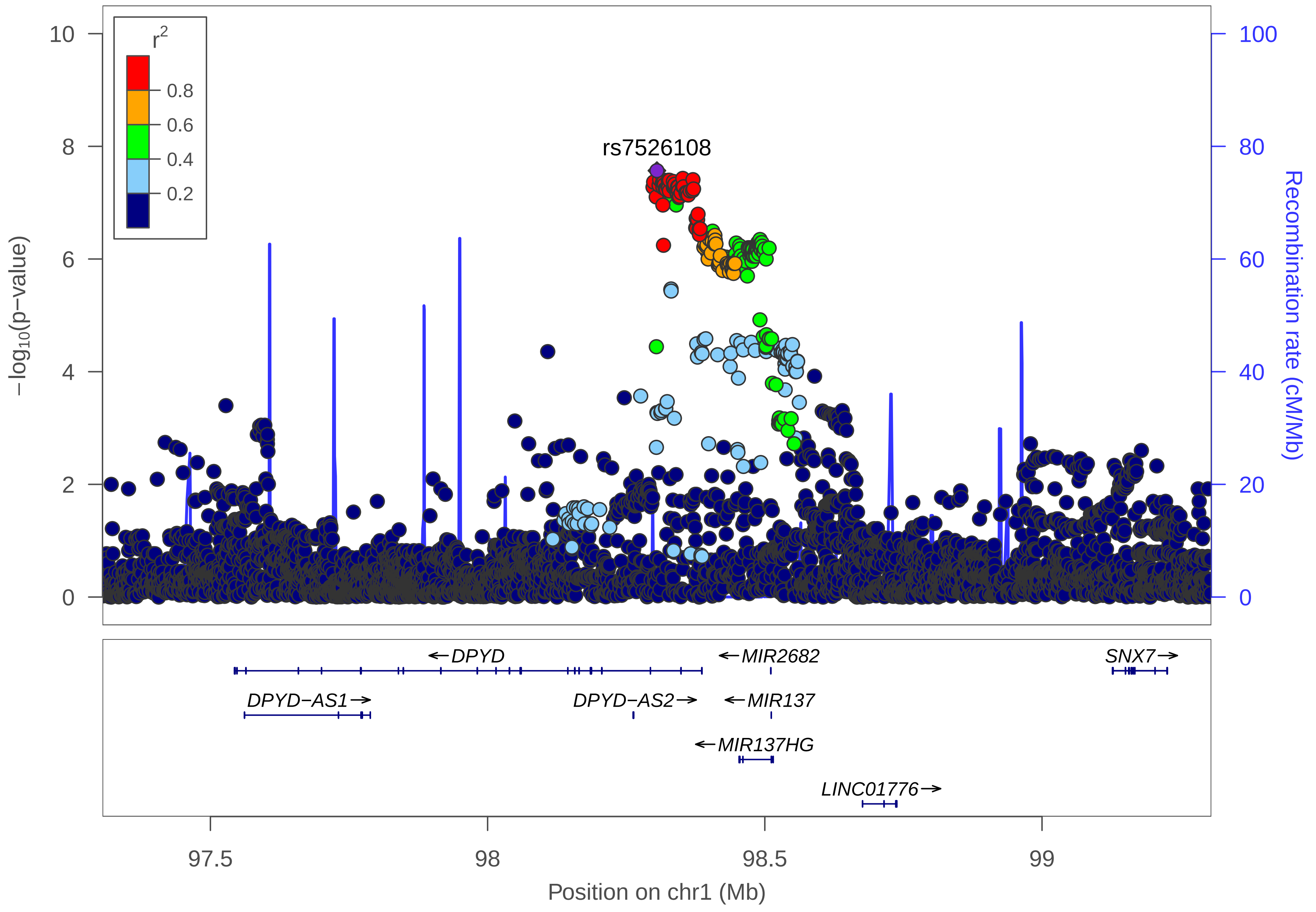

rs7526108
r2
0.8
0.6
0.4
0.2
−log10(p−value)
Recombination rate (cM/Mb)
10
8
6
4
2
0
100
80
60
40
20
0
←DPYD
←MIR2682
SNX7→
DPYD−AS1→
DPYD−AS2→
←MIR137
←MIR137HG
LINC01776→
97.5
98
98.5
99
Position on chr1 (Mb)

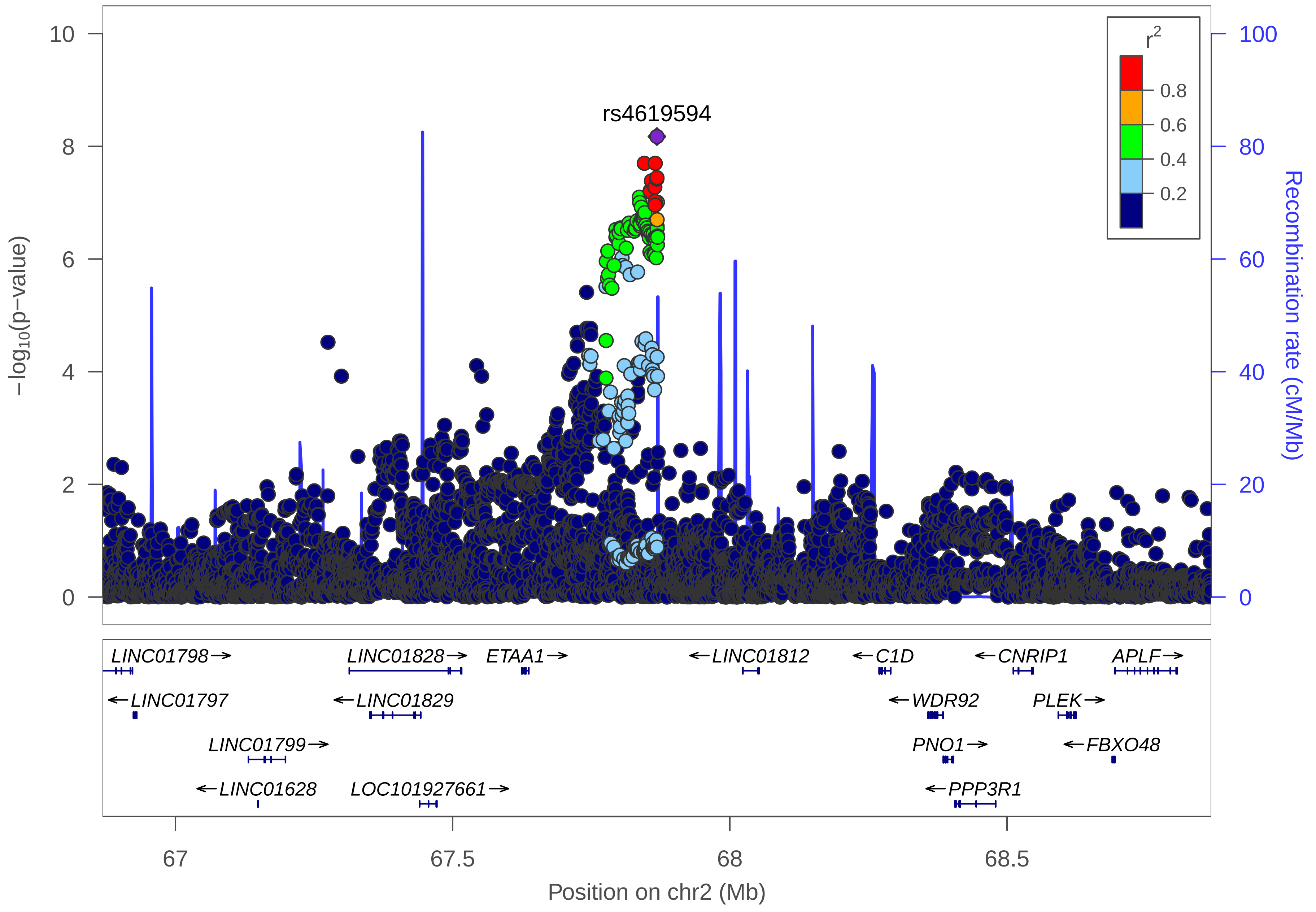

rs4619594
r2
0.8
0.6
0.4
0.2
−log10(p−value)
Recombination rate (cM/Mb)
10
8
6
4
2
0
100
80
60
40
20
0
LINC01798→
←LINC01797
LINC01799→
←LINC01628
LINC01828→
←LINC01829
LOC101927661→
ETAA1→
←LINC01812
←C1D
←WDR92
PNO1→
←PPP3R1
←CNRIP1
PLEK→
←FBXO48
APLF→
67
67.5
68
68.5
Position on chr2 (Mb)

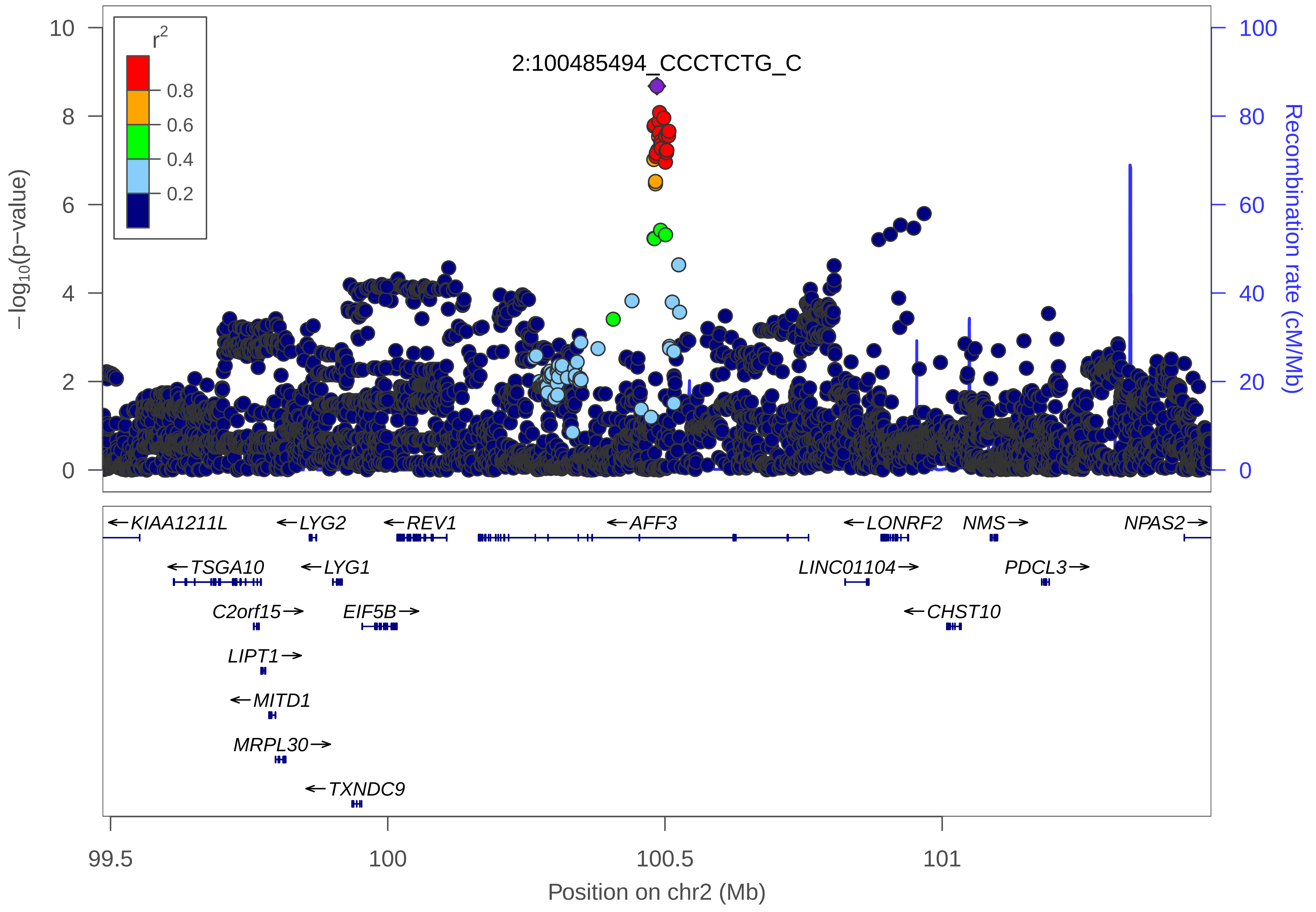

r2
0.8
0.6
0.4
0.2
2:100485494_CCCTCTG_C
−log10(p−value)
Recombination rate (cM/Mb)
10
8
6
4
2
0
100
80
60
40
20
0
←KIAA1211L
←LYG2
←REV1
←AFF3
←LONRF2
NMS→
NPAS2→
←TSGA10
←LYG1
LINC01104→
PDCL3→
C2orf15→
EIF5B→
←CHST10
LIPT1→
←MITD1
MRPL30→
←TXNDC9
99.5
100
100.5
101
Position on chr2 (Mb)

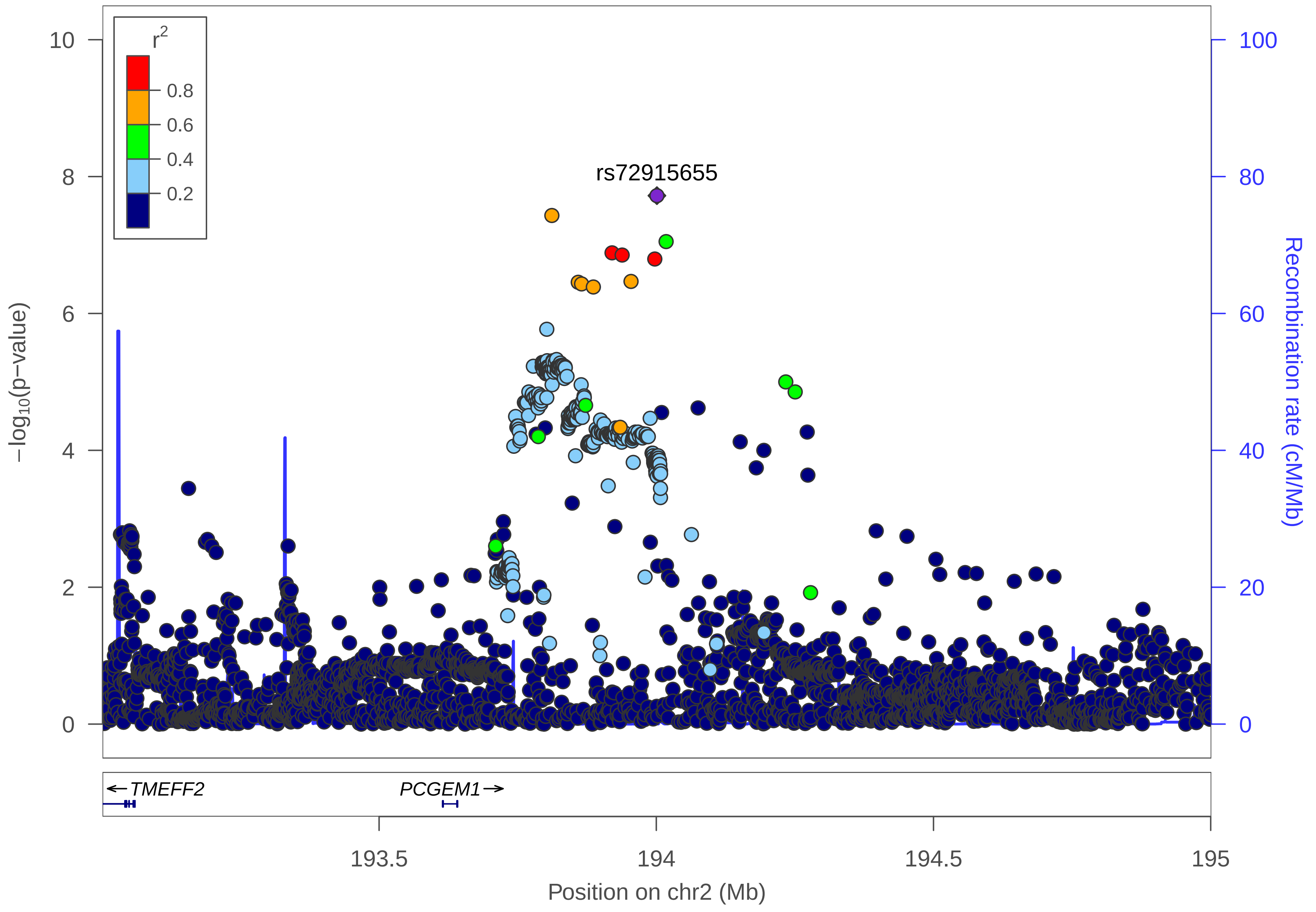

r2
0.8
0.6
0.4
0.2
rs72915655
10
8
6
4
2
0
−log10(p−value)
100
80
60
40
20
0
Recombination rate (cM/Mb)
← TMEFF2
PCGEM1 →
193.5
194
194.5
195
Position on chr2 (Mb)

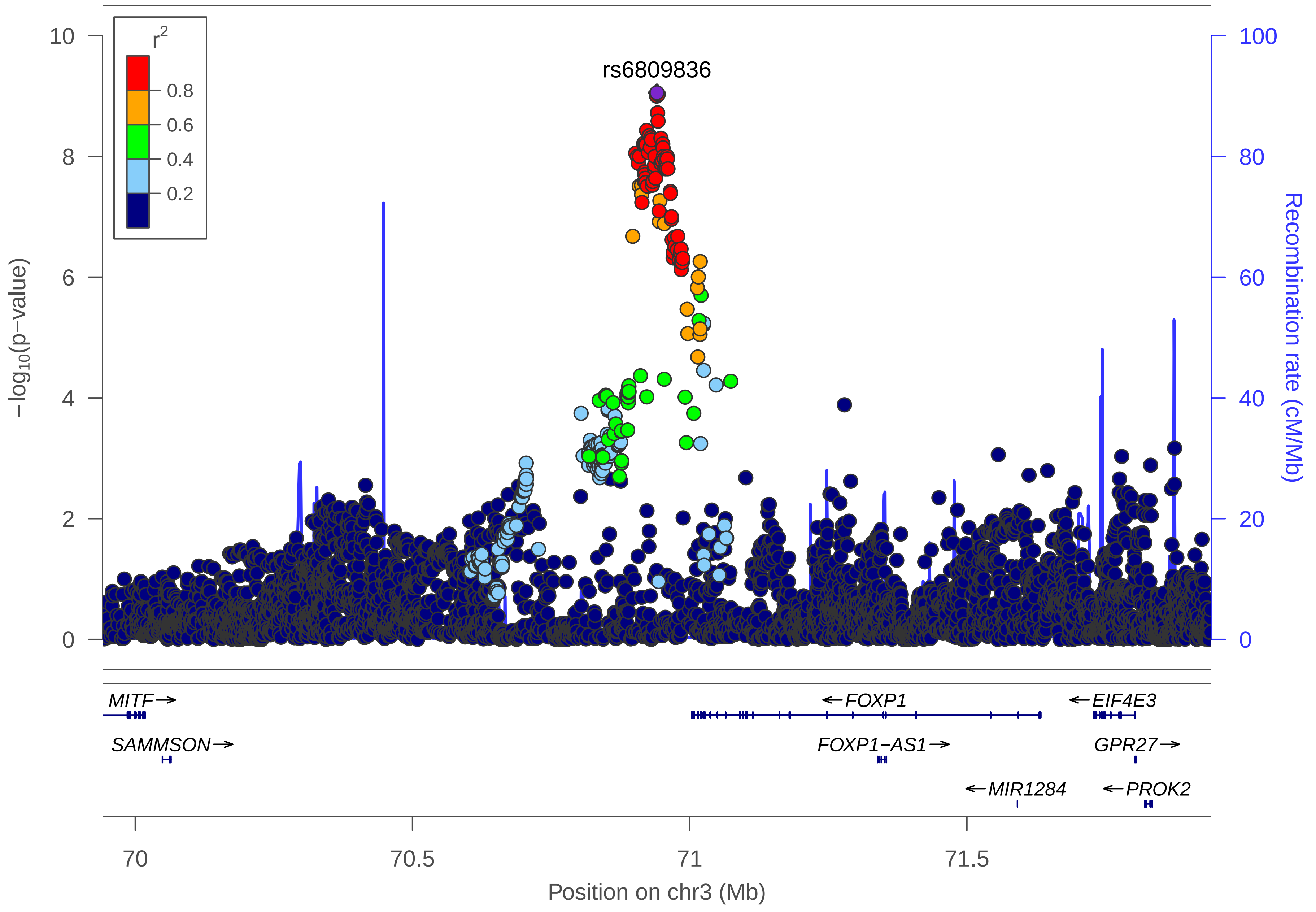

r2
0.8
0.6
0.4
0.2
rs6809836
−log10(p−value)
Recombination rate (cM/Mb)
10
8
6
4
2
0
100
80
60
40
20
0
MITF→
SAMMSON→
←FOXP1
FOXP1−AS1→
←MIR1284
←EIF4E3
GPR27→
←PROK2
70
70.5
71
71.5
Position on chr3 (Mb)

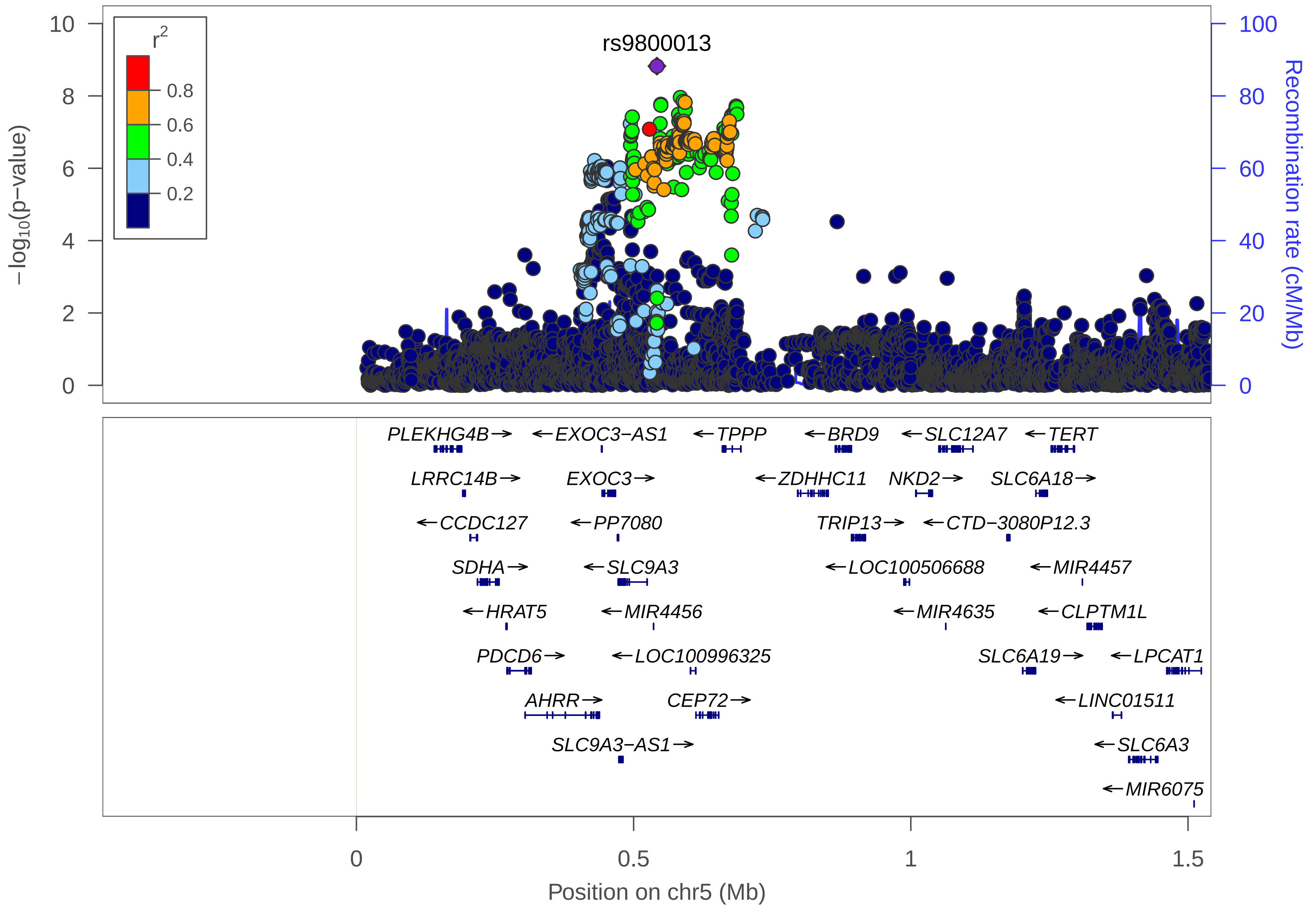
rs9800013
r2
0.8
0.6
0.4
0.2
−log10(p−value)
Recombination rate (cM/Mb)
10
8
6
4
2
0
100
80
60
40
20
0
PLEKHG4B→
←EXOC3−AS1
←TPPP
←BRD9
←SLC12A7
←TERT
LRRC14B→
EXOC3→
←ZDHHC11
NKD2→
SLC6A18→
←CCDC127
←PP7080
TRIP13→
←CTD−3080P12.3
SDHA→
←SLC9A3
←LOC100506688
←MIR4457
←HRAT5
←MIR4456
←MIR4635
←CLPTM1L
PDCD6→
←LOC100996325
SLC6A19→
←LPCAT1
AHRR→
CEP72→
←LINC01511
SLC9A3−AS1→
←SLC6A3
←MIR6075
0
0.5
1
1.5
Position on chr5 (Mb)

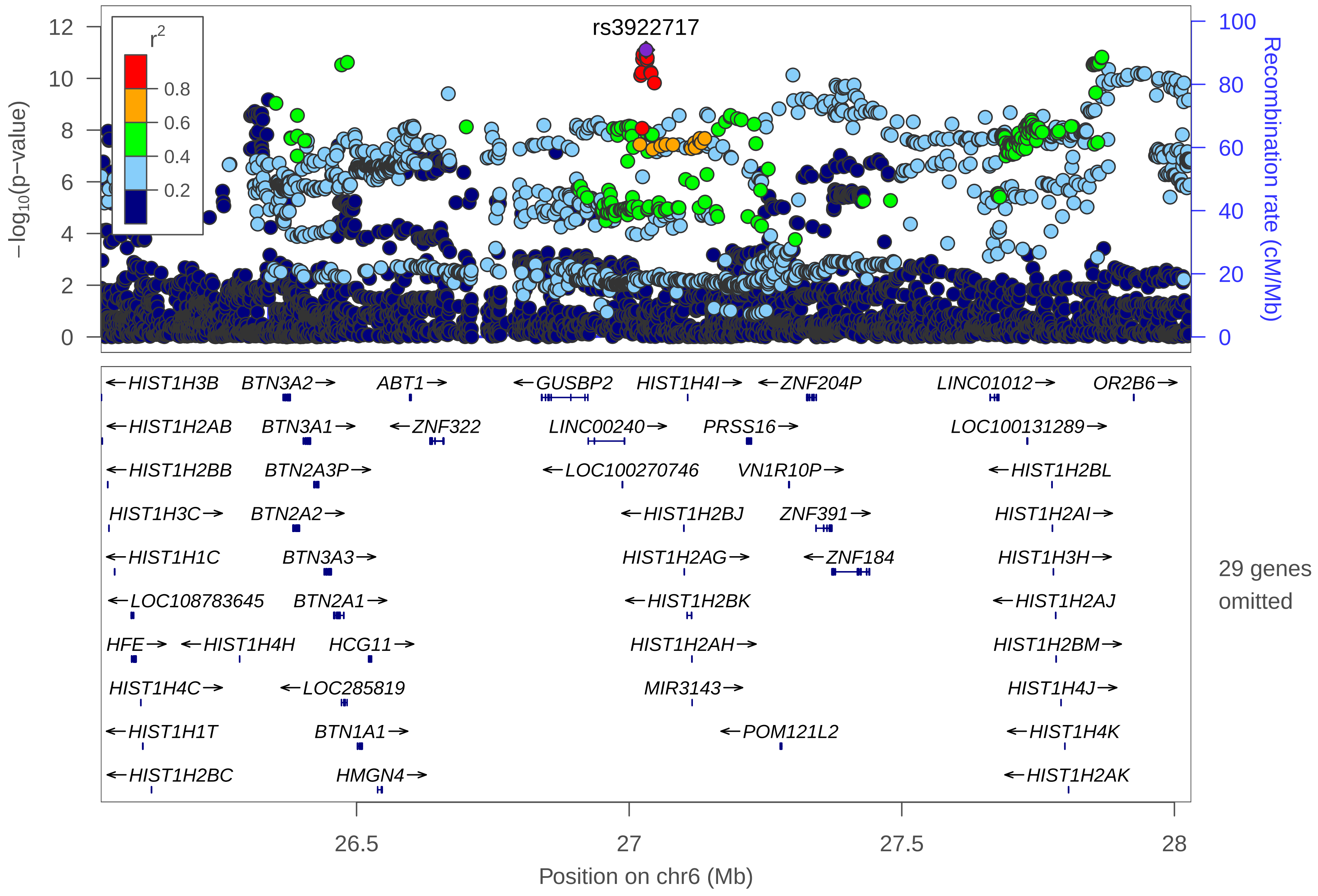

rs3922717
r2
0.8
0.6
0.4
0.2
−log10(p−value)
Recombination rate (cM/Mb)
HIST1H3B
BTN3A2
ABT1
GUSBP2
HIST1H4I
ZNF204P
LINC01012
OR2B6
HIST1H2AB
BTN3A1
ZNF322
LINC00240
PRSS16
LOC100131289
HIST1H2BB
BTN2A3P
LOC100270746
VN1R10P
HIST1H2BL
HIST1H3C
BTN2A2
HIST1H2BJ
ZNF391
HIST1H2AI
HIST1H1C
BTN3A3
HIST1H2AG
ZNF184
HIST1H3H
LOC108783645
BTN2A1
HIST1H2BK
HIST1H2AJ
HFE
HIST1H4H
HCG11
HIST1H2AH
HIST1H2BM
HIST1H4C
LOC285819
MIR3143
HIST1H4J
HIST1H1T
BTN1A1
POM121L2
HIST1H4K
HIST1H2BC
HMGN4
HIST1H2AK
29 genes omitted
26.5
27
27.5
28
Position on chr6 (Mb)

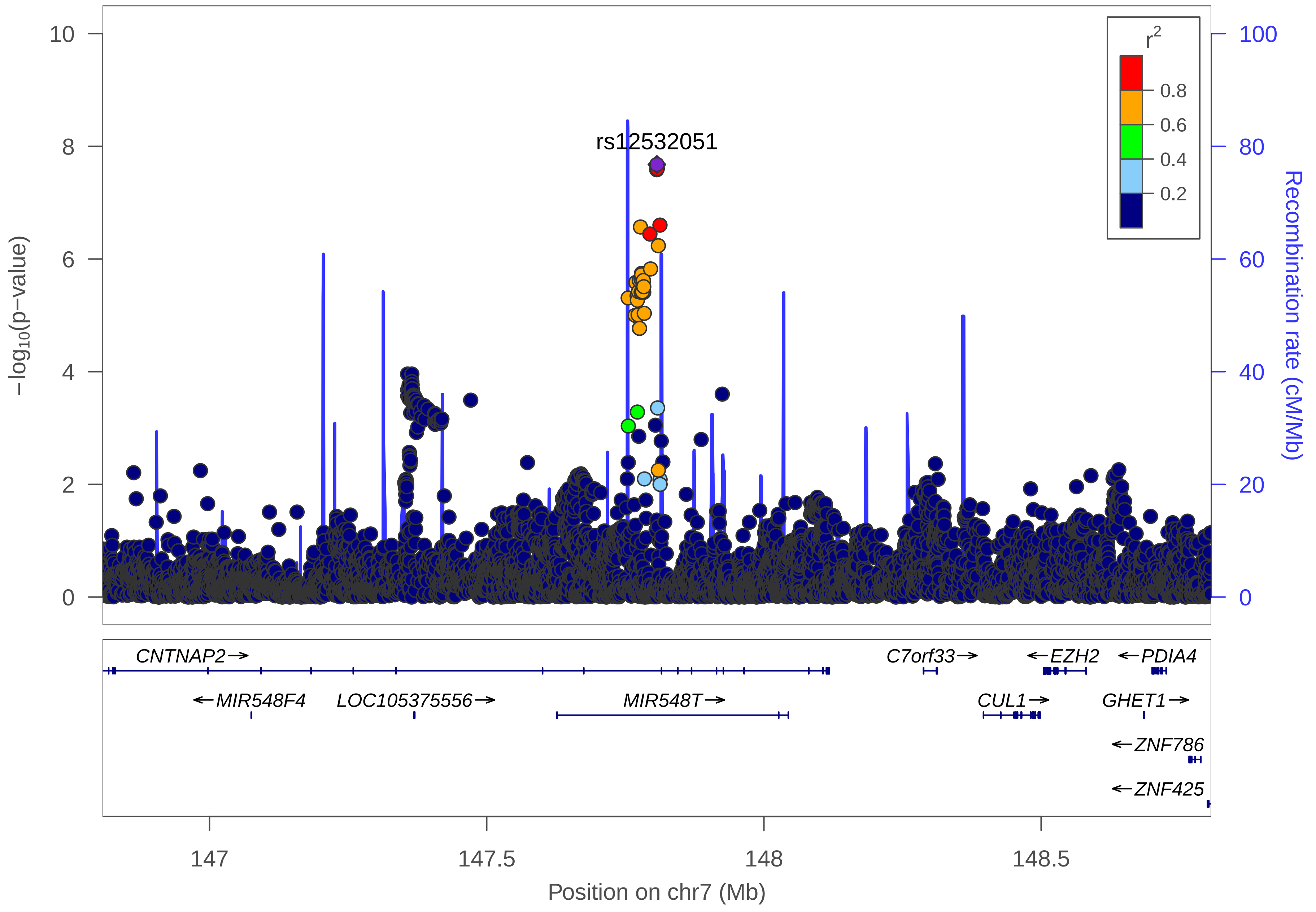

rs12532051
r2
0.8
0.6
0.4
0.2
−log10(p−value)
Recombination rate (cM/Mb)
0
2
4
6
8
10
0
20
40
60
80
100
CNTNAP2→
←MIR548F4
LOC105375556→
MIR548T→
C7orf33→
←EZH2
←PDIA4
CUL1→
GHET1→
←ZNF786
←ZNF425
147
147.5
148
148.5
Position on chr7 (Mb)

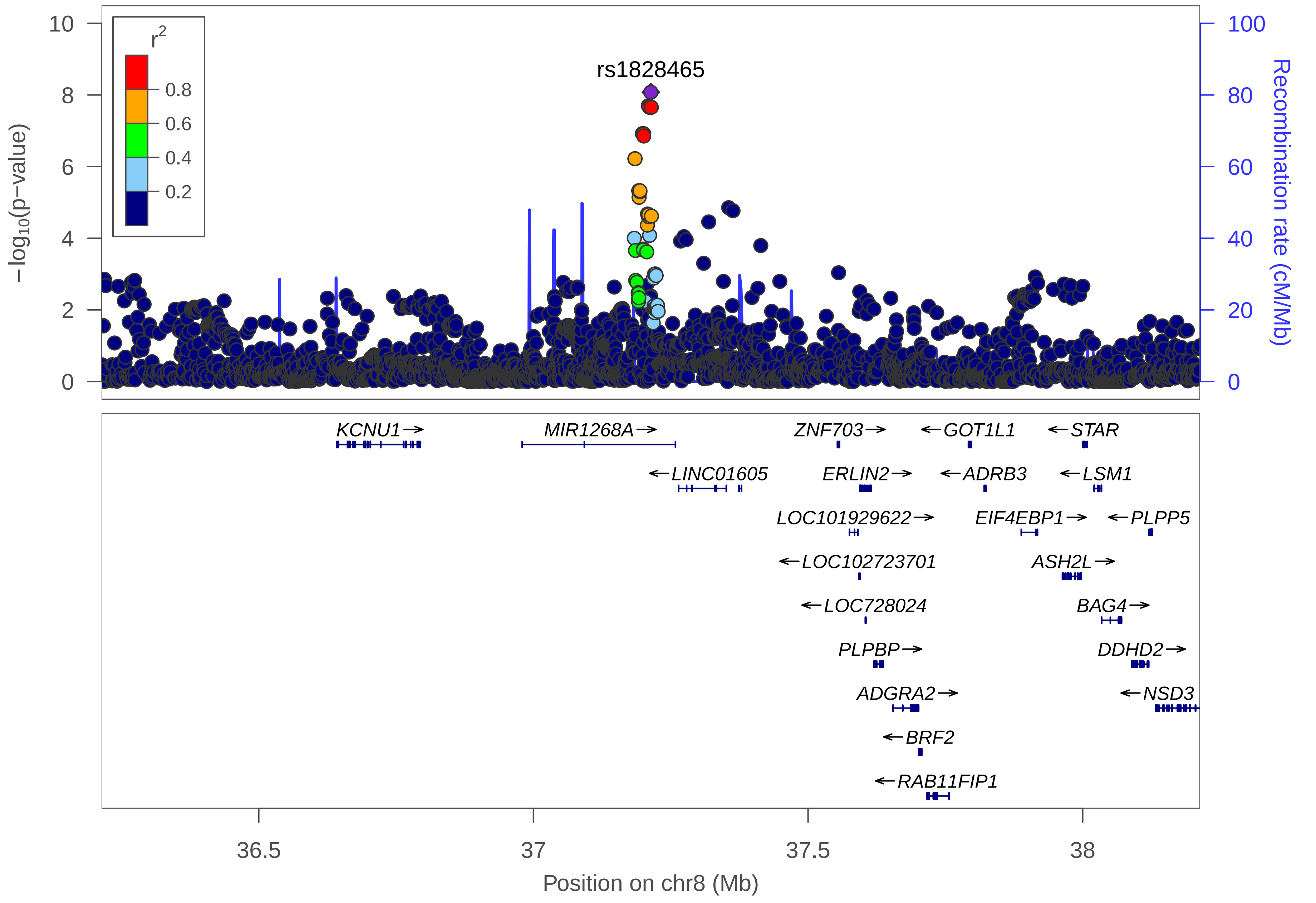
rs1828465
r2
0.8
0.6
0.4
0.2
−log10(p−value)
Recombination rate (cM/Mb)
10
8
6
4
2
0
100
80
60
40
20
0
KCNU1
MIR1268A
LINC01605
ZNF703
ERLIN2
LOC101929622
LOC102723701
LOC728024
PLPBP
ADGRA2
BRF2
RAB11FIP1
GOT1L1
ADRB3
EIF4EBP1
ASH2L
STAR
LSM1
PLPP5
BAG4
DDHD2
NSD3
36.5
37
37.5
38
Position on chr8 (Mb)

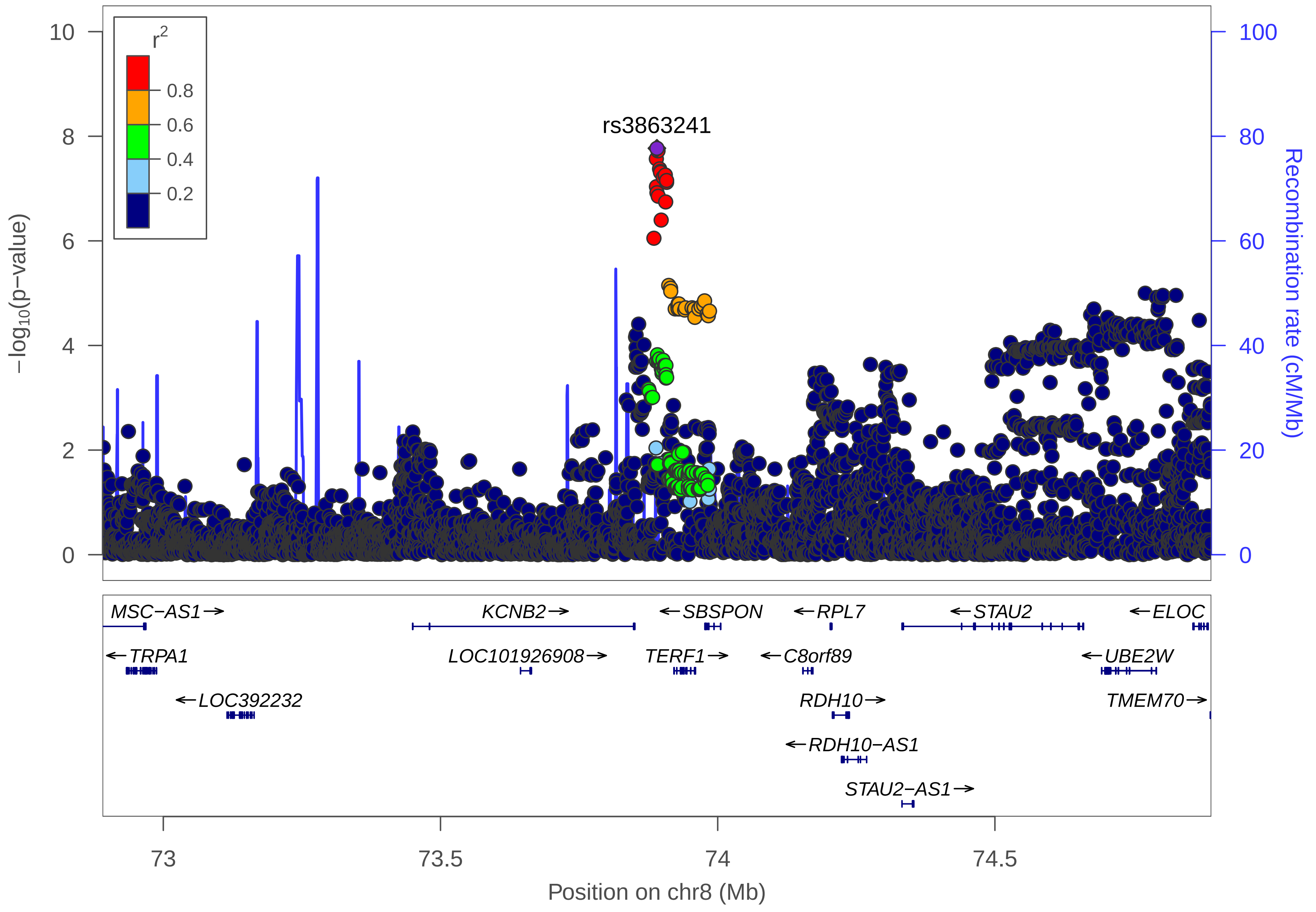

r2
0.8
0.6
0.4
0.2
rs3863241
10
8
6
4
2
0
−log10(p−value)
100
80
60
40
20
0
Recombination rate (cM/Mb)
MSC−AS1→
KCNB2→
←SBSPON
←RPL7
←STAU2
←ELOC
←TRPA1
LOC101926908→
TERF1→
←C8orf89
←UBE2W
←LOC392232
RDH10→
TMEM70→
←RDH10−AS1
STAU2−AS1→
73
73.5
74
74.5
Position on chr8 (Mb)

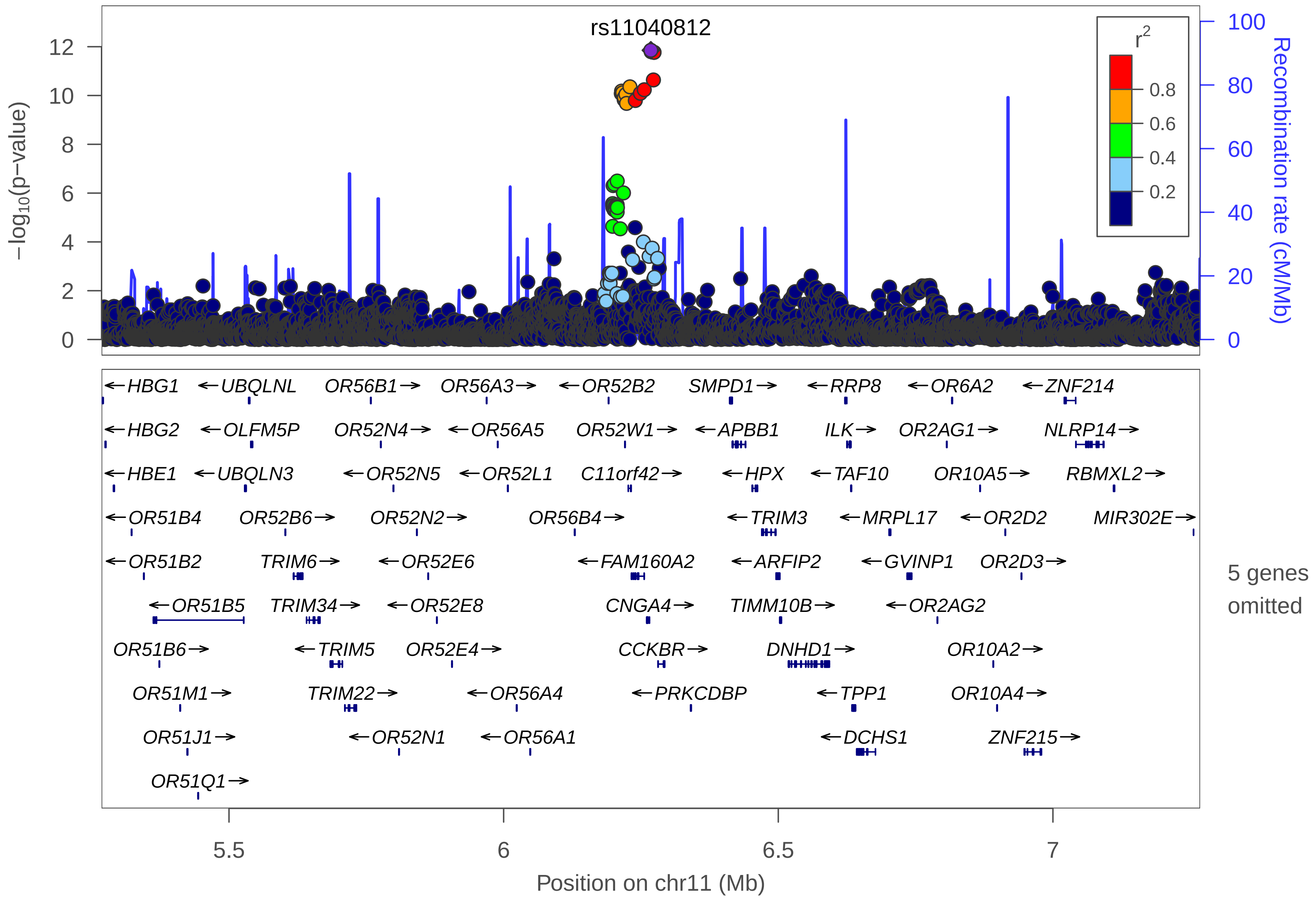
rs11040812
−log10(p−value)
Recombination rate (cM/Mb)
r2
0.8
0.6
0.4
0.2
0
2
4
6
8
10
12
20
40
60
80
100
←HBG1
←UBQLNL
OR56B1→
OR56A3→
←OR52B2
SMPD1→
←RRP8
←OR6A2
←ZNF214
←HBG2
←OLFM5P
OR52N4→
←OR56A5
OR52W1→
←APBB1
ILK→
OR2AG1→
NLRP14→
←HBE1
←UBQLN3
←OR52N5
←OR52L1
C11orf42→
←HPX
←TAF10
OR10A5→
RBMXL2→
←OR51B4
OR52B6→
OR52N2→
OR56B4→
←TRIM3
←MRPL17
←OR2D2
MIR302E→
←OR51B2
TRIM6→
←OR52E6
←FAM160A2
←ARFIP2
←GVINP1
OR2D3→
5 genes omitted
←OR51B5
TRIM34→
←OR52E8
CNGA4→
TIMM10B→
←OR2AG2
OR51B6→
←TRIM5
OR52E4→
CCKBR→
DNHD1→
OR10A2→
OR51M1→
TRIM22→
←OR56A4
←PRKCDBP
←TPP1
OR10A4→
OR51J1→
←OR52N1
←OR56A1
←DCHS1
ZNF215→
OR51Q1→
5.5
6
6.5
7
Position on chr11 (Mb)

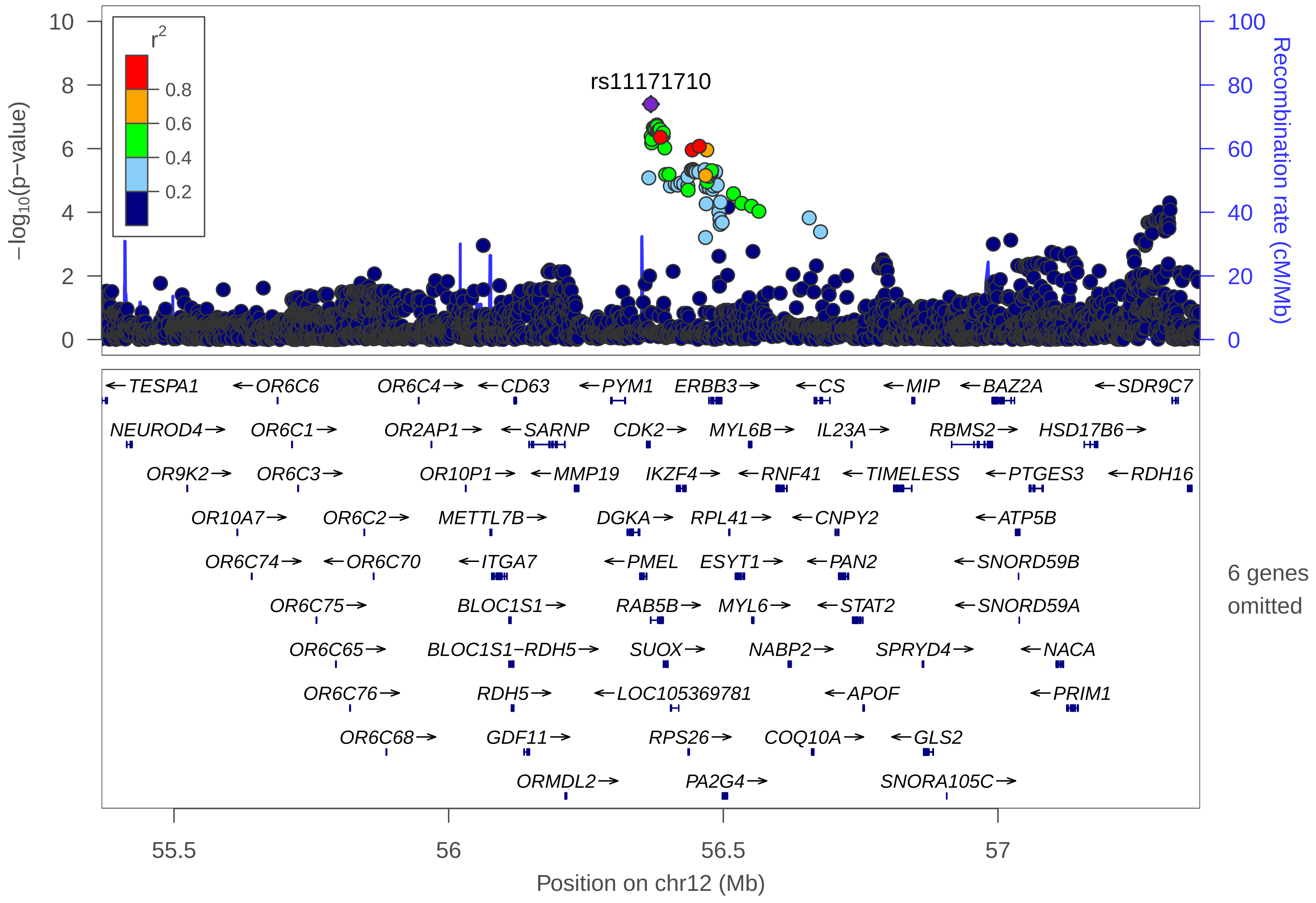

rs11171710
r2
0.8
0.6
0.4
0.2
10
8
6
4
2
0
100
80
60
40
20
0
−log10(p−value)
Recombination rate (cM/Mb)
←TESPA1
←OR6C6
OR6C4→
←CD63
←PYM1
ERBB3→
←CS
←MIP
←BAZ2A
←SDR9C7
NEUROD4→
OR6C1→
OR2AP1→
←SARNP
CDK2→
MYL6B→
IL23A→
RBMS2→
HSD17B6→
OR9K2→
OR6C3→
OR10P1→
←MMP19
IKZF4→
←RNF41
←TIMELESS
←PTGES3
←RDH16
OR10A7→
OR6C2→
METTL7B→
DGKA→
RPL41→
←CNPY2
←ATP5B
OR6C74→
←OR6C70
←ITGA7
←PMEL
ESYT1→
←PAN2
←SNORD59B
OR6C75→
BLOC1S1→
RAB5B→
MYL6→
←STAT2
←SNORD59A
OR6C65→
BLOC1S1−RDH5→
SUOX→
NABP2→
SPRYD4→
←NACA
OR6C76→
RDH5→
←LOC105369781
←APOF
←PRIM1
OR6C68→
GDF11→
RPS26→
COQ10A→
←GLS2
ORMDL2→
PA2G4→
SNORA105C→
6 genes omitted
55.5
56
56.5
57
Position on chr12 (Mb)

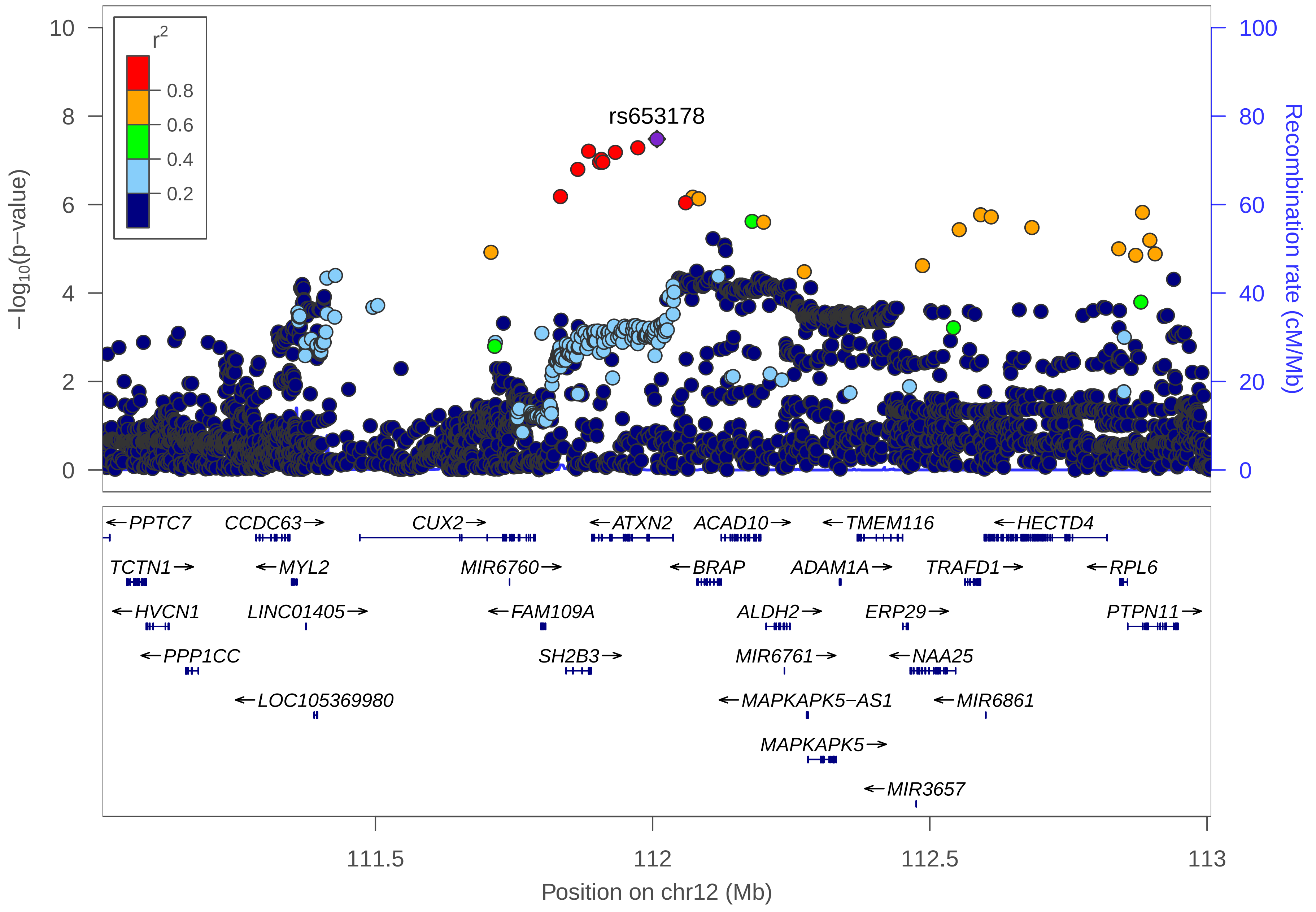

rs653178
$r^2$
0.8
0.6
0.4
0.2
10
8
6
4
2
0
−log10(p−value)
100
80
60
40
20
0
Recombination rate (cM/Mb)
←PPTC7
CCDC63→
CUX2→
←ATXN2
ACAD10→
←TMEM116
←HECTD4
TCTN1→
←MYL2
MIR6760→
←BRAP
ADAM1A→
TRAFD1→
←RPL6
←HVCN1
LINC01405→
←FAM109A
ALDH2→
ERP29→
PTPN11→
←PPP1CC
SH2B3→
MIR6761→
←NAA25
←LOC105369980
←MAPKAPK5−AS1
←MIR6861
MAPKAPK5→
←MIR3657
111.5
112
112.5
113
Position on chr12 (Mb)

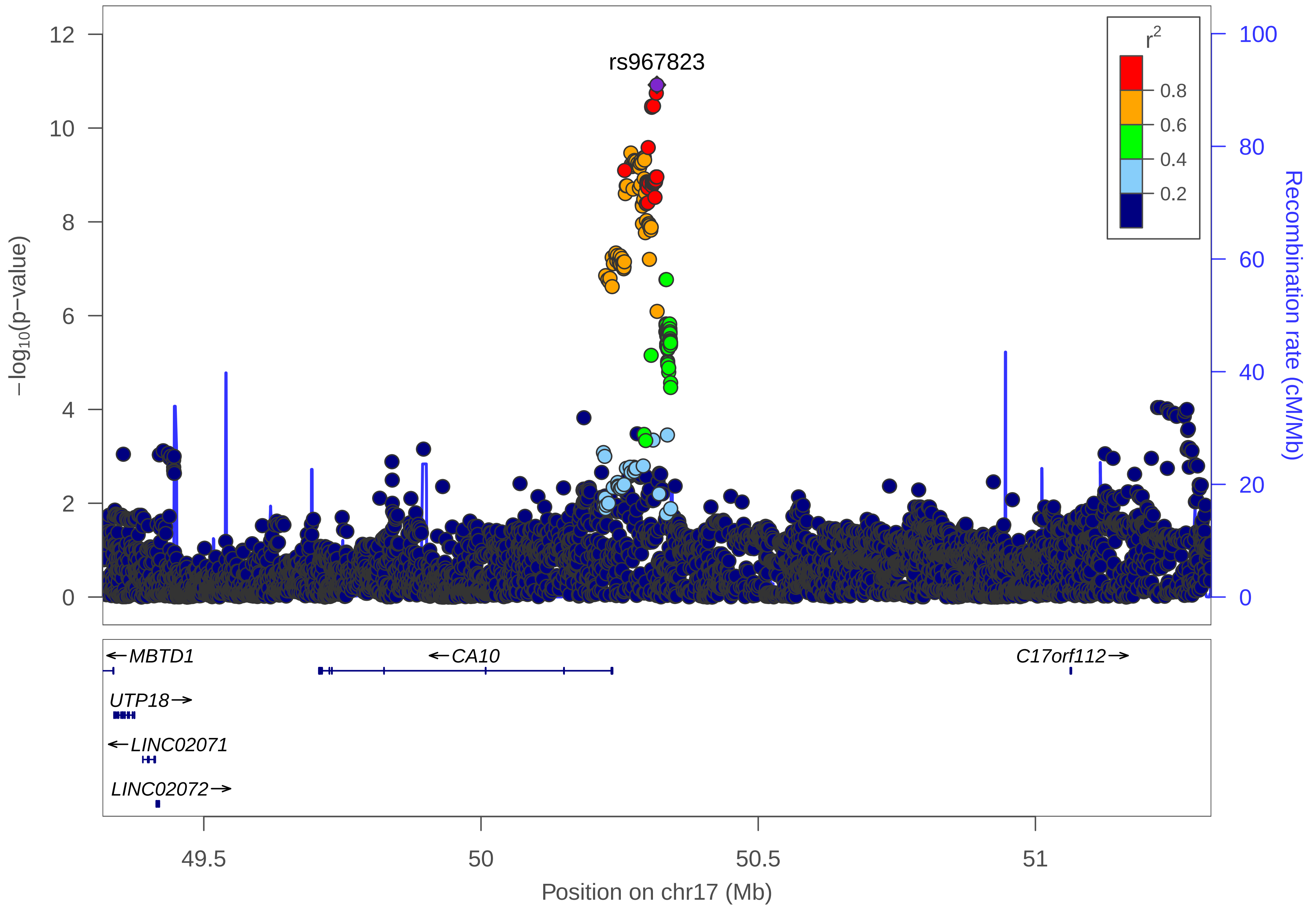

rs967823
r2
0.8
0.6
0.4
0.2
−log10(p−value)
Recombination rate (cM/Mb)
12
10
8
6
4
2
0
100
80
60
40
20
0
←MBTD1
←CA10
C17orf112→
UTP18→
←LINC02071
LINC02072→
49.5
50
50.5
51
Position on chr17 (Mb)

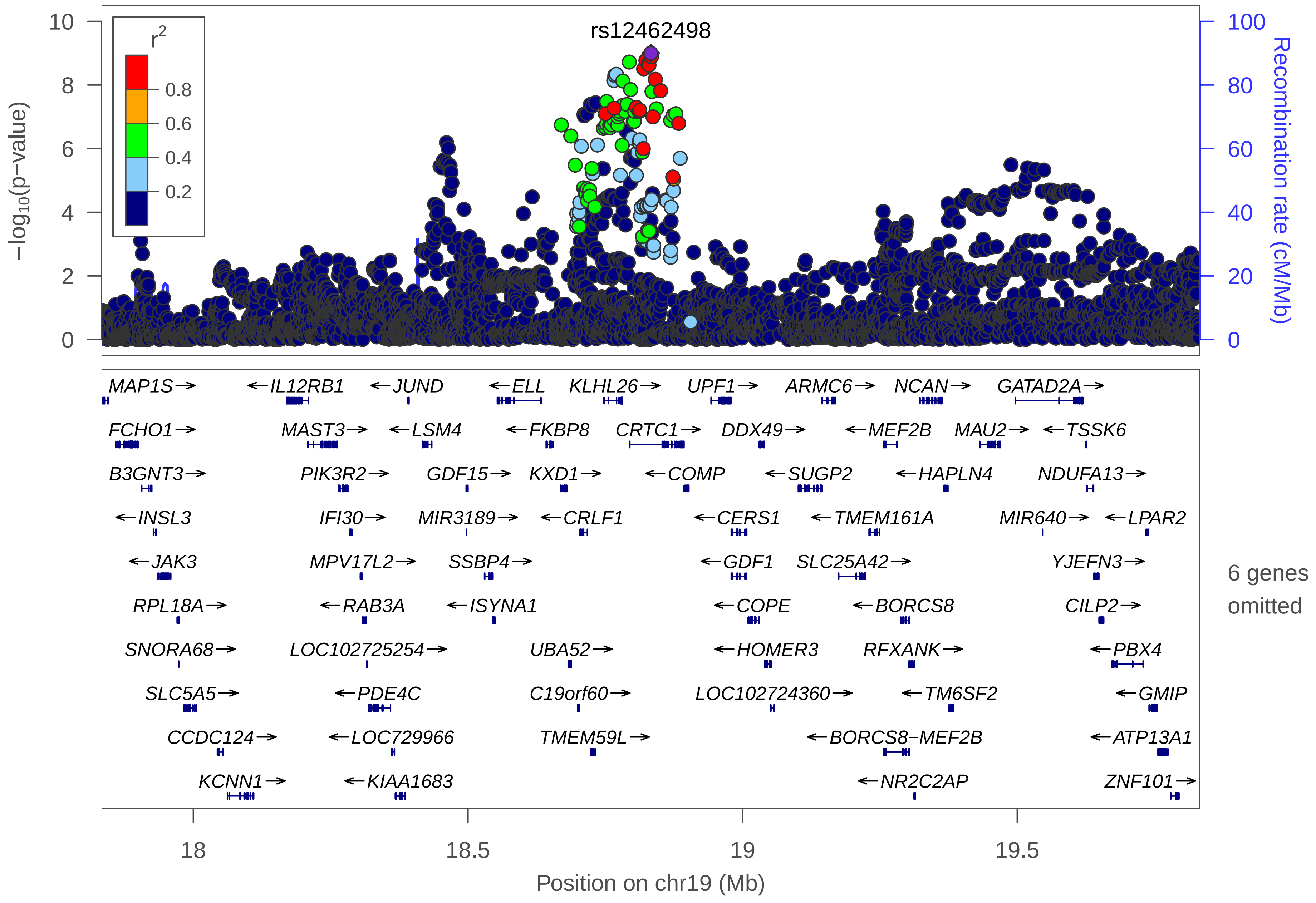
rs12462498
r2
0.8
0.6
0.4
0.2
−log10(p−value)
Recombination rate (cM/Mb)
0
2
4
6
8
10
0
20
40
60
80
100
MAP1S→
←IL12RB1
←JUND
←ELL
KLHL26→
UPF1→
ARMC6→
NCAN→
GATAD2A→
FCHO1→
MAST3→
←LSM4
←FKBP8
CRTC1→
DDX49→
←MEF2B
MAU2→
←TSSK6
B3GNT3→
PIK3R2→
GDF15→
KXD1→
←COMP
←SUGP2
←HAPLN4
NDUFA13→
←INSL3
IFI30→
MIR3189→
←CRLF1
←CERS1
←TMEM161A
MIR640→
←LPAR2
←JAK3
MPV17L2→
SSBP4→
←GDF1
SLC25A42→
YJEFN3→
RPL18A→
←RAB3A
←ISYNA1
←COPE
←BORCS8
CILP2→
SNORA68→
LOC102725254→
UBA52→
←HOMER3
RFXANK→
←PBX4
SLC5A5→
←PDE4C
C19orf60→
LOC102724360→
←TM6SF2
←GMIP
CCDC124→
←LOC729966
TMEM59L→
←BORCS8−MEF2B
←ATP13A1
KCNN1→
←KIAA1683
←NR2C2AP
ZNF101→
6 genes omitted
18
18.5
19
19.5
Position on chr19 (Mb)

### Irritable Bowel Syndrome and corresponding medications
### (IBS+M)

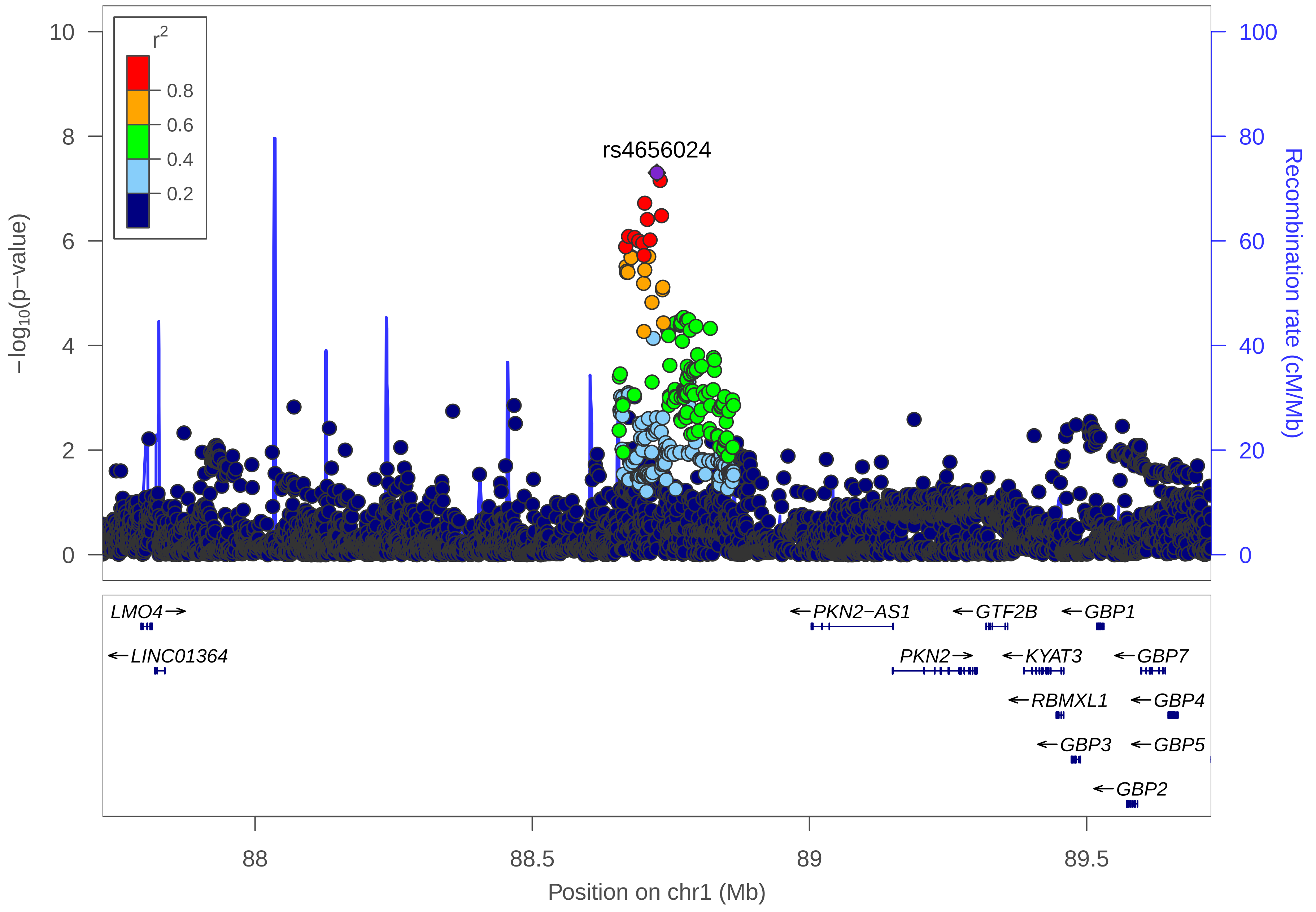

r2
0.8
0.6
0.4
0.2
rs4656024
−log10(p−value)
Recombination rate (cM/Mb)
0
2
4
6
8
10
0
20
40
60
80
100
LMO4
LINC01364
PKN2−AS1
PKN2
GTF2B
KYAT3
RBMXL1
GBP3
GBP2
GBP1
GBP7
GBP4
GBP5
88
88.5
89
89.5
Position on chr1 (Mb)

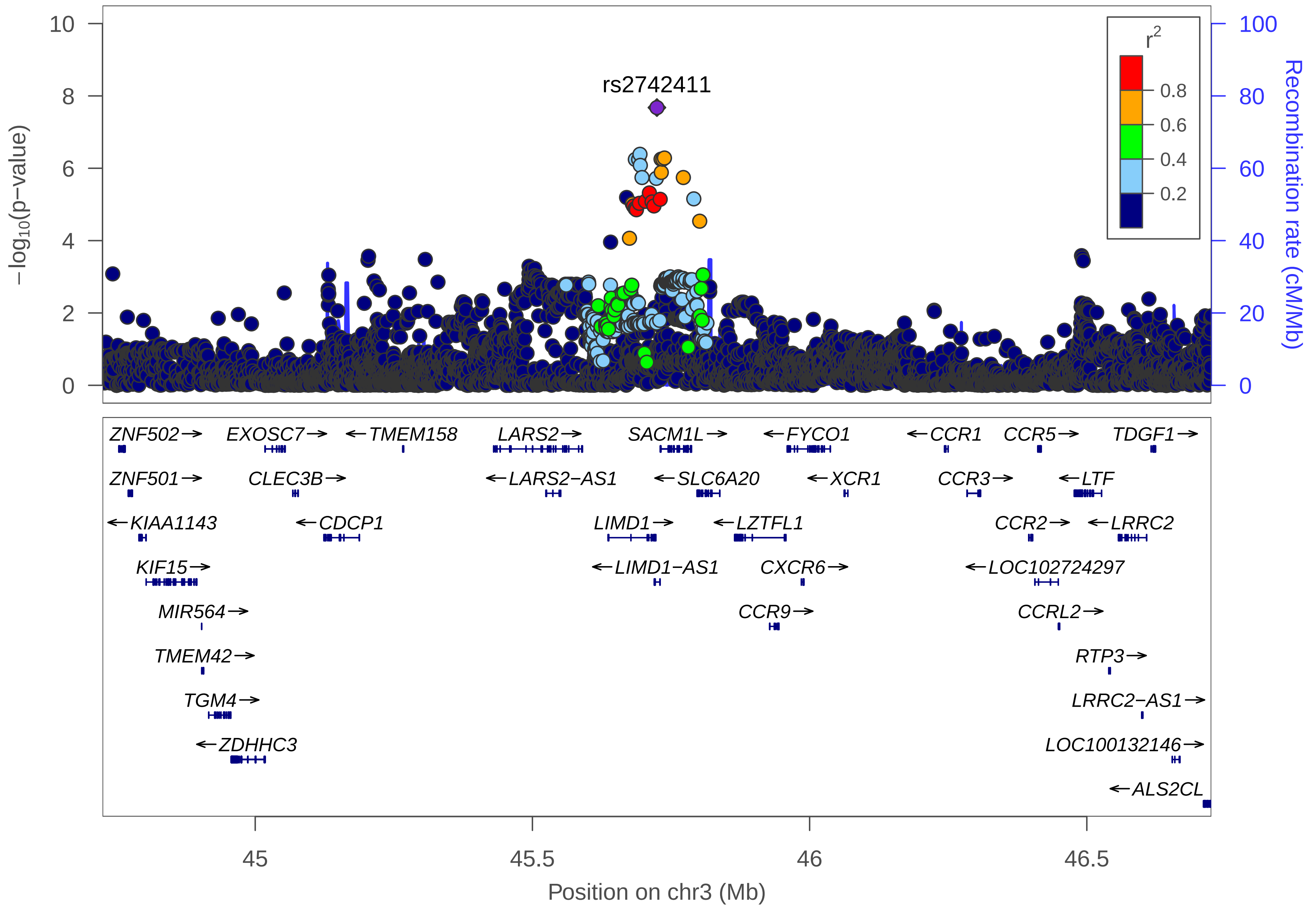

rs2742411
−log10(p−value)
Recombination rate (cM/Mb)
r2
0.8
0.6
0.4
0.2
0
2
4
6
8
10
0
20
40
60
80
100
ZNF502→
EXOSC7→
←TMEM158
LARS2→
SACM1L→
←FYCO1
←CCR1
CCR5→
TDGF1→
ZNF501→
CLEC3B→
←LARS2−AS1
←SLC6A20
←XCR1
CCR3→
←LTF
←KIAA1143
←CDCP1
LIMD1→
←LZTFL1
CCR2→
←LRRC2
KIF15→
←LIMD1−AS1
CXCR6→
←LOC102724297
MIR564→
CCR9→
CCRL2→
TMEM42→
RTP3→
TGM4→
LRRC2−AS1→
←ZDHHC3
LOC100132146→
←ALS2CL
45
45.5
46
46.5
Position on chr3 (Mb)

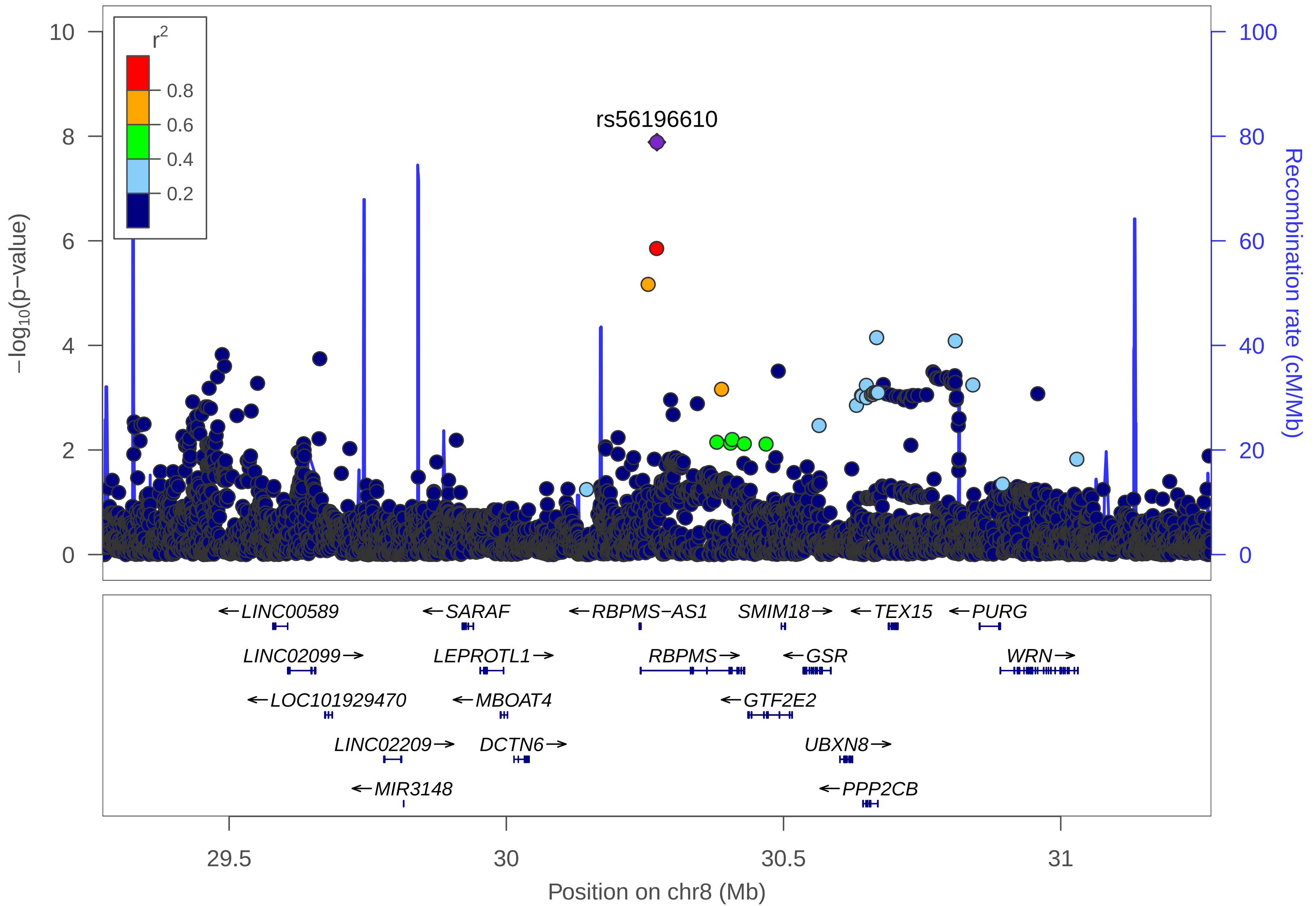
r2
0.8
0.6
0.4
0.2
rs56196610
−log10(p−value)
Recombination rate (cM/Mb)
0
2
4
6
8
10
0
20
40
60
80
100
←LINC00589
←SARAF
←RBPMS−AS1
SMIM18→
←TEX15
←PURG
LINC02099→
LEPROTL1→
RBPMS→
←GSR
WRN→
←LOC101929470
←MBOAT4
←GTF2E2
LINC02209→
DCTN6→
UBXN8→
←MIR3148
←PPP2CB
29.5
30
30.5
31
Position on chr8 (Mb)

### Inflammatory Bowel Disease (IBD)

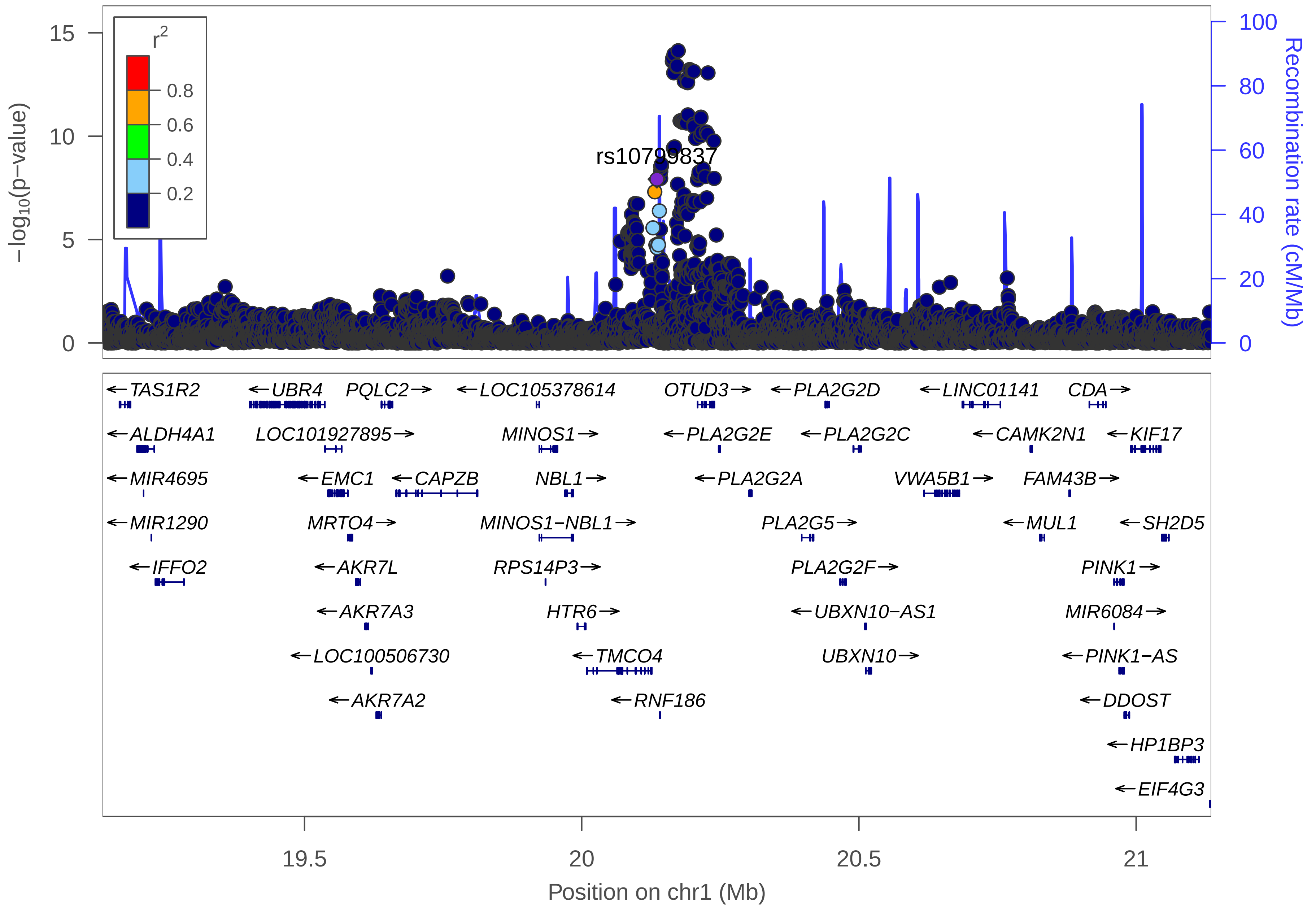

r2
0.8
0.6
0.4
0.2
15
10
5
0
−log10(p−value)
100
80
60
40
20
0
Recombination rate (cM/Mb)
rs10799837
TAS1R2
UBR4
PQLC2
LOC105378614
OTUD3
PLA2G2D
LINC01141
CDA
ALDH4A1
LOC101927895
MINOS1
PLA2G2E
PLA2G2C
CAMK2N1
KIF17
MIR4695
EMC1
CAPZB
NBL1
PLA2G2A
VWA5B1
FAM43B
MIR1290
MRTO4
MINOS1−NBL1
PLA2G5
MUL1
SH2D5
IFFO2
AKR7L
RPS14P3
PLA2G2F
PINK1
AKR7A3
HTR6
UBXN10−AS1
MIR6084
LOC100506730
TMCO4
UBXN10
PINK1−AS
AKR7A2
RNF186
DDOST
HP1BP3
EIF4G3
19.5
20
20.5
21
Position on chr1 (Mb)

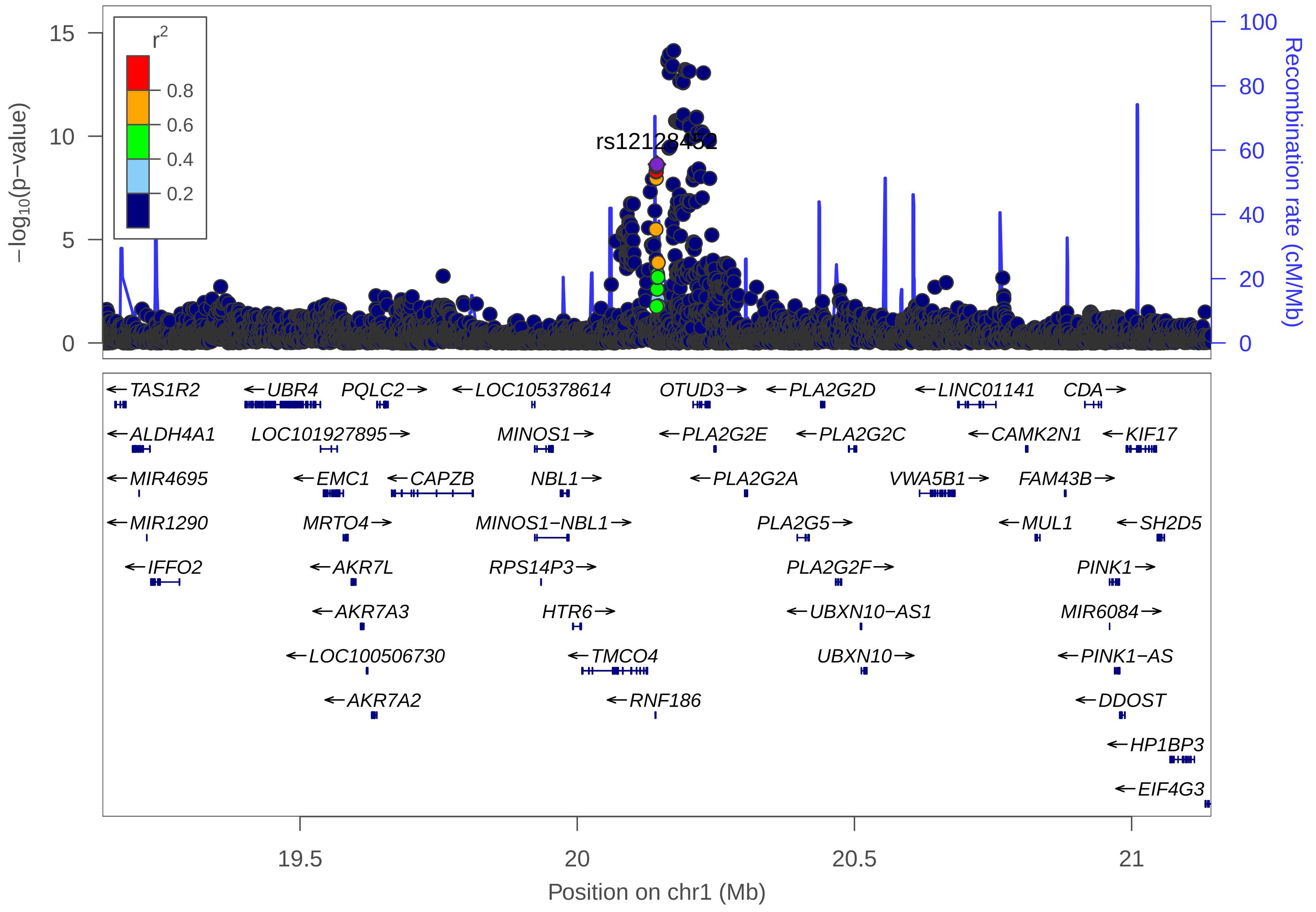

rs12128452
r2
0.8
0.6
0.4
0.2
15
10
5
0
100
80
60
40
20
0
−log10(p−value)
Recombination rate (cM/Mb)
TAS1R2
ALDH4A1
MIR4695
MIR1290
IFFO2
UBR4
LOC101927895
EMC1
MRTO4
AKR7L
AKR7A3
LOC100506730
AKR7A2
PQLC2
CAPZB
LOC105378614
MINOS1
NBL1
MINOS1−NBL1
RPS14P3
HTR6
TMCO4
RNF186
OTUD3
PLA2G2E
PLA2G2A
PLA2G2D
PLA2G2C
PLA2G5
PLA2G2F
UBXN10−AS1
UBXN10
LINC01141
CAMK2N1
VWA5B1
MUL1
CDA
KIF17
FAM43B
SH2D5
PINK1
MIR6084
PINK1−AS
DDOST
HP1BP3
EIF4G3
19.5
20
20.5
21
Position on chr1 (Mb)

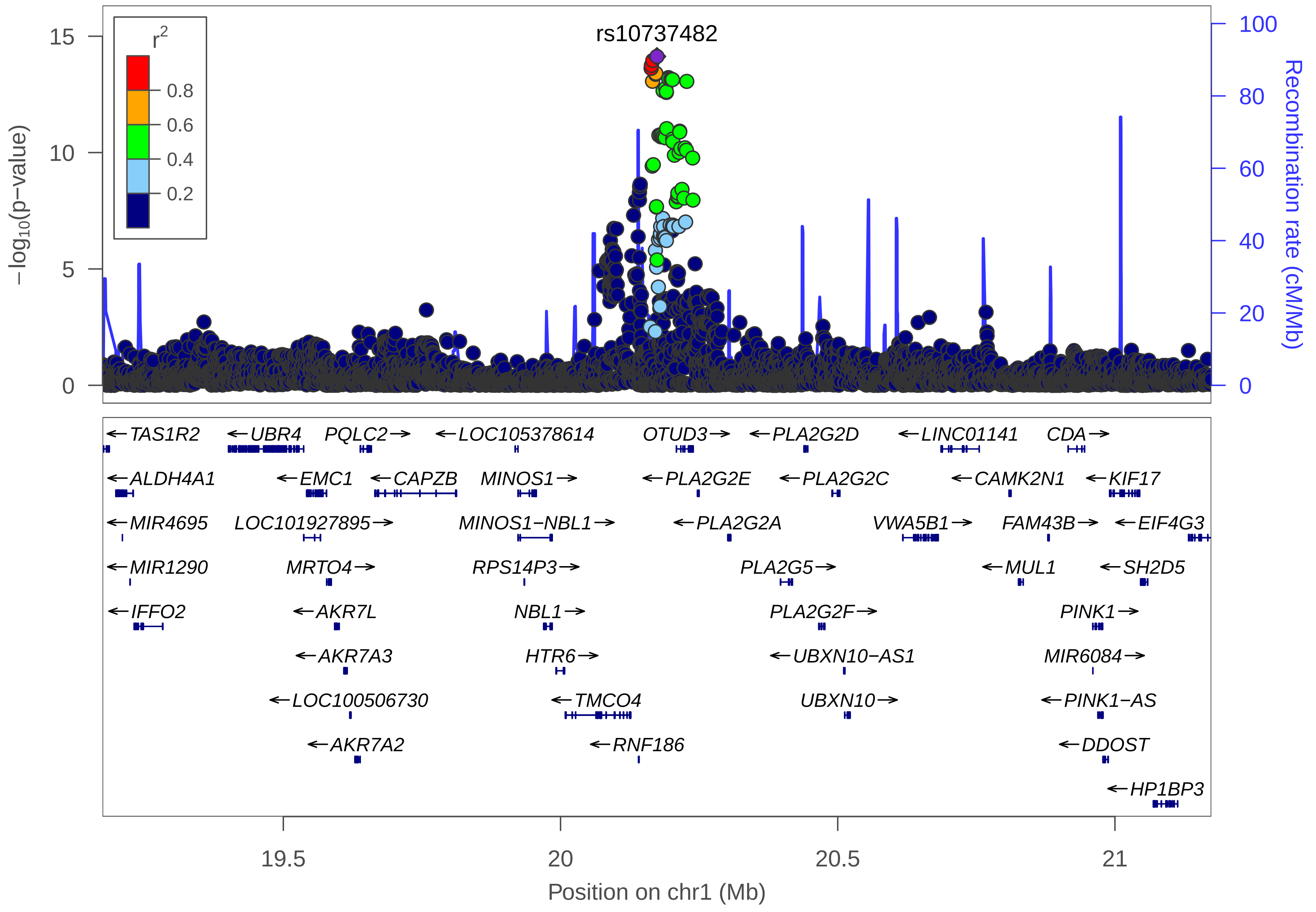

rs10737482
r2
0.8
0.6
0.4
0.2
15
10
5
0
100
80
60
40
20
0
−log10(p−value)
Recombination rate (cM/Mb)
←TAS1R2
←UBR4
PQLC2→
←LOC105378614
OTUD3→
←PLA2G2D
←LINC01141
CDA→
←ALDH4A1
←EMC1
←CAPZB
MINOS1→
←PLA2G2E
←PLA2G2C
←CAMK2N1
←KIF17
←MIR4695
LOC101927895→
MINOS1−NBL1→
←PLA2G2A
VWA5B1→
FAM43B→
←EIF4G3
←MIR1290
MRTO4→
RPS14P3→
PLA2G5→
←MUL1
←SH2D5
←IFFO2
←AKR7L
NBL1→
PLA2G2F→
PINK1→
←AKR7A3
HTR6→
←UBXN10−AS1
MIR6084→
←LOC100506730
←TMCO4
UBXN10→
←PINK1−AS
←AKR7A2
←RNF186
←DDOST
←HP1BP3
19.5
20
20.5
21
Position on chr1 (Mb)

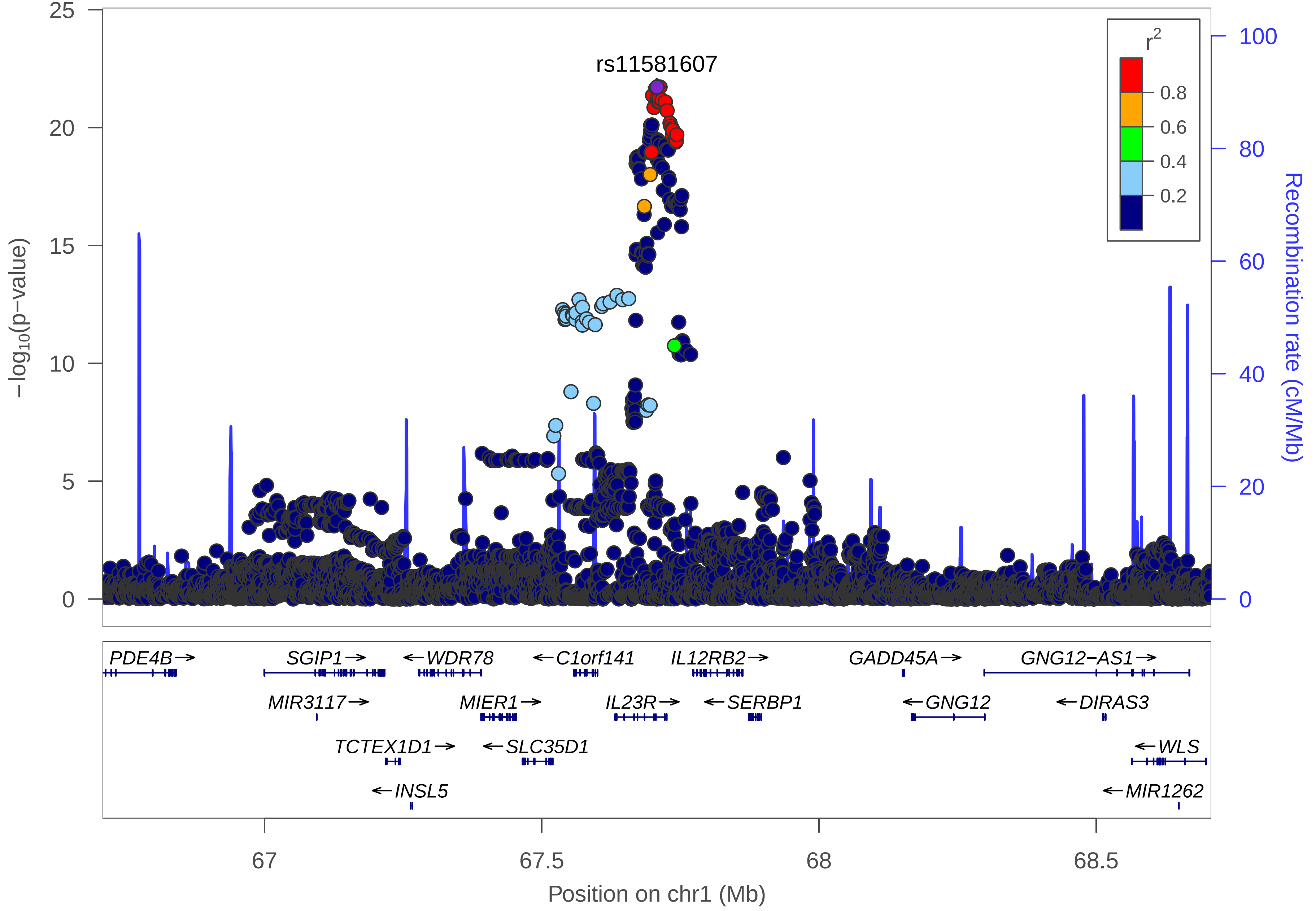

rs11581607
r²
0.8
0.6
0.4
0.2
−log10(p−value)
Recombination rate (cM/Mb)
25
20
15
10
5
0
100
80
60
40
20
0
PDE4B→
SGIP1→
←WDR78
←C1orf141
IL12RB2→
GADD45A→
GNG12−AS1→
MIR3117→
MIER1→
IL23R→
←SERBP1
←GNG12
←DIRAS3
TCTEX1D1→
←SLC35D1
←WLS
←INSL5
←MIR1262
67
67.5
68
68.5
Position on chr1 (Mb)

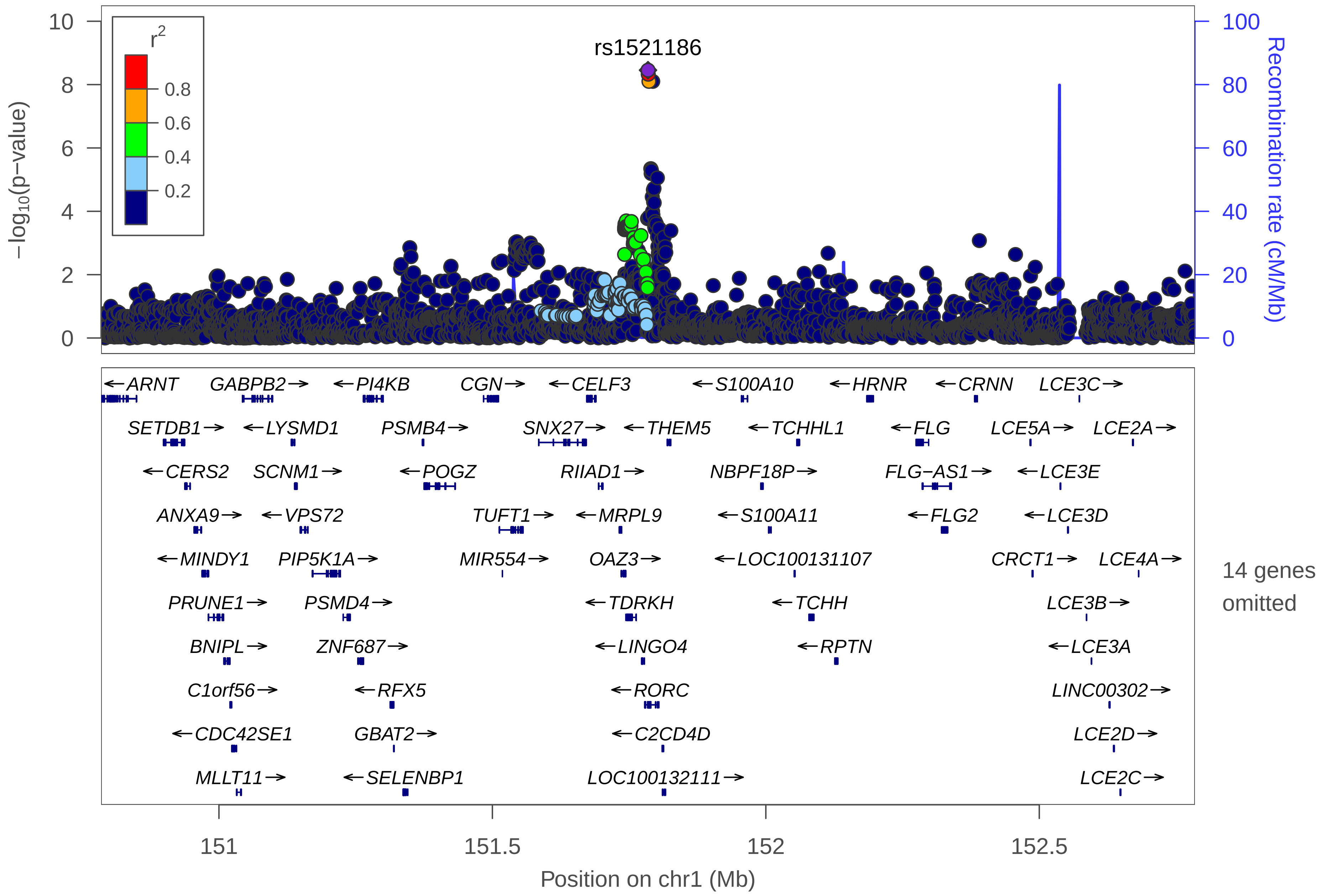

rs1521186
r2
0.8
0.6
0.4
0.2
−log10(p−value)
Recombination rate (cM/Mb)
0
2
4
6
8
10
20
40
60
80
100
ARNT
GABPB2
PI4KB
CGN
CELF3
S100A10
HRNR
CRNN
LCE3C
SETDB1
LYSMD1
PSMB4
SNX27
THEM5
TCHHL1
FLG
LCE5A
LCE2A
CERS2
SCNM1
POGZ
RIIAD1
NBPF18P
FLG−AS1
LCE3E
ANXA9
VPS72
TUFT1
MRPL9
S100A11
FLG2
LCE3D
MINDY1
PIP5K1A
MIR554
OAZ3
LOC100131107
CRCT1
LCE4A
PRUNE1
PSMD4
TDRKH
TCHH
LCE3B
BNIPL
ZNF687
LINGO4
RPTN
LCE3A
C1orf56
RFX5
RORC
LINC00302
CDC42SE1
GBAT2
C2CD4D
LCE2D
MLLT11
SELENBP1
LOC100132111
LCE2C
14 genes omitted
151
151.5
152
152.5
Position on chr1 (Mb)

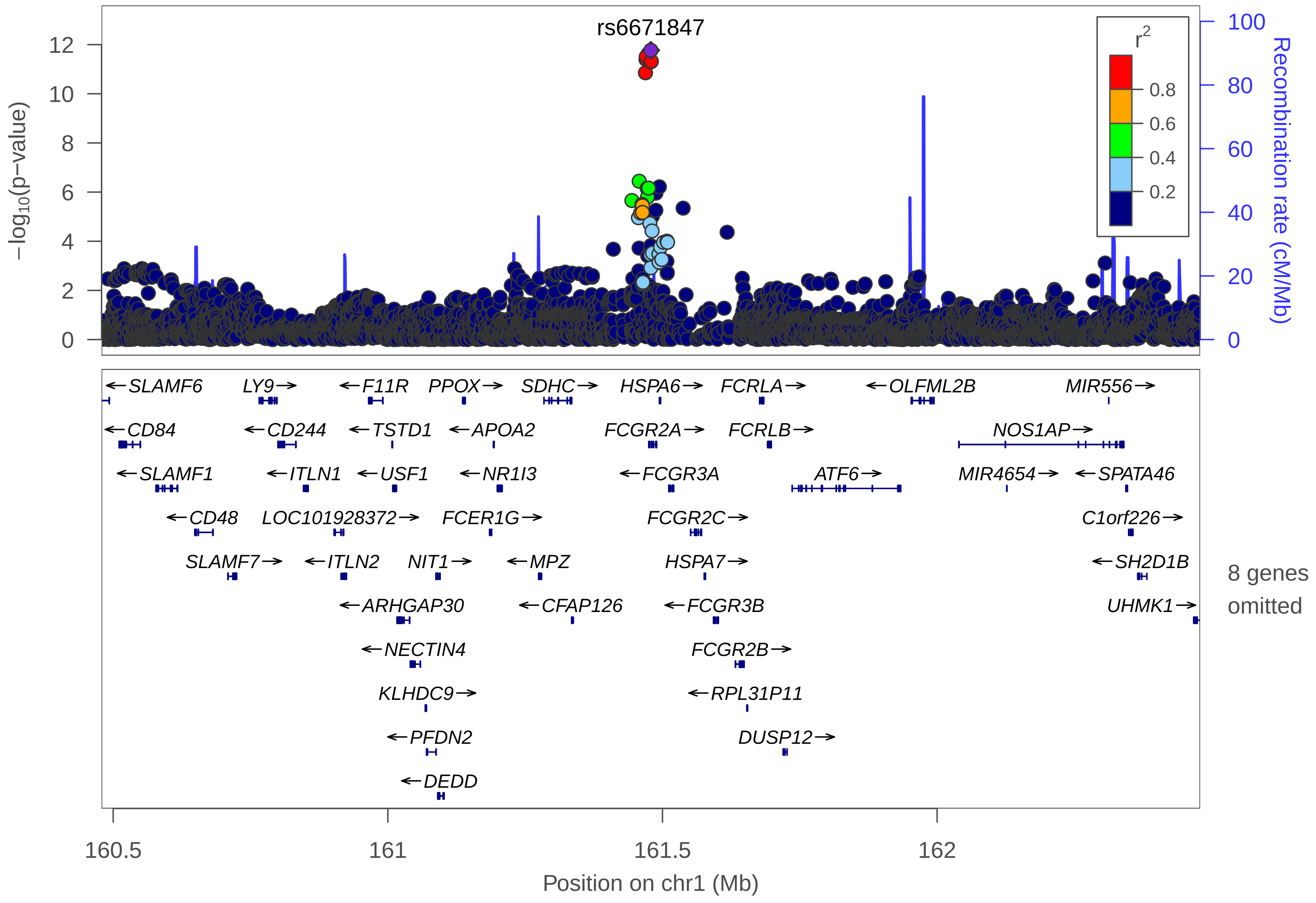

rs6671847
12
10
8
6
4
2
0
−log10(p−value)
100
80
60
40
20
0
Recombination rate (cM/Mb)
r2
0.8
0.6
0.4
0.2
←SLAMF6
LY9→
←F11R
PPOX→
SDHC→
HSPA6→
FCRLA→
←OLFML2B
MIR556→
←CD84
←CD244
←TSTD1
←APOA2
FCGR2A→
FCRLB→
NOS1AP→
←SLAMF1
←ITLN1
←USF1
←NR1I3
←FCGR3A
ATF6→
MIR4654→
←SPATA46
←CD48
LOC101928372→
FCER1G→
FCGR2C→
C1orf226→
SLAMF7→
←ITLN2
NIT1→
←MPZ
HSPA7→
←SH2D1B
8 genes omitted
←ARHGAP30
←CFAP126
←FCGR3B
UHMK1→
←NECTIN4
FCGR2B→
KLHDC9→
←RPL31P11
←PFDN2
DUSP12→
←DEDD
160.5
161
161.5
162
Position on chr1 (Mb)

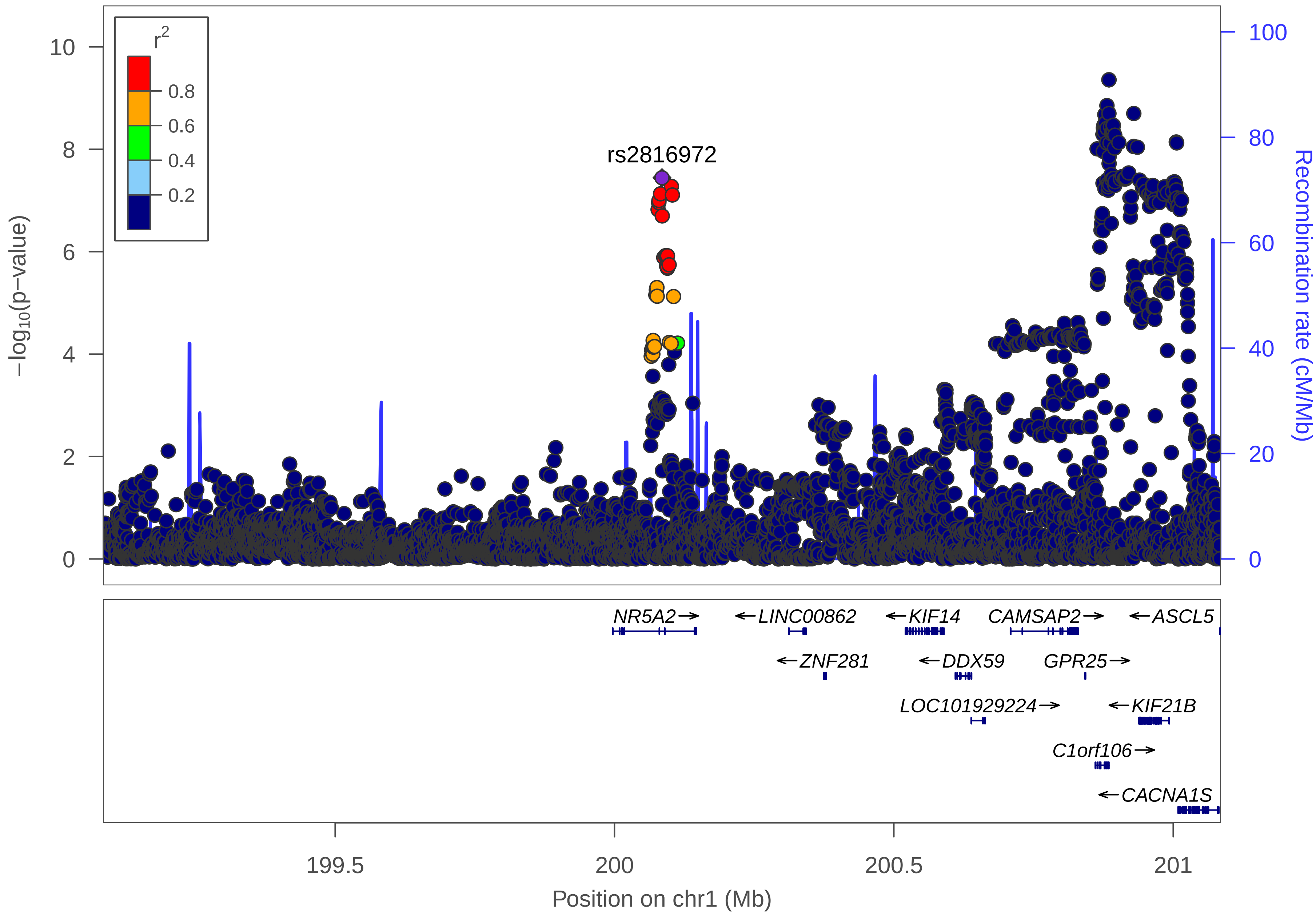

r²
0.8
0.6
0.4
0.2
rs2816972
−log10(p−value)
Recombination rate (cM/Mb)
0
2
4
6
8
10
0
20
40
60
80
100
NR5A2→
←LINC00862
←KIF14
CAMSAP2→
←ASCL5
←ZNF281
←DDX59
GPR25→
LOC101929224→
←KIF21B
C1orf106→
←CACNA1S
199.5
200
200.5
201
Position on chr1 (Mb)

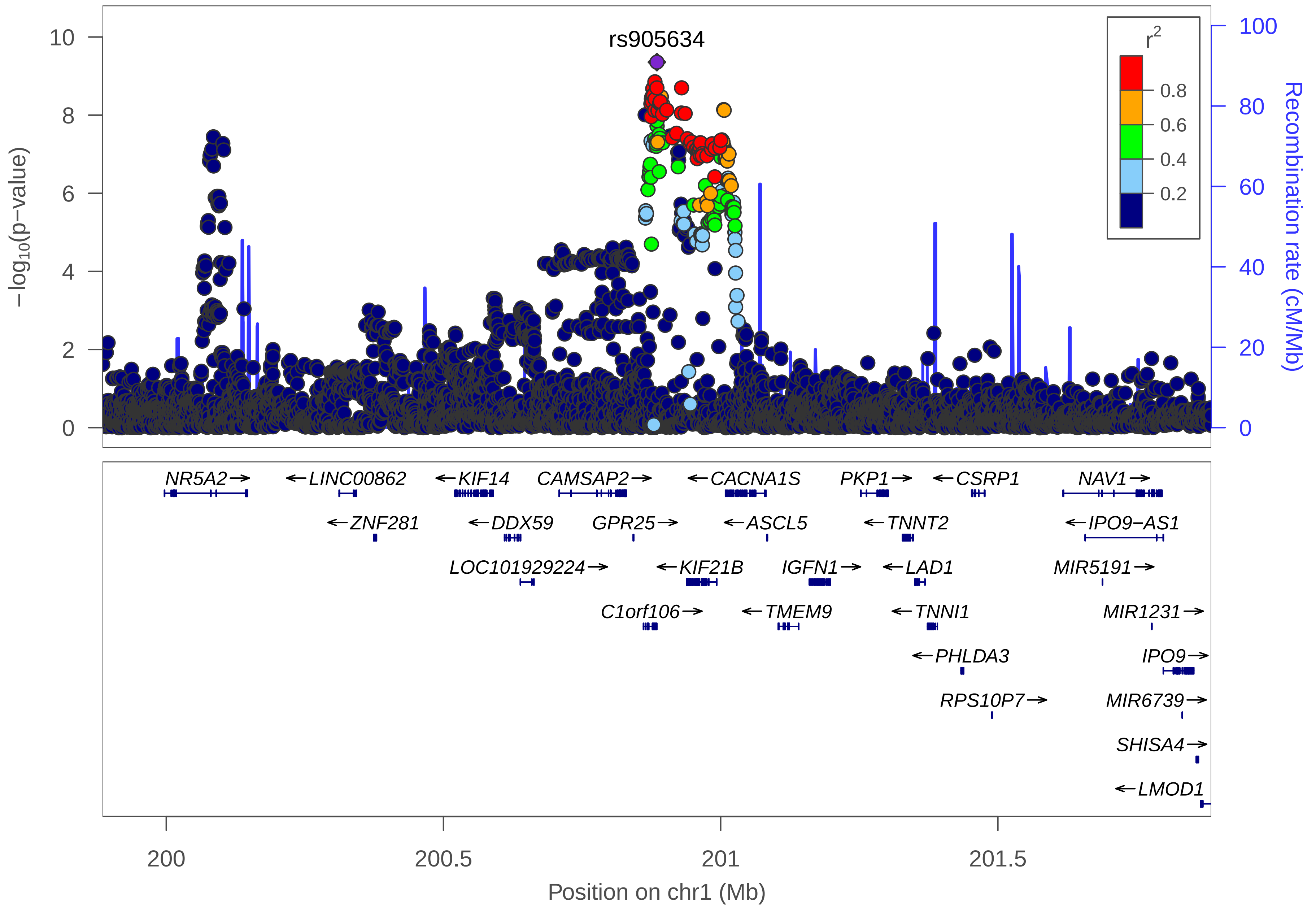

rs905634
r2
0.8
0.6
0.4
0.2
−log10(p−value)
Recombination rate (cM/Mb)
10
8
6
4
2
0
100
80
60
40
20
0
NR5A2→
←LINC00862
←KIF14
CAMSAP2→
←CACNA1S
PKP1→
←CSRP1
NAV1→
←ZNF281
←DDX59
GPR25→
←ASCL5
←TNNT2
←IPO9−AS1
LOC101929224→
←KIF21B
IGFN1→
←LAD1
MIR5191→
C1orf106→
←TMEM9
←TNNI1
MIR1231→
←PHLDA3
IPO9→
RPS10P7→
MIR6739→
SHISA4→
←LMOD1
200
200.5
201
201.5
Position on chr1 (Mb)

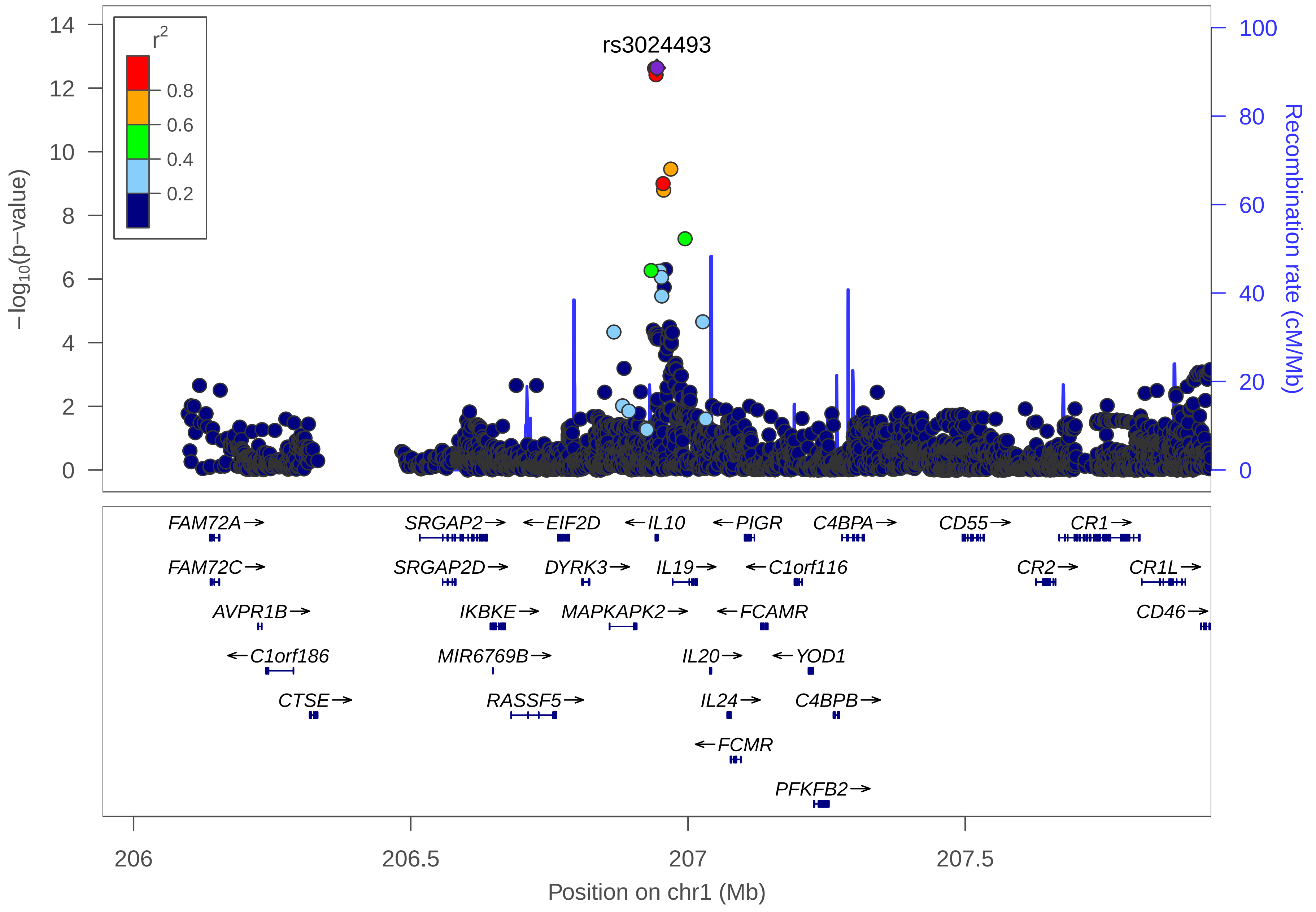

rs3024493
r2
0.8
0.6
0.4
0.2
−log10(p−value)
Recombination rate (cM/Mb)
FAM72A→
SRGAP2→
←EIF2D
←IL10
←PIGR
C4BPA→
CD55→
CR1→
FAM72C→
SRGAP2D→
DYRK3→
IL19→
←C1orf116
CR2→
CR1L→
AVPR1B→
IKBKE→
MAPKAPK2→
←FCAMR
CD46→
←C1orf186
MIR6769B→
IL20→
←YOD1
CTSE→
RASSF5→
IL24→
C4BPB→
←FCMR
PFKFB2→
206
206.5
207
207.5
Position on chr1 (Mb)

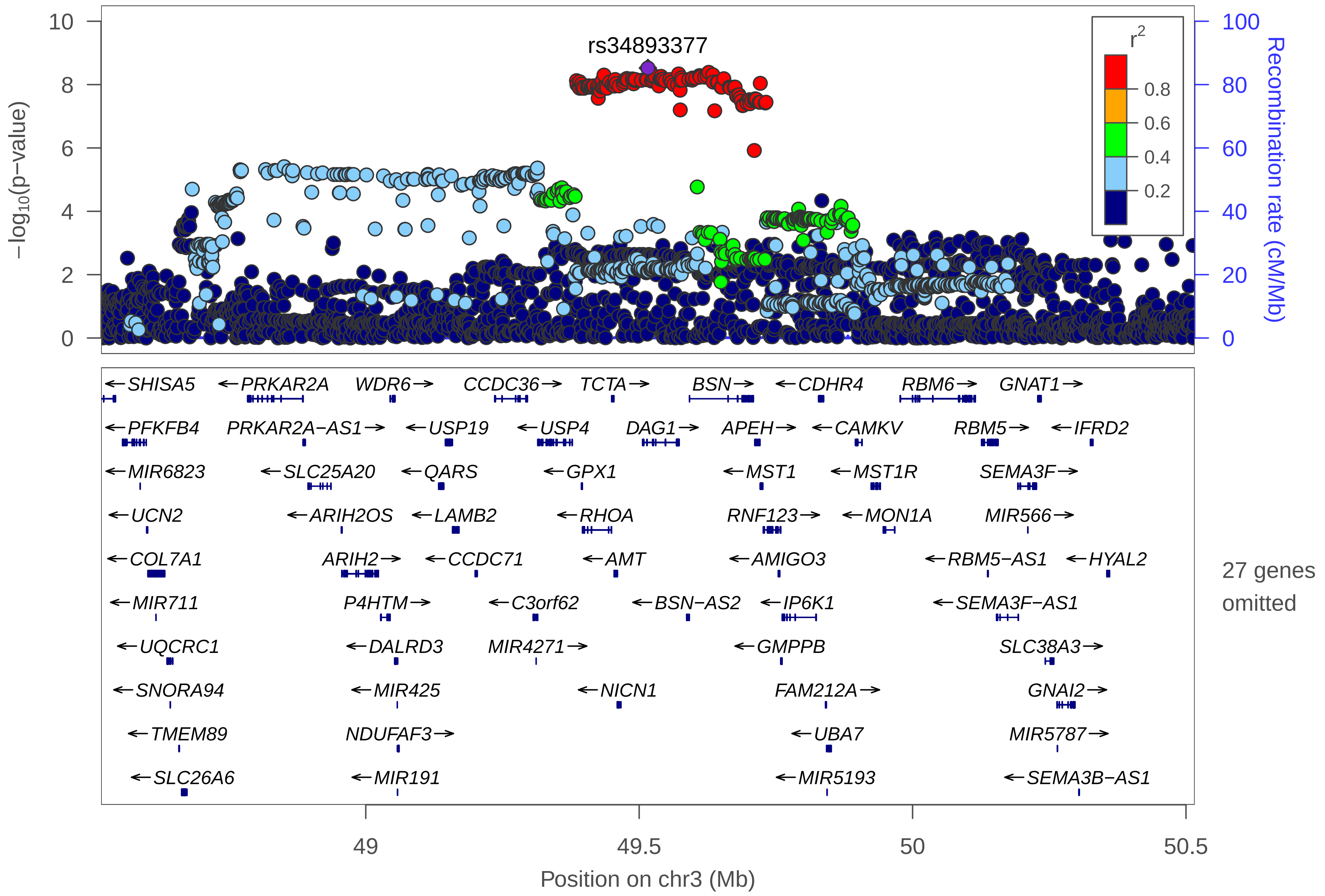

rs34893377
r²
0.8
0.6
0.4
0.2
10
8
6
4
2
0
−log10(p−value)
100
80
60
40
20
0
Recombination rate (cM/Mb)
←SHISA5
←PRKAR2A
WDR6→
CCDC36→
TCTA→
BSN→
←CDHR4
RBM6→
GNAT1→
←PFKFB4
PRKAR2A−AS1→
←USP19
←USP4
DAG1→
APEH→
←CAMKV
RBM5→
←IFRD2
←MIR6823
←SLC25A20
←QARS
←GPX1
←MST1
←MST1R
SEMA3F→
←UCN2
←ARIH2OS
←LAMB2
←RHOA
RNF123→
←MON1A
MIR566→
←COL7A1
ARIH2→
←CCDC71
←AMT
←AMIGO3
←RBM5−AS1
←HYAL2
←MIR711
P4HTM→
←C3orf62
←BSN−AS2
←IP6K1
←SEMA3F−AS1
←UQCRC1
←DALRD3
MIR4271→
←GMPPB
SLC38A3→
←SNORA94
←MIR425
←NICN1
FAM212A→
GNAI2→
←TMEM89
NDUFAF3→
←UBA7
MIR5787→
←SLC26A6
←MIR191
←MIR5193
←SEMA3B−AS1
27 genes omitted
49
49.5
50
50.5
Position on chr3 (Mb)

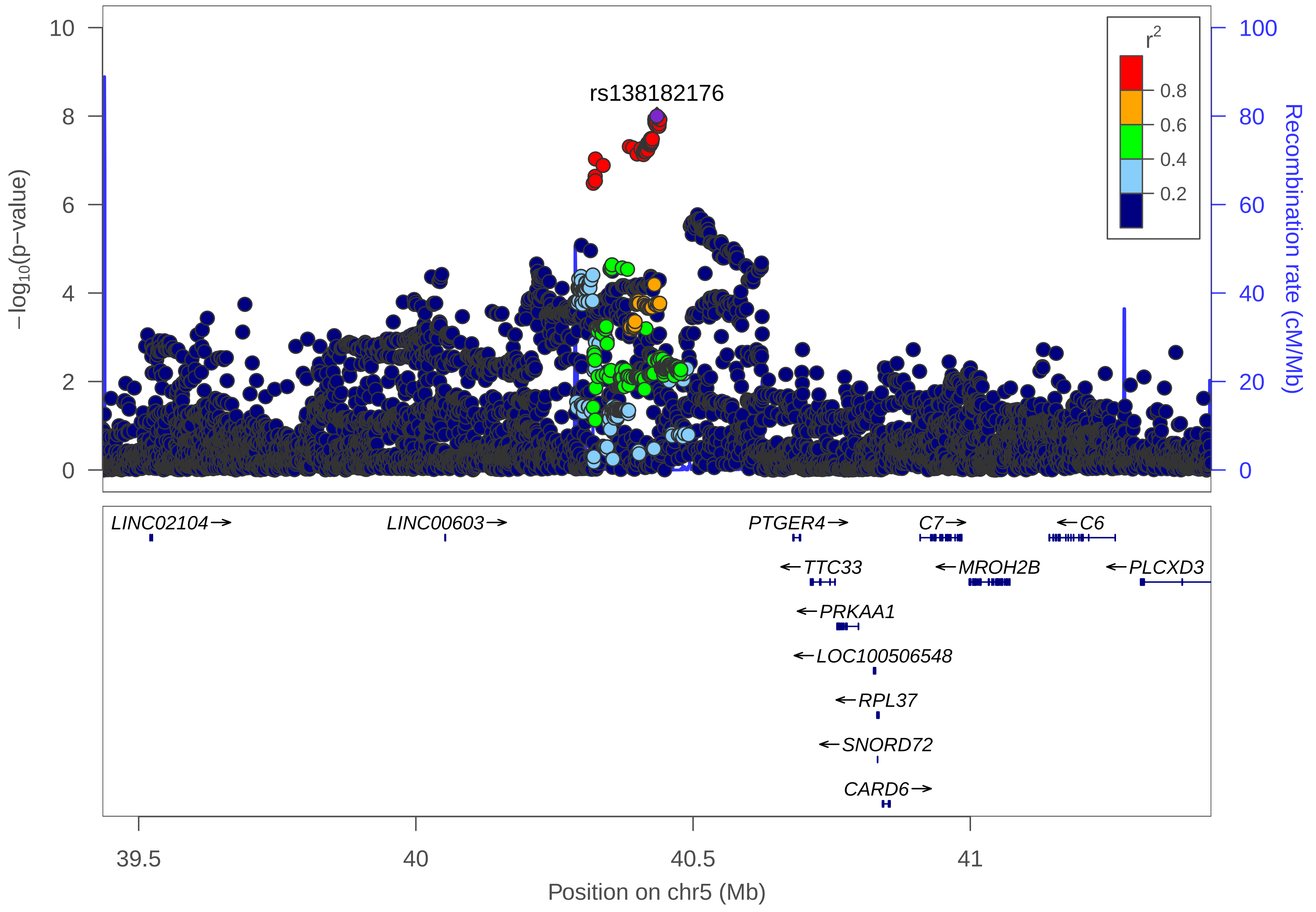

rs138182176
r2
0.8
0.6
0.4
0.2
−log10(p−value)
Recombination rate (cM/Mb)
0
2
4
6
8
10
0
20
40
60
80
100
LINC02104→
LINC00603→
PTGER4→
C7→
←C6
←TTC33
←MROH2B
←PLCXD3
←PRKAA1
←LOC100506548
←RPL37
←SNORD72
CARD6→
39.5
40
40.5
41
Position on chr5 (Mb)

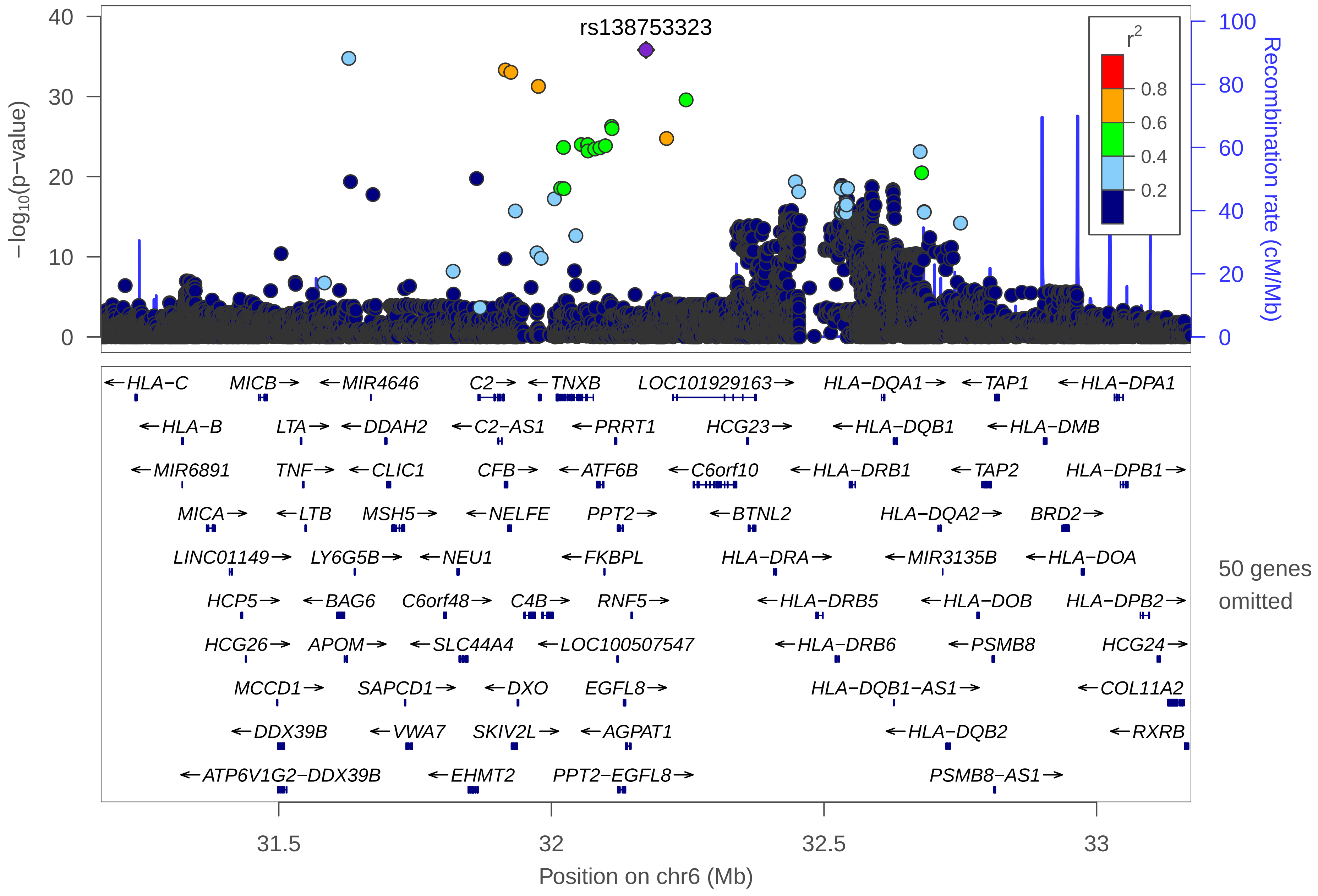
rs138753323
r2
0.8
0.6
0.4
0.2
−log10(p−value)
Recombination rate (cM/Mb)
←HLA−C
MICB→
←MIR4646
C2→
←TNXB
LOC101929163→
HLA−DQA1→
←TAP1
←HLA−DPA1
←HLA−B
LTA→
←DDAH2
←C2−AS1
←PRRT1
HCG23→
←HLA−DQB1
←HLA−DMB
←MIR6891
TNF→
←CLIC1
CFB→
←ATF6B
←C6orf10
←HLA−DRB1
←TAP2
HLA−DPB1→
MICA→
←LTB
MSH5→
←NELFE
PPT2→
←BTNL2
HLA−DQA2→
BRD2→
LINC01149→
LY6G5B→
←NEU1
←FKBPL
HLA−DRA→
←MIR3135B
←HLA−DOA
50 genes omitted
HCP5→
←BAG6
C6orf48→
C4B→
RNF5→
←HLA−DRB5
←HLA−DOB
HLA−DPB2→
HCG26→
APOM→
←SLC44A4
←LOC100507547
←HLA−DRB6
←PSMB8
HCG24→
MCCD1→
SAPCD1→
←DXO
EGFL8→
HLA−DQB1−AS1→
←COL11A2
←DDX39B
←VWA7
SKIV2L→
←AGPAT1
←HLA−DQB2
←RXRB
←ATP6V1G2−DDX39B
←EHMT2
PPT2−EGFL8→
PSMB8−AS1→
31.5
32
32.5
33
Position on chr6 (Mb)

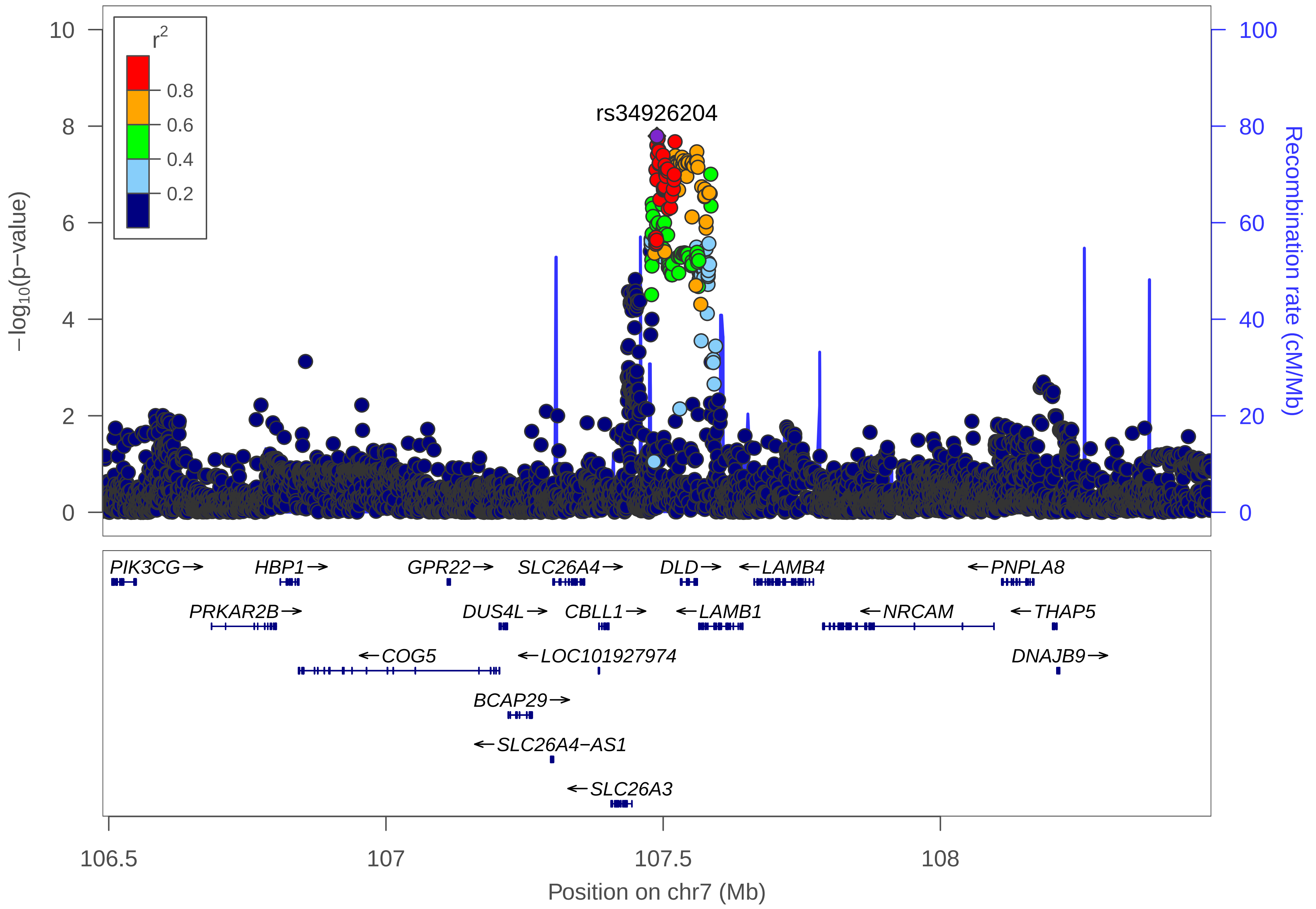

rs34926204
$-\log_{10}$(p−value)
Recombination rate (cM/Mb)
$r^2$
0.8
0.6
0.4
0.2
10
8
6
4
2
0
100
80
60
40
20
0
PIK3CG→
HBP1→
GPR22→
SLC26A4→
DLD→
←LAMB4
←PNPLA8
PRKAR2B→
DUS4L→
CBLL1→
←LAMB1
←NRCAM
←THAP5
←COG5
←LOC101927974
DNAJB9→
BCAP29→
←SLC26A4−AS1
←SLC26A3
106.5
107
107.5
108
Position on chr7 (Mb)

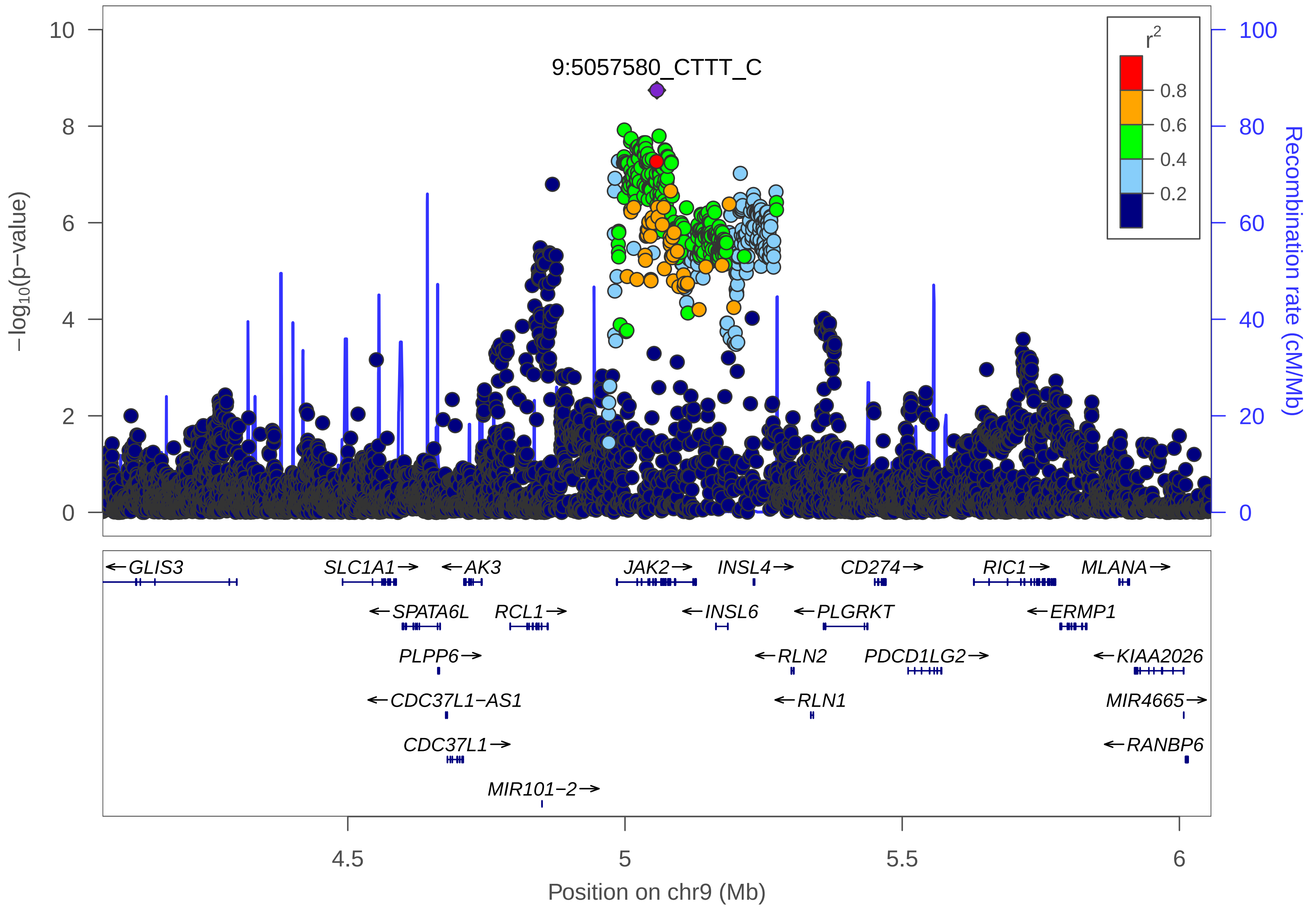

9:5057580_CTTT_C
r2
0.8
0.6
0.4
0.2
−log10(p−value)
Recombination rate (cM/Mb)
10
8
6
4
2
0
100
80
60
40
20
0
GLIS3
SLC1A1
AK3
JAK2
INSL4
CD274
RIC1
MLANA
SPATA6L
RCL1
INSL6
PLGRKT
ERMP1
PLPP6
RLN2
PDCD1LG2
KIAA2026
CDC37L1−AS1
RLN1
MIR4665
CDC37L1
RANBP6
MIR101−2
4.5
5
5.5
6
Position on chr9 (Mb)

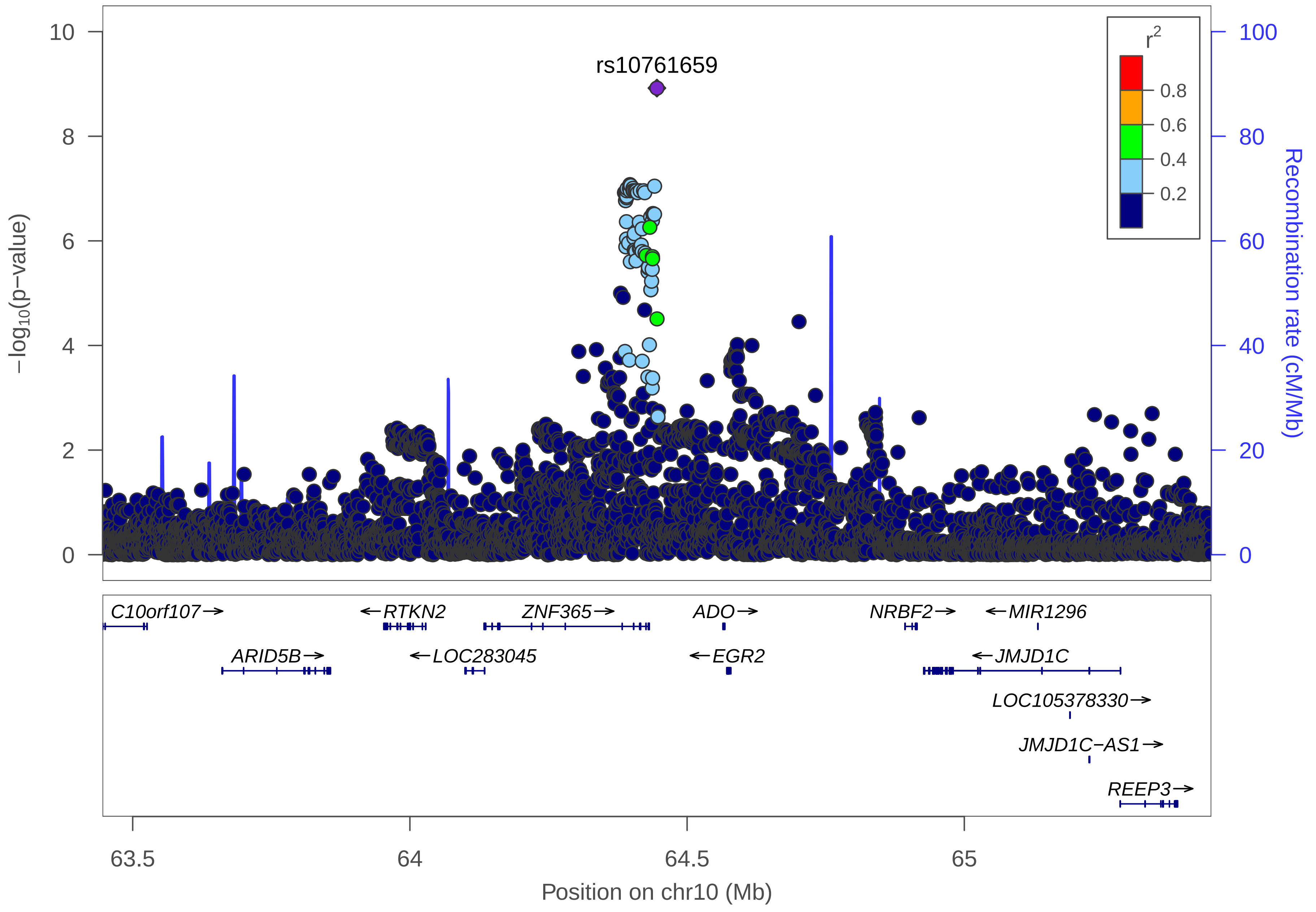
rs10761659
r2
0.8
0.6
0.4
0.2
−log10(p−value)
Recombination rate (cM/Mb)
C10orf107
RTKN2
ZNF365
ADO
NRBF2
MIR1296
ARID5B
LOC283045
EGR2
JMJD1C
LOC105378330
JMJD1C−AS1
REEP3
63.5
64
64.5
65
Position on chr10 (Mb)

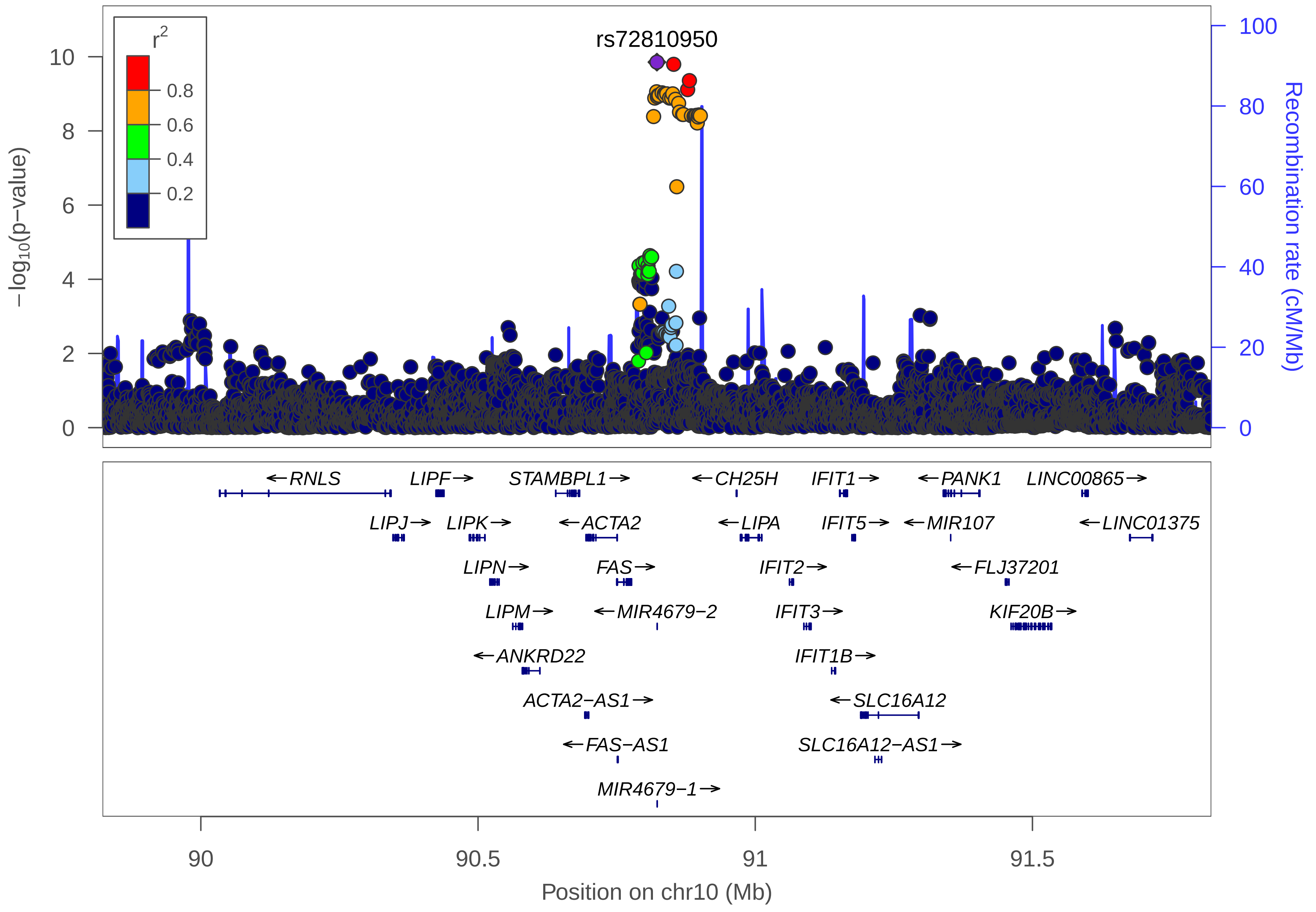

rs72810950
r2
0.8
0.6
0.4
0.2
10
8
6
4
2
0
−log10(p−value)
100
80
60
40
20
0
Recombination rate (cM/Mb)
←RNLS
LIPF→
STAMBPL1→
←CH25H
IFIT1→
←PANK1
LINC00865→
LIPJ→
LIPK→
←ACTA2
←LIPA
IFIT5→
←MIR107
←LINC01375
LIPN→
FAS→
IFIT2→
←FLJ37201
LIPM→
←MIR4679−2
IFIT3→
KIF20B→
←ANKRD22
IFIT1B→
ACTA2−AS1→
←SLC16A12
←FAS−AS1
SLC16A12−AS1→
MIR4679−1→
90
90.5
91
91.5
Position on chr10 (Mb)

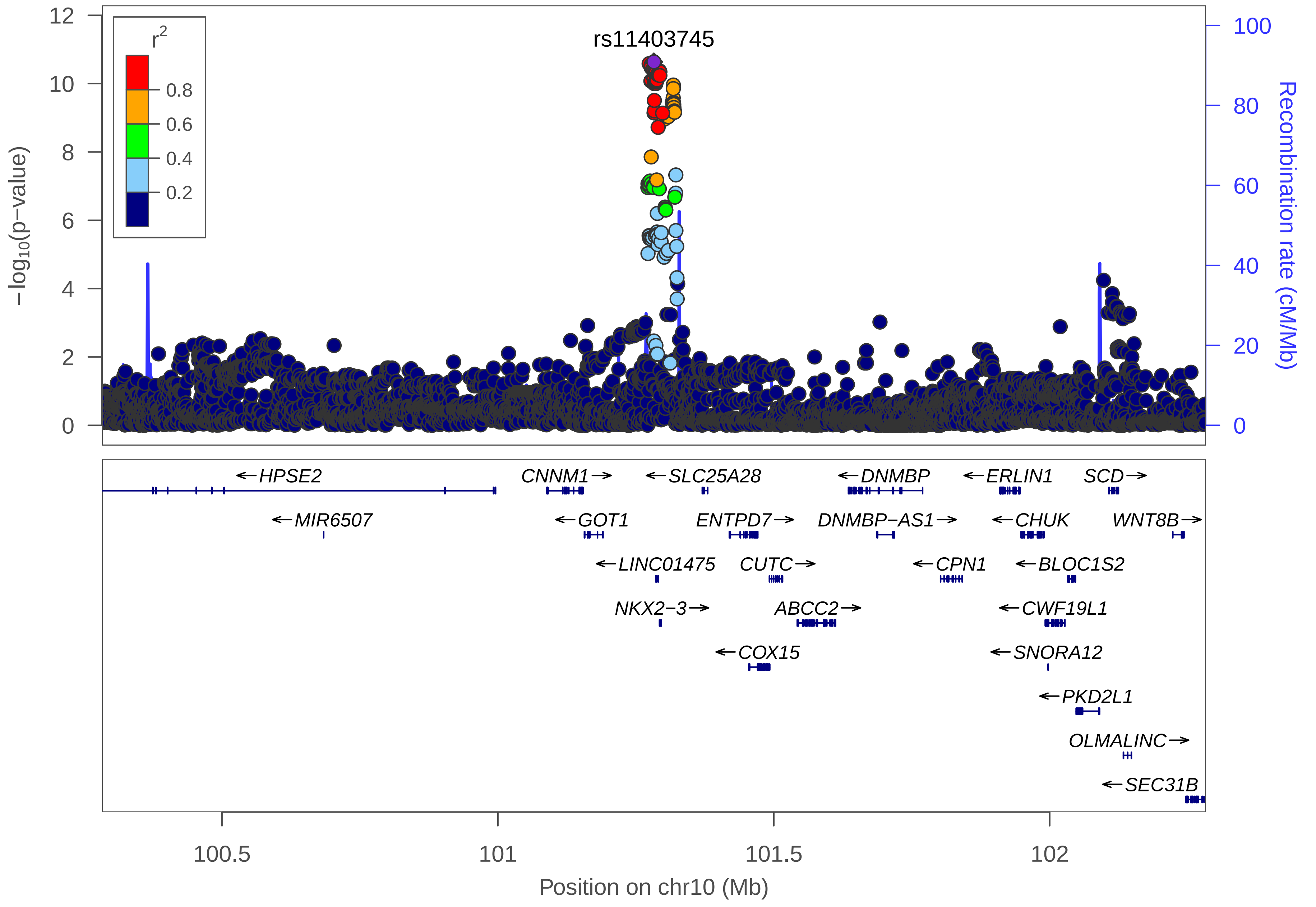

rs11403745
r2
0.8
0.6
0.4
0.2
−log10(p−value)
Recombination rate (cM/Mb)
0
2
4
6
8
10
12
0
20
40
60
80
100
HPSE2
MIR6507
CNNM1
GOT1
LINC01475
NKX2−3
SLC25A28
ENTPD7
CUTC
ABCC2
COX15
DNMBP
DNMBP−AS1
CPN1
ERLIN1
CHUK
BLOC1S2
CWF19L1
SNORA12
PKD2L1
SCD
WNT8B
OLMALINC
SEC31B
100.5
101
101.5
102
Position on chr10 (Mb)

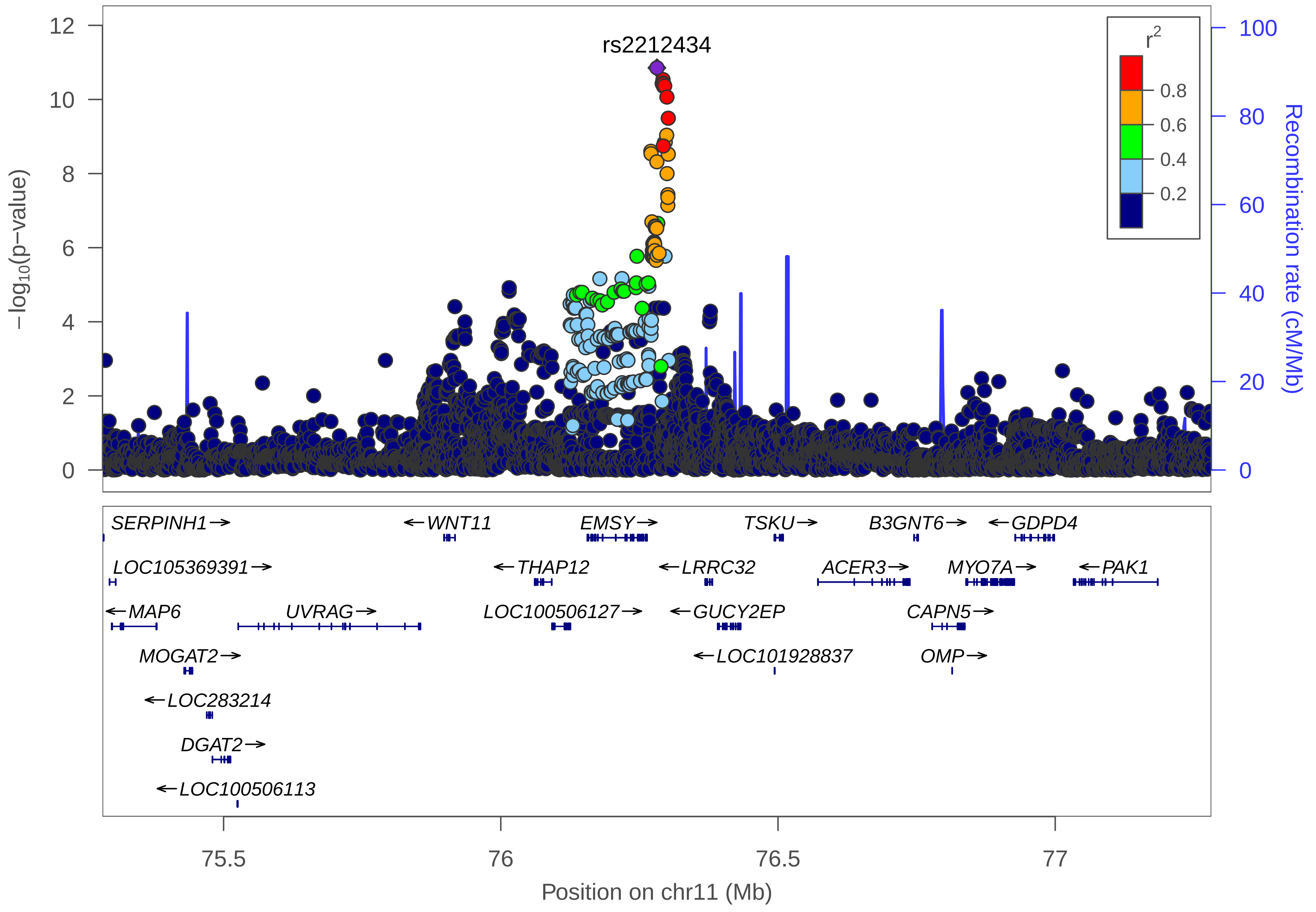

rs2212434
r2
0.8
0.6
0.4
0.2
−log10(p−value)
Recombination rate (cM/Mb)
12
10
8
6
4
2
0
100
80
60
40
20
0
SERPINH1→
←WNT11
EMSY→
TSKU→
B3GNT6→
←GDPD4
LOC105369391→
←THAP12
←LRRC32
ACER3→
MYO7A→
←PAK1
←MAP6
UVRAG→
LOC100506127→
←GUCY2EP
CAPN5→
MOGAT2→
←LOC101928837
OMP→
←LOC283214
DGAT2→
←LOC100506113
75.5
76
76.5
77
Position on chr11 (Mb)

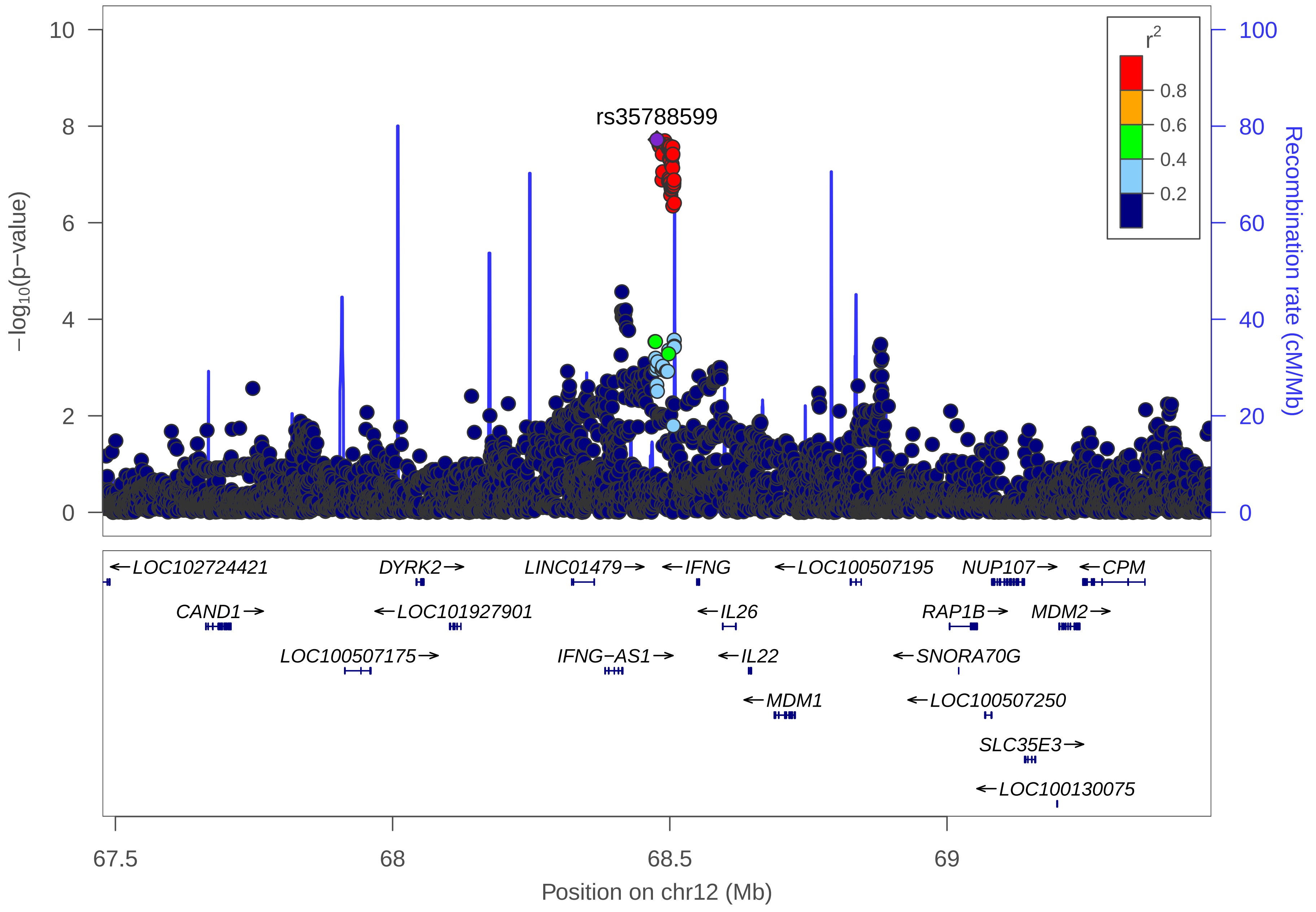
rs35788599
−log10(p−value)
Recombination rate (cM/Mb)
r2
0.8
0.6
0.4
0.2
0
2
4
6
8
10
0
20
40
60
80
100
←LOC102724421
DYRK2→
LINC01479→
←IFNG
←LOC100507195
NUP107→
←CPM
CAND1→
←LOC101927901
←IL26
RAP1B→
MDM2→
LOC100507175→
IFNG−AS1→
←IL22
←SNORA70G
←MDM1
←LOC100507250
SLC35E3→
←LOC100130075
67.5
68
68.5
69
Position on chr12 (Mb)

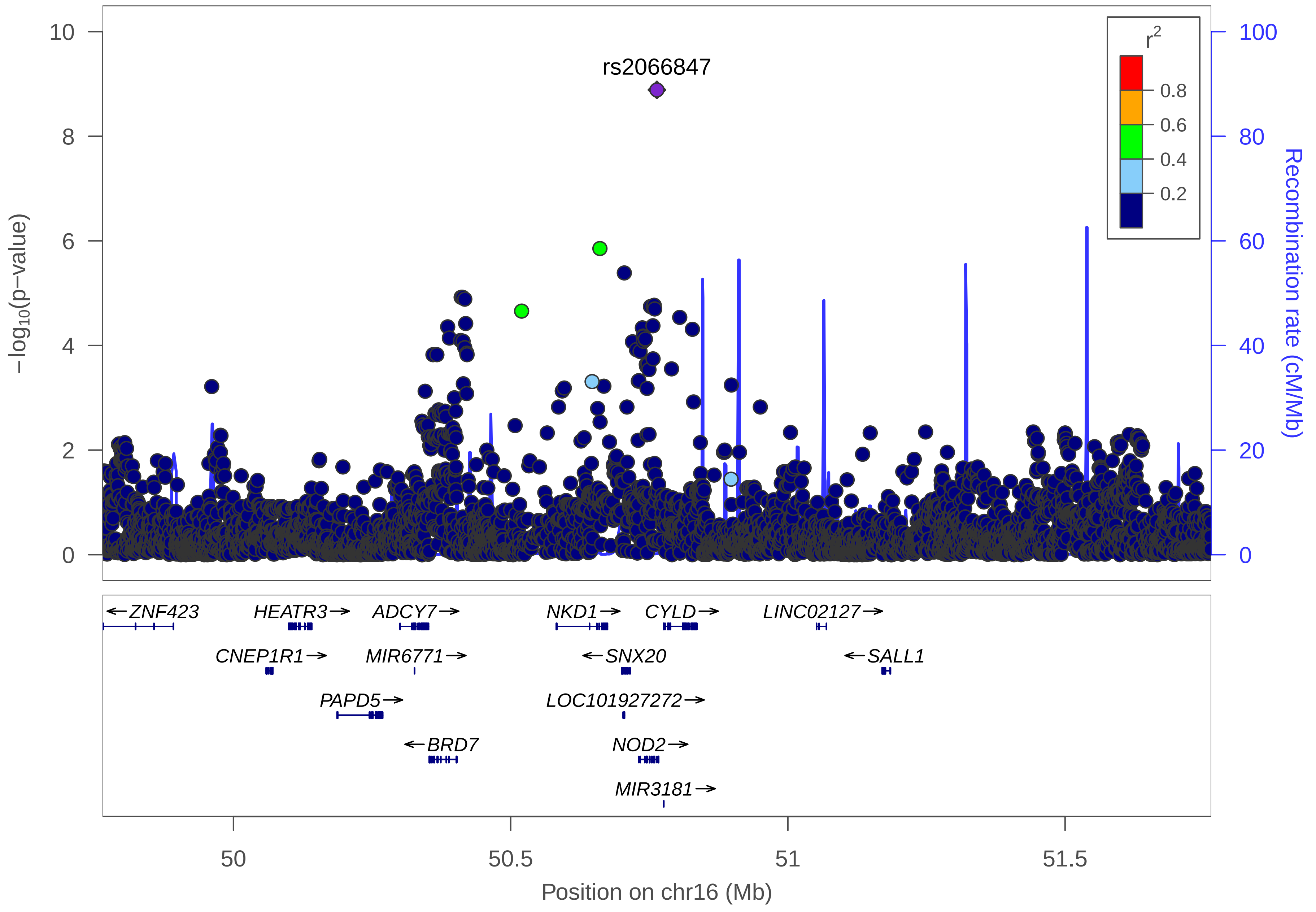

rs2066847
r2
0.8
0.6
0.4
0.2
−log10(p−value)
Recombination rate (cM/Mb)
10
8
6
4
2
0
100
80
60
40
20
0
←ZNF423
HEATR3→
ADCY7→
NKD1→
CYLD→
LINC02127→
CNEP1R1→
MIR6771→
←SNX20
←SALL1
PAPD5→
LOC101927272→
←BRD7
NOD2→
MIR3181→
50
50.5
51
51.5
Position on chr16 (Mb)

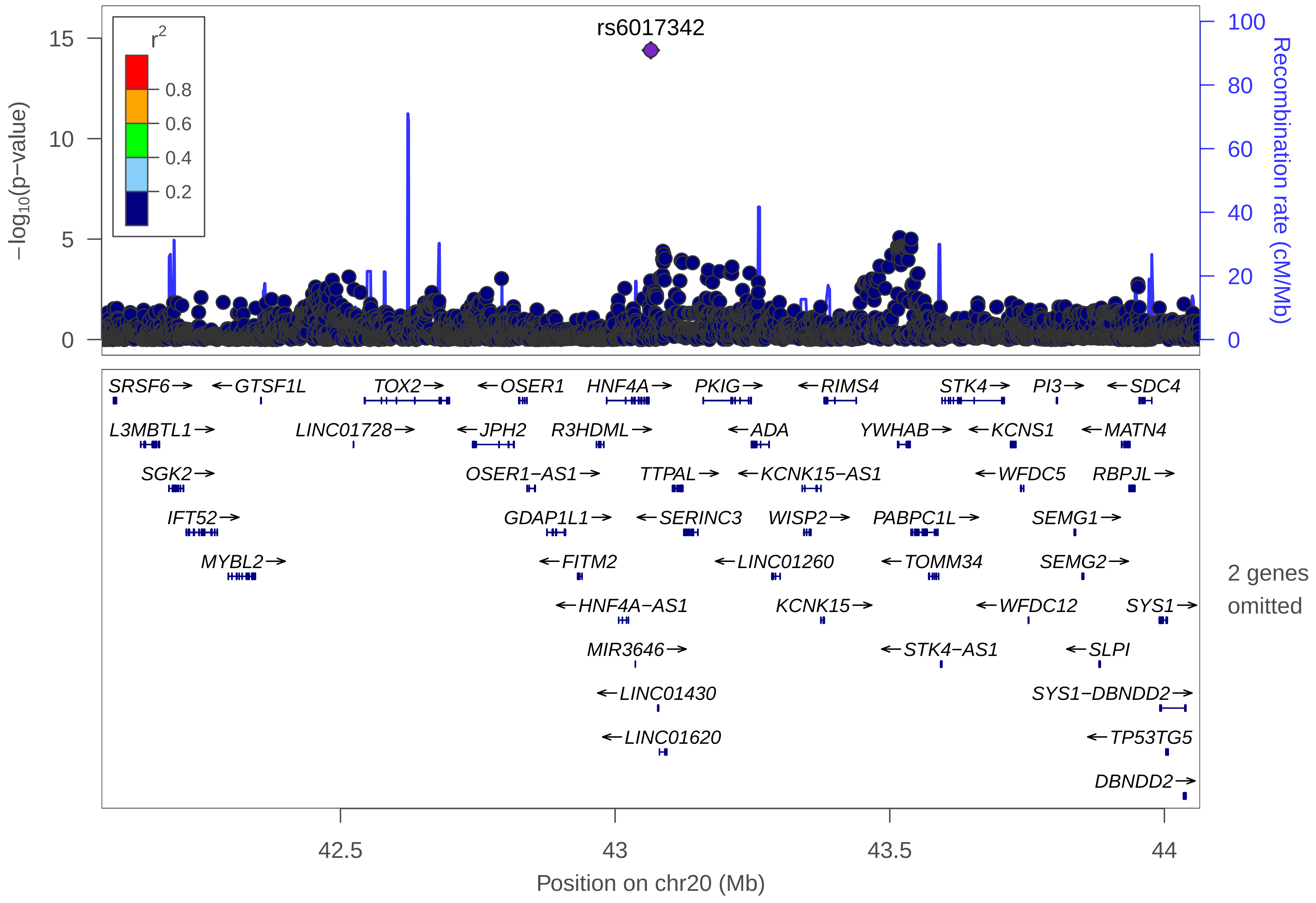
rs6017342
r2
0.8
0.6
0.4
0.2
−log10(p−value)
Recombination rate (cM/Mb)
SRSF6
GTSF1L
TOX2
OSER1
HNF4A
PKIG
RIMS4
STK4
PI3
SDC4
L3MBTL1
LINC01728
JPH2
R3HDML
ADA
YWHAB
KCNS1
MATN4
SGK2
OSER1−AS1
TTPAL
KCNK15−AS1
WFDC5
RBPJL
IFT52
GDAP1L1
SERINC3
WISP2
PABPC1L
SEMG1
MYBL2
FITM2
LINC01260
TOMM34
SEMG2
HNF4A−AS1
KCNK15
WFDC12
SYS1
MIR3646
STK4−AS1
SLPI
LINC01430
SYS1−DBNDD2
LINC01620
TP53TG5
DBNDD2
2 genes omitted
42.5
43
43.5
44
Position on chr20 (Mb)

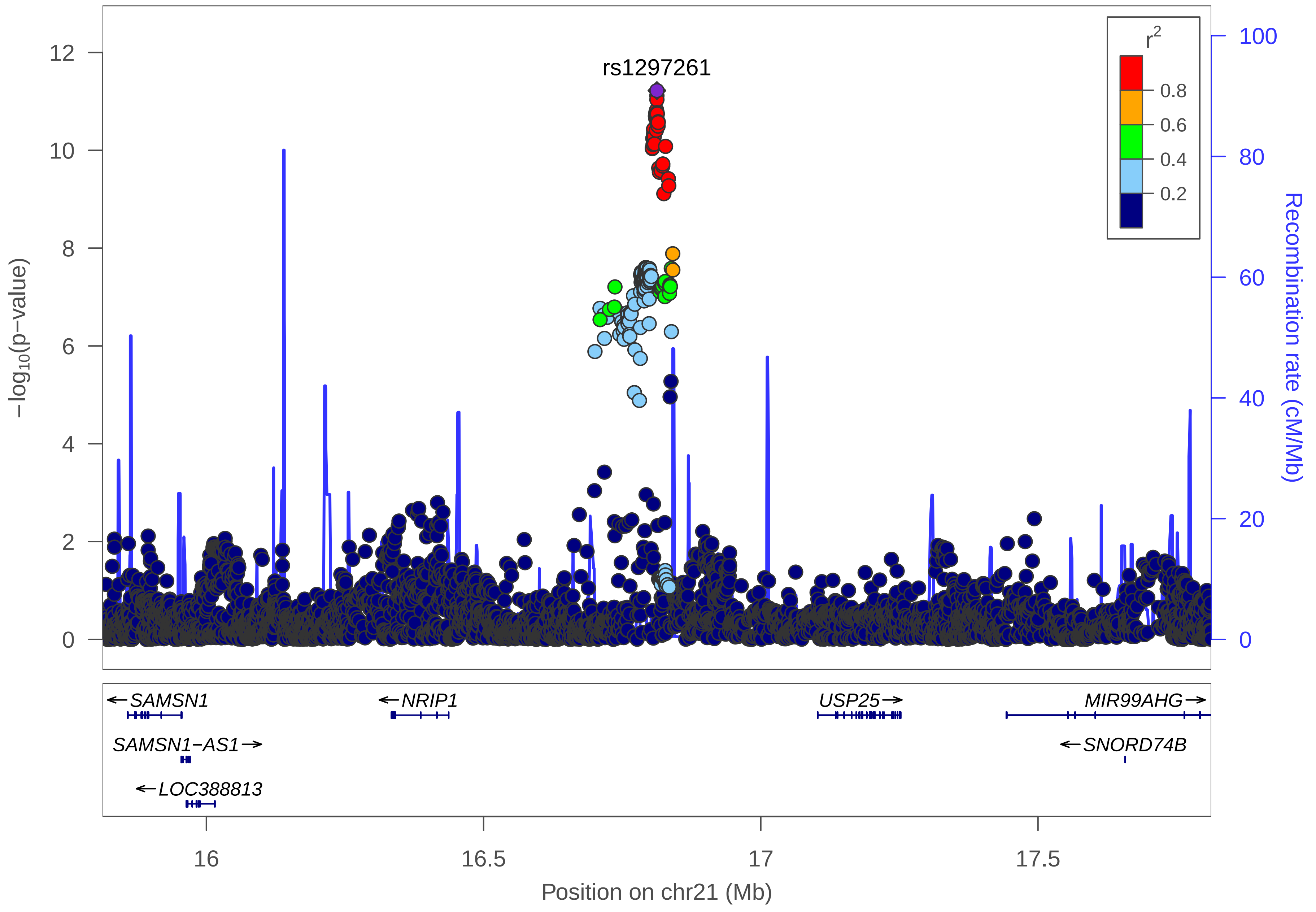

rs1297261
r2
0.8
0.6
0.4
0.2
−log10(p−value)
Recombination rate (cM/Mb)
12
10
8
6
4
2
0
100
80
60
40
20
0
SAMSN1
NRIP1
USP25
MIR99AHG
SAMSN1−AS1
SNORD74B
LOC388813
16
16.5
17
17.5
Position on chr21 (Mb)

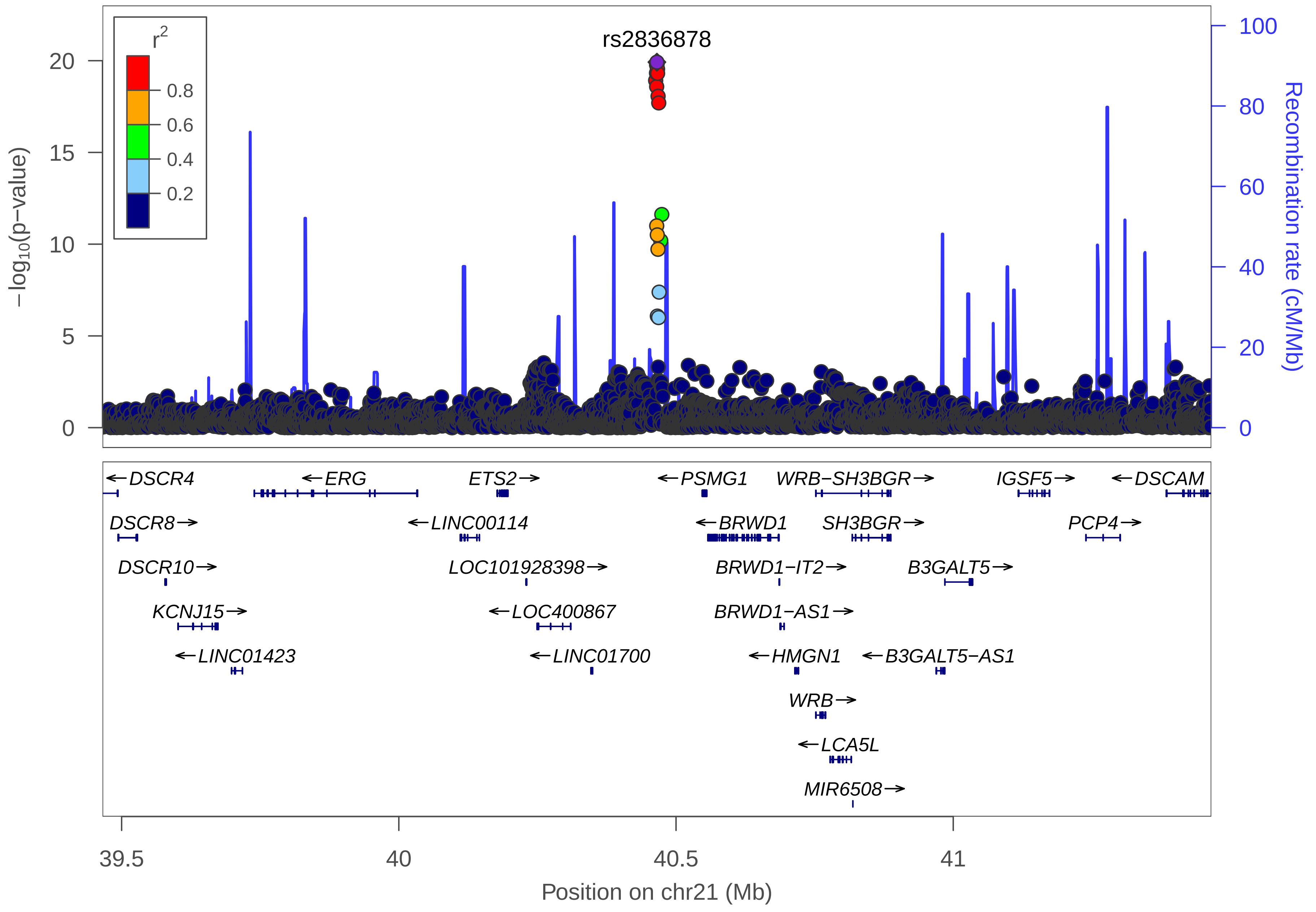

rs2836878
r2
0.8
0.6
0.4
0.2
−log10(p−value)
20
15
10
5
0
Recombination rate (cM/Mb)
100
80
60
40
20
0
←DSCR4
←ERG
ETS2→
←PSMG1
WRB−SH3BGR→
IGSF5→
←DSCAM
DSCR8→
←LINC00114
←BRWD1
SH3BGR→
PCP4→
DSCR10→
LOC101928398→
BRWD1−IT2→
B3GALT5→
KCNJ15→
←LOC400867
BRWD1−AS1→
←LINC01423
←LINC01700
←HMGN1
←B3GALT5−AS1
WRB→
←LCA5L
MIR6508→
39.5
40
40.5
41
Position on chr21 (Mb)
